# Single-cell, multi-region profiling of the macaque brain across the lifespan

**DOI:** 10.1101/2025.10.31.685880

**Authors:** Wei Yang, Kelsi L. Watkins, Alex R. DeCasien, Mary B. O’Neill, Martin O. Bohlen, Diana R. O’Day, Madeleine Duran, Chengxiang Qiu, Anastasia Meleshko, Anh Vo, Melia Menke, Diego Calderon, Cayo Biobank Research Unit, Jérôme Sallet, James P. Higham, Melween I. Martínez, Cole Trapnell, Lea M. Starita, Michael J. Montague, Michael L. Platt, Kenneth L. Chiou, Jay Shendure, Noah Snyder-Mackler

**Affiliations:** Department of Genome Sciences, University of Washington, Seattle, WA, USA; Center for Evolution and Medicine, Arizona State University, Tempe, AZ, USA; School of Life Sciences, Arizona State University, Tempe, AZ, USA; Computational and Evolutionary Neurogenomics Unit, National Institute on Aging, Bethesda, MD, USA; Brotman Baty Institute for Precision Medicine, University of Washington, Seattle, WA, USA; Department of Biology, University of Alabama at Birmingham, AL, USA; Department of Neuroscience, University of Pennsylvania, Philadelphia, PA, USA; Department of Psychology, University of Pennsylvania, Philadelphia, PA, USA; Department of Marketing, University of Pennsylvania, Philadelphia, PA, USA; Howard Hughes Medical Institute, Seattle, WA, USA; Seattle Hub for Synthetic Biology, Seattle WA, USA; Stem Cell and Brain Research Institute, Université Lyon, Lyon, France; Caribbean Primate Research Center, University of Puerto Rico, San Juan, PR, USA; Department of Bioengineering and Therapeutic Sciences, University of California San Francisco, San Francisco, CA, USA; Department of Biomedical Engineering, Duke University, Durham, NC, USA; School of Human Evolution and Social Change, Arizona State University, Tempe, AZ, USA; ASU-Banner Neurodegenerative Disease Research Center, Arizona State University, Tempe, AZ, USA; Allen Discovery Center for Cell Lineage Tracing, Seattle, WA, USA; Department of Anthropology, New York University, New York, NY, USA; New York Consortium in Evolutionary Primatology, New York, NY, USA; Department of Experimental Psychology, University of Oxford, Oxford, UK

**Author notes:** These authors contributed equally to this manuscript (Wei Yang, Kelsi L. Watkins).

## Abstract

Brain aging is a complex process with profound health and societal consequences. However, the molecular and cellular pathways that govern its temporal progression–along with any cell type-, region-, and sex-specific heterogeneity in such progression–remain poorly defined. Here, we present a transcriptomic atlas of 5.3 million cells from 582 samples spanning 11 brain regions of 55 rhesus macaques (29 female, 26 male), aged 5 months (early life) to 21 years (late adulthood). We annotate 12 major cell classes and 225 subclusters, including region-specific subtypes of excitatory and inhibitory neurons, astrocytes, and ependymal cells. We identify a vulnerable excitatory neuron population in the superficial cortical lamina and a cortical interneuron population that are less abundant later in life, along with subtle, region-specific, age-associated compositional differences in subpopulations of microglia and oligodendrocytes, whose detection required single-cell resolution. Finally, we chart convergent and divergent age-associated molecular signatures across brain regions and cell classes–where some of these signatures are sex-specific and could underlie sex biases in neurological disorders. We find that age-associated transcriptional programs not only overlap substantially with those seen in Alzheimer’s disease (AD), but also unfold along distinct temporal trajectories across brain regions, suggesting that aging and AD may share molecular roots that emerge at different life stages and in region-specific, sex-specific windows of vulnerability. This work provides a temporal, regional, and sex-stratified atlas of the aging primate brain, offering insights into cell type-specific vulnerabilities and regional heterogeneity with translational human relevance.

## Introduction

Aging is the strongest risk factor for both neurodegenerative disease and cognitive decline (*1*). The molecular and cellular changes that underlie age-associated phenotypes may begin years or even decades before the onset of clinical symptoms, making temporally-resolved profiling of healthy brains critical for understanding the normal trajectory of brain aging. Regional vulnerability has been reported in both aging and neurodegeneration (*2*, *3*), and sex differences in brain aging and disease add further complexity (*4*).

The precise relationships between chronological brain aging, molecular and cellular deterioration, and neurodegenerative phenotypes–and how these patterns may differ in males vs. females—remain unknown. Single-cell transcriptomics allows us to identify age-associated molecular patterns with cell-type resolution, offering a potentially powerful lens into the biological processes that may precede, accompany, or drive functional decline. Unfortunately, age-resolved, regionally diverse, sex-balanced molecular data from healthy human brains are very challenging to obtain, and thus remain uncommon. Even where such data is available, it is usually confounded by variable postmortem intervals, tissue harvest protocols, and disease comorbidities.

The rhesus macaque (*Macaca mulatta*), a nonhuman primate closely related to humans, provides an exceptional animal model with translational relevance for human brain aging (*5*). Macaques and humans exhibit extensive anatomical, circuit, and cellular brain homologies. Macaques also have a relatively long lifespan and experience naturally occurring age-related phenotypes, including sex-biased patterns, that recapitulate many aspects of human brain aging (*6*–*8*).

Here we leverage a tissue bank derived from a unique cohort of free-ranging rhesus macaques living under naturalistic conditions with minimal human interference (*9*) to construct a regionally resolved single-cell gene expression atlas of brain aging across the lifespan. We delineated cell population shifts to identify age-vulnerable cell classes and map the age-related trajectories of molecular and cellular processes across cell classes and brain regions in both males and females to identify cell-, region-, and sex-specific patterns. Finally, we integrate these findings with transcriptional analyses of Alzheimer’s disease in humans to infer evolutionary links between normal and pathological aging.

## Results

### Single nucleus, multi-region transcriptional profiling of the macaque brain across the lifespan

We performed single nucleus RNA-sequencing (snRNA-seq) on 5.3 million nuclei from 582 brain samples from 55 rhesus macaques (29 female, 26 male; aged 5 months to 21 years) derived from a free-ranging, provisioned island population living on Cayo Santiago (*10*). From each animal, we dissected up to 11 distinct regions (48-55 individuals profiled per brain region; mean 10.6 brain regions profiled per individual): anterior cingulate cortex (ACC), dorsolateral prefrontal cortex (dlPFC), inferior posterior parietal cortex (IPP), primary motor cortex (M1), entorhinal cortex (EC), hippocampus (HIP), nucleus accumbens (NAc), caudate nucleus (CN), mediodorsal thalamic nucleus (mdTN), midbrain (MB), and lateral cerebellum (lCb) (**Fig. 1A**, upper; **Fig. S1A**). We specifically targeted individuals to uniformly cover the lifespan of healthy wild macaques (*5*, *11*) and balanced representation of males and females (**Fig. 1A**, lower).

**Figure 1.**
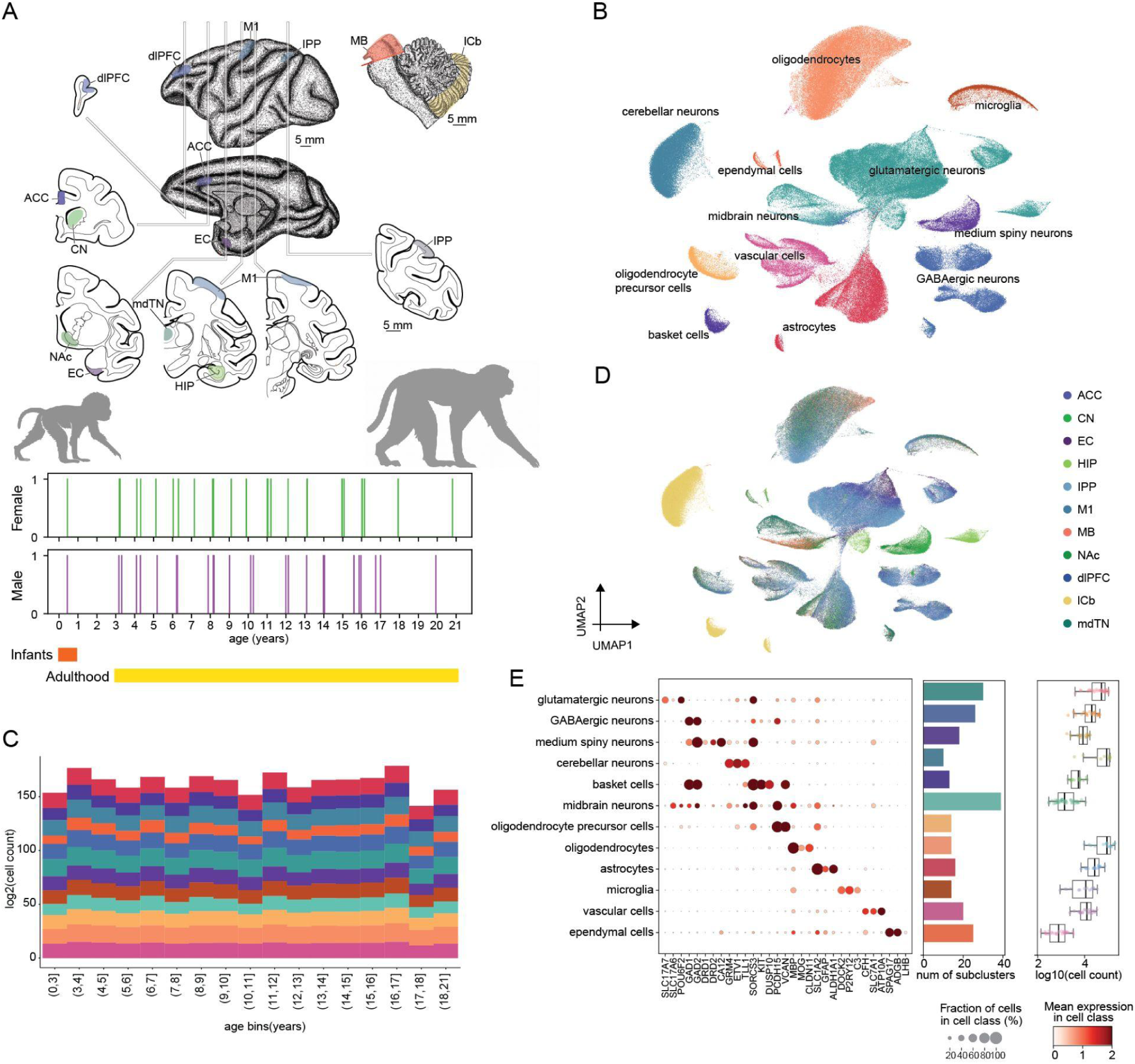
Single-cell, multi-region profiling of the macaque brain across the lifespan. **A)** Schematic of 11 brain regions profiled in this study (top), together with the age and sex distribution of the 55 individual macaques (bottom). **B)** UMAP projection of randomly downsampled snRNA-seq dataset (∼10% of all cells) colored by cell class assignment. **C)** Barplot showing log_2_(cell count) of each cell class across the 11 profiled regions in each age bin. **D)** UMAP projection of the same downsampled set of cells shown in panel B, here colored by anatomical region-of-origin. **E)** Marker gene expression dotplot for each cell class (left). Barplot showing number of subclusters from unsupervised Louvain clustering of each cell class (middle). Boxplot showing log_10_(cell count) for subclusters associated with each cell class (right).

Using an optimized sci-RNA-seq3 protocol (*12*), we recovered a median of 5,726 nuclei per sample, and a median of 1,918 transcripts (measured by unique molecular identifiers, UMIs) per nucleus, a sixfold gain in molecular depth over our previously published macaque brain atlas (*13*) (**Fig. S1B**; **Table S1**). Cerebellar "sentinel" samples were included in each experimental batch to monitor for technical variation, which was minimal (**Fig. S1C**).

Unsupervised Louvain clustering, informed by marker gene expression and label transfer from existing primate, human, and mouse brain atlases (*13*–*16*), facilitated annotation of 12 major cell classes spanning neuronal and non-neuronal lineages. Neuronal classes included glutamatergic neurons, GABAergic neurons, midbrain neurons, medium spiny neurons (MSNs), cerebellar neurons (including projecting granule cells and purkinje cells), basket cells (cerebellar basket). Non-neuronal classes included astrocytes, oligodendrocytes, oligodendrocyte precursors, microglia, vascular cells and ependymal cells. lineages (**Fig. 1B,E**; **Supplementary Note 1**). These classes were robustly detected across individuals, regions and age groups (**Fig. 1C-D**; **Fig. S1D**). Cell class distributions reflected expected anatomical specializations, with a few exceptions likely attributable to minor dissection errors (**Table S2**; **Supplementary Note 1**).

As a preliminary analysis directed at understanding how age shapes transcriptomic variance, we generated pseudobulk transcriptomes for each region, age group, and individual. Principal component analysis (PCA) revealed that inter-regional variation dominates global expression structure, while age- and sex-related variance is more apparent within individual regions (**Fig. S1E**; **Supplementary Note 2**). Upon performing variance partitioning for individual regions, age explained a modest fraction of pseudobulk transcriptomic variance (median 1.1–2.6% across regions), with the cerebellum showing the strongest age-association (2.6%) (**Fig. S2A**). Sex explained 1.5-2.2% of variance of only 15-23% of genes (**Fig. S2B**; **Table S3**), consistent with analyses of bulk RNA-seq data from our own studies and the human GTEx project (*8*, *17*). We also noticed substantial transcriptomic differences between infants and other age groups, so we decided to perform subsequent linear tests on animals >3 years old to focus on transcriptional differences across the adult lifespan rather than those occurring between development and adulthood.

### Unsupervised identification and annotation of 225 neuronal and non-neuronal subclusters

To identify more fine-grained cell subtypes, we performed unsupervised subclustering separately within each of the 12 major cell classes, resulting in 225 transcriptionally distinct subclusters spanning neuronal and non-neuronal lineages (**Fig. 1E**; **Fig. S2C**; **Fig. S3**; **Table S4**). These subclusters reflected expected ontogenic relationships and were annotated using both comparison to reference datasets and manual curation (**Supplementary Note 3**). Throughout the remainder of the manuscript, we refer to clusters derived from unsupervised subclustering of our data as ‘subclusters’. These subclusters are annotated to neuronal and glial subtypes defined by external datasets, which we refer to as ‘subtypes’. For each subtype, we have identified one or more subclusters mapping to it, suggesting that our dataset might provide finer resolution than existing subtype resolution.

For example, glutamatergic neurons were sorted into subclusters that were either broadly distributed across regions or exhibited strong region-specificity, specifically in the hippocampus, entorhinal cortex, and thalamus. The hippocampal subclusters comprised neurons from the well-organized CA1 and CA3 subfields, as well as the dentate gyrus (DG) region. The entorhinal cortex subclusters from L2/3 and L5 clustered separately from neocortical subclusters for the same layers, a similar organization as observed in the human brain (*18*). Subclusters in the thalamus exhibited unique molecular signatures (*NXPH1+* and *RNF220+*) that were absent in the neocortex, as observed in both mice and humans (*18*) (**Fig. 2A**). Subclustering of other cell classes captured both canonical and rare populations, such as *LAMP5+* interneurons in cortex (**Fig. 2B**), dopaminergic neurons in midbrain (**Table S4**), and specialized oligodendrocyte, astrocyte and immune subtypes (**Fig. S4A-E**). We were able to confidently annotate 85% of these subclusters using the following complementary approaches: (i) computationally comparing our subclusters to subtypes described in reference human, mouse, and macaque datasets (*13*–*16*, *19*), and (ii) manually cross-referencing subcluster marker genes to known subtype-specific marker genes (*15*, *16*, *18*–*24*). Most subclusters showed strong concordance with known marker genes and external references from human, mouse, and macaque brains (**Fig. 2**; **Fig. S5**; **Table S4**). This high-resolution, multi-region dataset and corresponding annotations establish a foundation for investigating how age and sex shape the heterogeneity of the primate brain at single-cell resolution.

**Figure 2.**
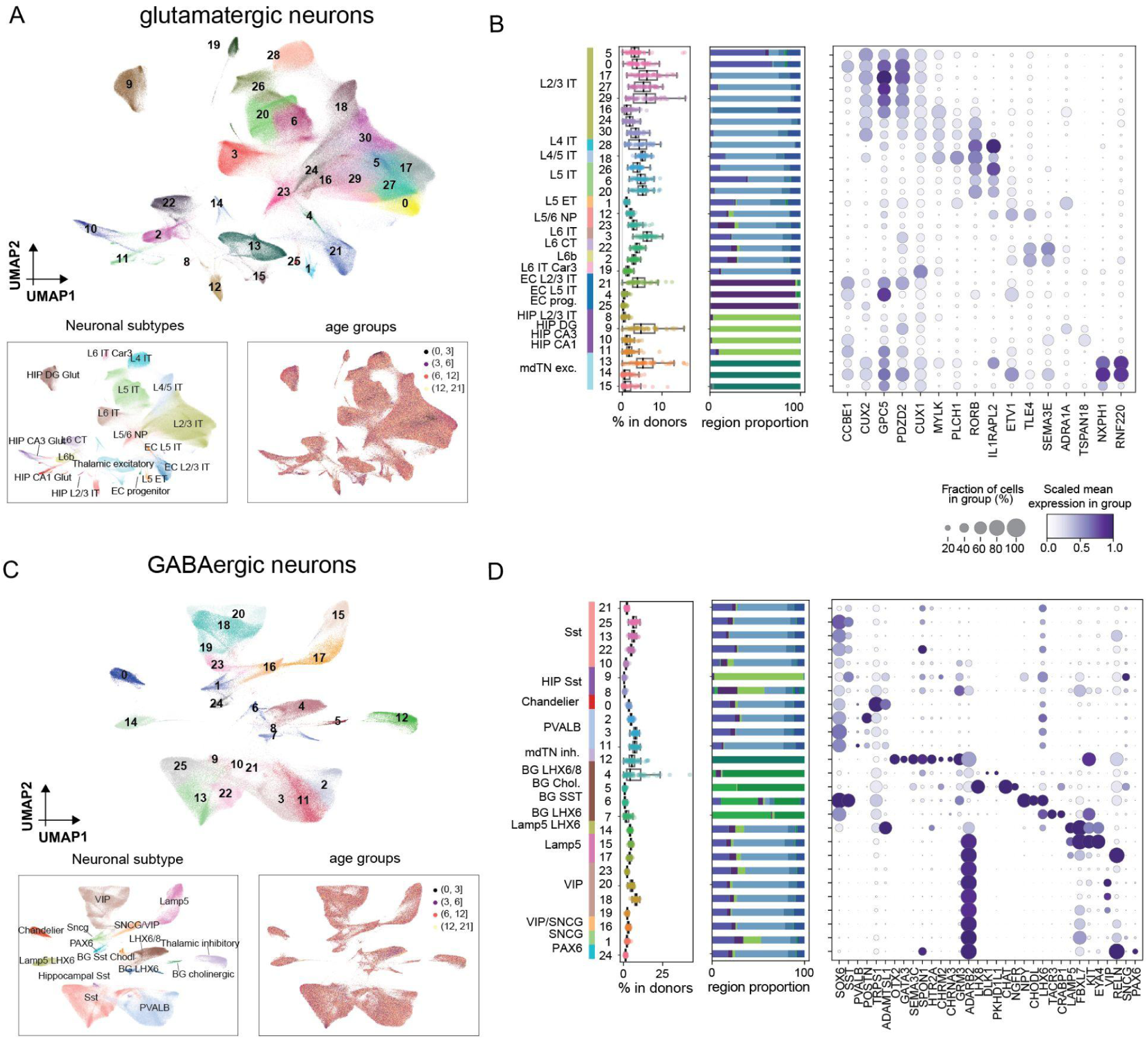
Heterogeneity and regional specificity of glutamatergic and GABAergic neuronal classes. **A)** UMAP visualization of glutamatergic neuronal subclusters identified by unsupervised Louvain clustering (top). Insets show the same UMAP colored by annotations based on label transfer from neuronal subtypes defined by external literature (*14*) (bottom left) or by age group (bottom right). Subclusters are well-stratified by subtype labels, but not by age group. **B)** Each row corresponds to a glutamatergic neuronal subcluster. Boxplot showing proportion of each subcluster out of all glutamatergic neurons across individual donors, without stratification by region (left). Subclusters exhibit relatively consistent proportions across individuals. Barplots showing the regional distribution of each subcluster (middle). Color assignments are identical to Fig. 1D. Many glutamatergic subclusters exhibit regional specificity. Dotplot of scaled mean expression of marker genes of glutamatergic subtypes from a list of reference datasets (*15*, *16*, *18*–*24*) in our subclusters (right). **C-D**) Same as panels A-B but for GABAergic neurons.

### Age-associated dynamics of cell class and cell subcluster composition

To investigate the temporal dynamics of cell-class composition in these 11 brain regions over the course of the primate lifespan, we performed differential abundance analysis for each region. As differences between infants/juveniles and adults are much larger than those among adults, we excluded individuals younger than 3 years old from this analysis (**Fig. 3A, S1E**). Also, rather than using linear models of proportions, which ignore variation in the number of cells present in each sample, we applied a Poisson log-normal model directly on the cell counts as implemented in *Hooke* (*25*).

**Figure 3.**
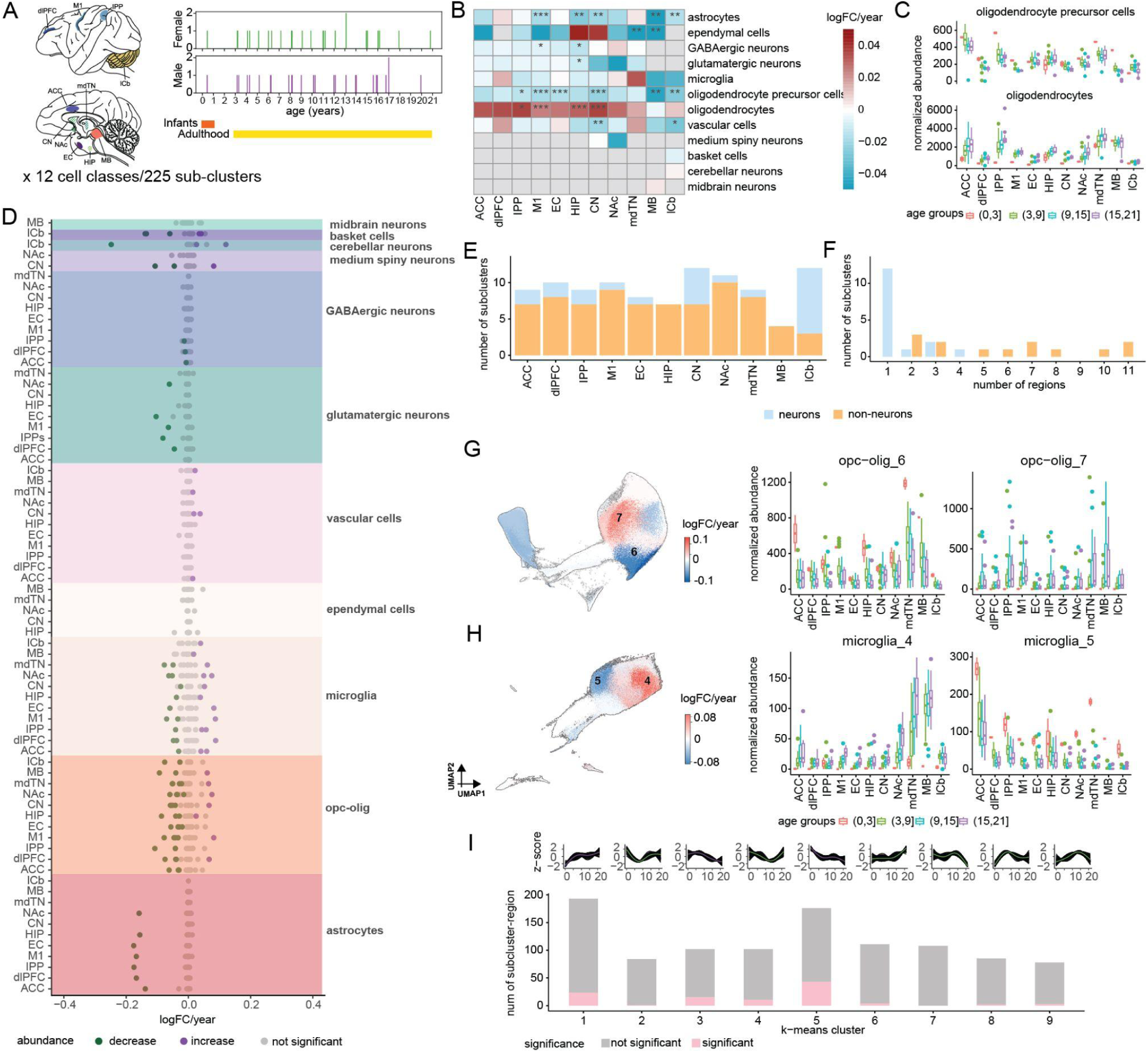
Age-associated dynamics of cell class and cell subcluster composition. **A)** Schematic of differential abundance test (3-21 years) applied to 12 cell classes and 225 subclusters in 11 brain regions. **B)** Heatmap showing log-fold change of abundance change each year (coefficient output from the linear model) for each cell class-by-region combination. LFSR <0.001: ***, 0.01: **, 0.05:*. **C)** Boxplot showing normalized cell abundance (0-21 years) in each individual after transformation by the poisson log-normal model in *Hooke* (*25*) for each age group across regions. **D)** Dotplot showing log-fold change of abundance each year (coefficient output from the linear model) for each cell subcluster and region combination. Differences are colored by decrease in abundance (green), increase in abundance (purple), and change not statistically significant (grey). **E)** Barplot showing number of age-associated subclusters in each region colored by neuronal (orange) and non-neuronal (blue) subclusters. **F)** Barplots showing the number of regions with significant differences in abundance were detected for each subcluster. **G)** UMAP showing abundance change of oligodendrocyte and OPC subclusters in log-fold change each year. Subclusters with brain-wide differences were labeled and their cell abundance were shown on the boxplots to the right (left). Boxplot showing normalized cell abundance (0-21 years) in each individual after transformation by the poisson log-normal model in *Hooke* (*25*) for each age group across regions for oligodendrocyte and OPC subclusters with brain-wide differences (right). **H)** Same as panel G for microglial subclusters. **I)** Non-linear trajectory (0-21 years) of cell abundance differences for each subcluster and region combination (top). Barplot summarizing the number of subclusters in each trajectory (bottom), with age-associated subclusters identified *Hooke* (*25*) labeled with pink color.

We modeled cell counts in each of the 12 cell classes across 11 brain regions (85 combinations of cell class and region; 48-51 animals per region) (**Table S6**). Both GABAergic and glutamatergic neurons were significantly less abundant in the hippocampus of older animals (**Fig. 3B**), plausibly reflecting age-related atrophy linked to cognitive decline (*26*). Additionally, the relative abundance of oligodendrocytes increased with age in most regions except the midbrain, while the relative abundance of oligodendrocyte precursor cells decreased with age across the brain (**Fig. 3B-C**), consistent with prior studies (*8*, *27*, *28*).

### Non-neuronal subclusters exhibit greater sensitivity to aging than neuronal subclusters

To detect finer age-associated shifts in cellular composition that may be masked at the cell class level, we quantified differential abundance of subclusters relative to their cell class within each brain region of adult macaques (age > 3 years). Focusing on subclusters with >100 cells to ensure statistical power (**Table S5**), we assessed 1,039 subcluster-region combinations (range: 57–113 subclusters per region). To further boost sensitivity, especially in regions with lower cell counts, we applied multivariate adaptive shrinkage (MASH), which leverages shared patterns across regions (*29*, *30*). For subcluster-regions showing age-related abundance differences in adult animals, we further characterized their non-linear trajectories throughout the lifespan (0-21 years). This granular analysis revealed that age significantly patterned the composition of specific cell subclusters in adult macaques (**Fig. 3A,D**; **Fig. S6A, Table S7**). Altogether, 101 of 1,039 subcluster-region combinations (10%) showed significant age-associated differences in abundance (local false sign rate, LFSR < 0.05). These effects varied by region and cell class. The lateral cerebellum had the highest proportion of age-associated subclusters (n = 12, 21%), while the midbrain had the fewest (n = 4, 4%). Among cell classes, basket cell subclusters were most affected by age (39% of its subcluster-region combinations were age-associated), whereas midbrain neurons and ependymal cells remained largely stable across the lifespan.

Across the brain, non-neuronal subclusters exhibited more extensive age-related differences in abundance compared to neuronal subclusters. This trend was consistent across all regions examined, with the exception of the lateral cerebellum, potentially due to its lower representation of non-neuronal cells (**Fig. 3E**). Similar patterns have been observed in mouse brains, in which non-neuronal cell classes—especially glia—undergo notable shifts in cell state proportions with age, while neuronal clusters remain relatively stable, suggesting conservation of this phenomenon across mammals (*31*).

In particular, certain astrocytes, oligodendrocyte precursor cells & oligodendrocytes (“OPC-olig”), and microglia subclusters each exhibited age-associated shifts across more than five regions. (**Fig. 3F, S6A**). One homeostatic astrocyte subcluster (subcluster 1; *LYPD6B*⁺ *HRH1*⁺) was very abundant in early life but sharply declined in early adulthood (**Fig. S6B**). Oligodendrocyte precursor cells (OPC-olig subclusters 12 and 13) and differentiating oligodendrocytes (OPC-olig subclusters 6) declined broadly across the brain, consistent with a global reduction in progenitor pools with age (**Fig. 3B,G**; **Fig. S6A**). In contrast, the abundance of pre-myelinating oligodendrocytes (OPC-olig subclusters 4 and 8) remained stable, while myelinating oligodendrocytes underwent state transitions toward a subcluster characterized by upregulation of the EL4 regulon and downregulation of myelination genes (OPC-olig subcluster 7; **Fig. 3G**; **Fig. S5C**; **Fig. S6C**). This altered state closely resembles a damaged oligodendrocyte phenotype described in amyotrophic lateral sclerosis (*32*), suggesting a potentially conserved mechanism of oligodendrocyte dysfunction in aging and disease. The increase in abundance of this subcluster was most pronounced in M1 and NAc (**Table S7**).

In contrast to glial cells, neuronal subclusters were largely stable in abundance across the lifespan. However, we identified some subsets of neurons that were selectively vulnerable to age. These include glutamatergic neuron subcluster 5 (*KCNK13*⁺ *C4H6orf141*⁺), corresponding to a subcluster of neocortical layer 2/3 intratelencephalic (L2/3 IT) neurons (**Fig S6A**), which was very abundant in early life but showed dramatic reductions later in life (**Fig. S6D**). Notably, this neuronal subtype—characterized by upregulation of synaptic signaling and downregulation of genes involved in reactive oxygen species (ROS) generation and neurodegenerative diseases compared to other L2/3 IT neurons—is significantly depleted in Alzheimer’s disease (AD) (*15*, *18*) (**Fig. S6E**). We also observed age-associated depletion of inhibitory interneuron GABAergic neuron subcluster 13 (*SP110^+^ STK32A^+^*), corresponding to somatostatin (SST) cortical interneurons (IPP, dlPFC, ACC) (**Table S4**), which are also impacted by AD (*15*).

### Microglial subclusters reveal age-associated transitions and mirror human APOEε4 aging signatures

Aging in macaques is associated with substantial shifts in microglial subcluster abundance. Migratory microglia (subcluster 5, *ENPEP^+^ SEMA6A^+^*) declined with age, while neuroinflammation-associated microglia (subcluster 4, *GLDN^+^ RASAL2^+^*) increased with age. These shifts occurred across nearly all brain regions, with the cerebellum as the sole exception (**Fig. 3H**; **Fig. S6A**). Microglia subcluster 4 is enriched for proinflammatory cytokine response genes (*CCL3, IL1B, CD83, SPP1, LPL, ITGAX*) typically upregulated in response to amyloid β pathology (*33*) (**Fig. S6F**), suggesting convergence between aging-related inflammation and inflammatory responses in AD. We observed regional variation in age-associated abundance differences, with microglia subcluster 4 in dlPFC, M1, and EC showing the largest increases, while border-associated macrophages (microglia subcluster 7) were regionally restricted but displayed significant age-related increases in the ACC and NAc (**Table S7**; **Fig. S6A**).

To investigate the regulatory landscape of age-associated microglial state transitions, we grouped microglia subclusters into two groups. Group 1 (subclusters 3-6, 8, 10, 13) spans a trajectory from surveilling to redox-active and, ultimately, proinflammatory states, while Group 2 (subclusters 0, 1, 9, 12) comprises more stable surveilling microglia (**Fig. S5D**; **Fig. S7A–B**). Pseudotime analysis of Group 1 identified an age-correlated trajectory (**Fig S7C-D**) and a continuum of gene expression differences (**Fig. S7A, E**). Earlier-activated genes (pseudotime gene cluster 1) were enriched for axon guidance pathways, consistent with microglial roles in tract development (*34*). Later-activated genes (pseudotime gene clusters 2, 4) were enriched for genes in lipid and fatty acid metabolism (*e.g. CPM*, *SPP1*, *GPNMB*, *ACSL1*, *APOE*, which were upregulated toward the end of the trajectory) (**Fig. S7E–F**). These metabolic pathways have been linked to disease-associated microglial states and are known to be modulated by genetic variation (*35*).

### Continuity between development and brain aging

Our densely sampled time-course dataset enabled us to capture diverse, non-linear trajectories of cell subcluster abundance across the lifespan. To model these patterns, we applied a cubic spline fit to the abundance of each subcluster-region combination across lifespan (0-21 years), and clustered the resulting trajectories into nine groups (**Fig. 3I**). We found that many subclusters showed consistent, monotonic patterns across lifespan. In particular, group 1 and 5 subclusters exhibited sharp shifts in early life (0–5 years) followed by more gradual changes throughout adulthood (5–21 years). These continuous trajectories suggest that brain aging reflects the extended activity of developmental programs rather than a functionally distinct process, consistent with recent work highlighting the molecular continuity between development and aging (*36*).

### Age substantially patterns gene regulation throughout the brain

Exploratory analyses of pseudobulked gene expression recapitulated findings by us and others, based on bulk RNA-seq (*8*), that regional heterogeneity in brain gene expression exceeds age-related variation (**Fig S1E**). However, it does not follow that age is unimportant—rather, it may simply be that strong regional differences in cellular composition and gene expression mask age-related effects that are present at a cell-class- and/or region-specific level (**Fig. S1E**). To test this idea, we examined differences among cell-class- and region-specific age-related transcriptional programs by identifying age-related differentially expressed genes (aDEG). Specifically, we modeled gene expression in a negative binomial linear mixed model framework, including age, sex, and sequencing depth as fixed effects, and donor identity as a random effect, in adult macaques (age > 3 years). For statistical power, we set a heuristic threshold that removed subcluster-region combinations with fewer than 300 cells or 30 animals (**Table S5**). We then applied multivariate adaptive shrinkage (MASH) (*25*, *29*) to all subcluster-region combinations in a parent cell class, which refined aDEG effect size estimates and facilitated robust detection of shared aDEGs across subcluster-region combinations (**Methods**).

Age was associated with substantial differences in gene regulation across the brain: 62% of the genes we tested (9,372/15,117) were significantly differentially expressed as a function of age (LFSR < 0.05) in at least one subcluster-region combination (**Fig. 4A**). Across subcluster-region combinations, we found that upregulated genes outnumbered downregulated genes, with an average of 58% of aDEGs being upregulated (one-sample t-test p-value < 2.2e-16) (**Fig. S8A**). When examining individual brain regions, we observed regional variation in the percentage of upregulated aDEGs. Subclusters from the hippocampus (HIP) and ACC were significantly less likely to exceed the 58% average, while subclusters from the EC and MB were significantly more likely to exceed the 58% average (two-sided binomial test p-value < 0.025).

**Figure 4.**
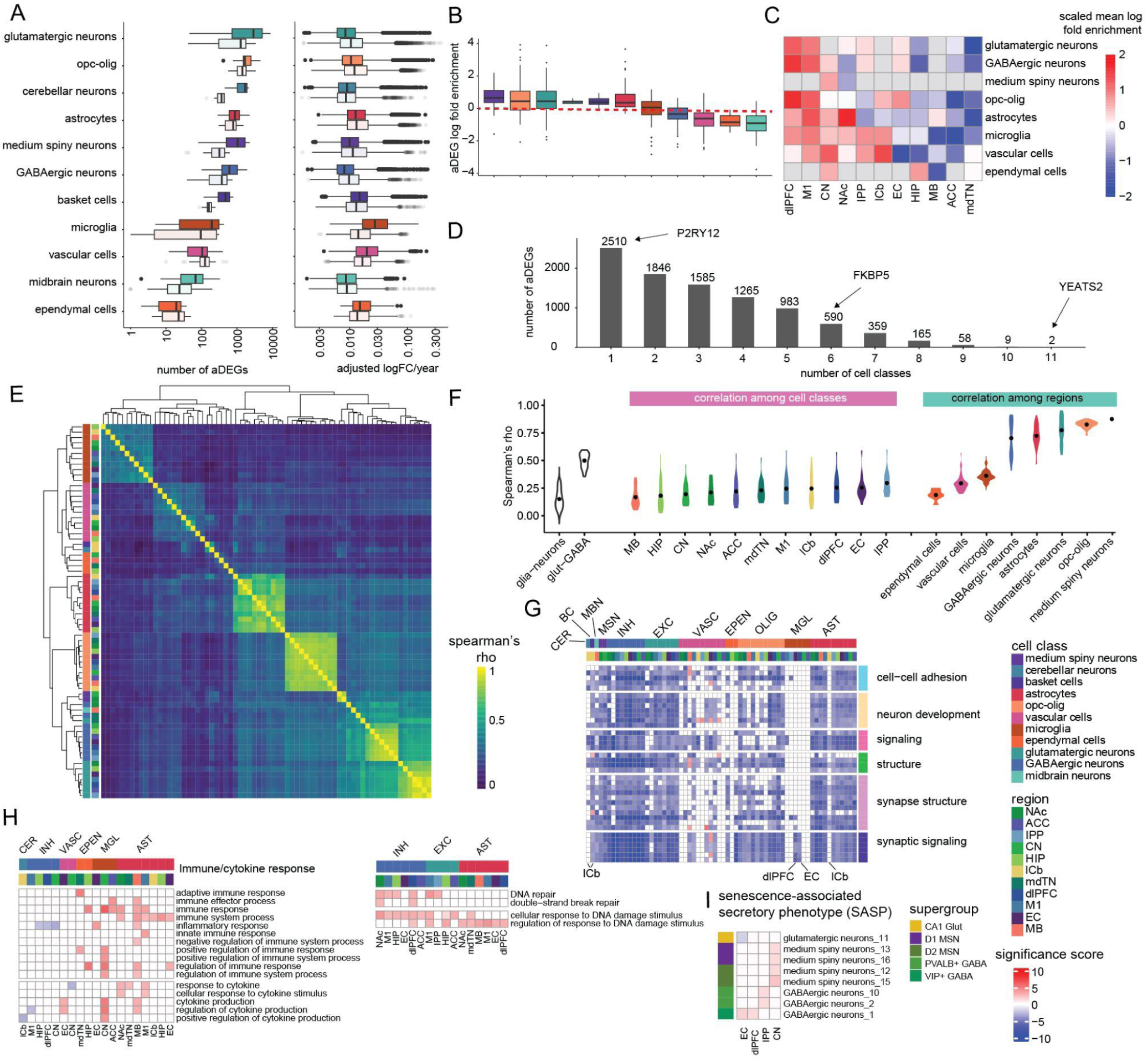
Cell type-specific effects of age on the brain transcriptome. **A)** Boxplots showing the number of aDEGs (left) with significant (LFSR<0.05) association with age and the adjusted log-fold change per year for these aDEGs (right) in each combination of subcluster-*x*-region. Light color indicates upregulated aDEGs and dark color indicates downregulated aDEGs. **B)** Boxplot showing the log-fold enrichment of proportion of aDEG (number of aDEG/number of genes tested) over average proportions across all combinations of subcluster-*x*-region after correcting number of aDEGs by cluster size and sequencing depth with a loess model. The dotted line corresponds to no enrichment. **C)** Heatmap showing the average log-fold enrichment of subcluster-*x*-regions in each cell type. Each row is scaled by z-transformation to identify regions with higher average residuals. **D)** Barplot showing the number of aDEGs unique to a single cell class or shared among multiple (2–11) cell classes. Examples of broadly shared aDEGs (*e.g. YEATS2, FKBP5*) and specific aDEGs (*e.g. P2RY12*) are highlighted. **E)** Spearman’s correlation of age coefficients of all genes from the linear mixed model after MASH correction among all combinations of cell class-*x*-region combinations. **F)** Violin plot illustrating Spearman’s *rho* in panel E across four categories of comparisons: (i) glial vs. neuronal classes, (ii) glutamatergic vs. GABAergic neurons, (iii) different cell classes within the same brain region; and (iv) different brain regions within the same cell class. **G)** Heatmap showing GSEA results of GO pathways broadly shared among combinations of cell types and regions. Significance score is calculated by -NES sign x log10(p-value). AST: astrocytes; MGL: microglia; OLIG: olig-OPC; EPEN: ependymal cells; VASC: vascular cells; EXC: glutamatergic neurons; INH: GABAergic neurons; MSN: medium spiny neurons; CER: cerebellar neurons; MBN: midbrain neurons; BC: basket cells. **H)** Heatmaps showing GSEA results of immune response and DNA damage/repair pathways which show cell type and region specific enrichments. **I)** Heatmap showing specific neuronal subclusters in certain regions with enrichment in senescence-associated secretory phenotype (SASP).

Our ability to detect aDEGs in each cell class was influenced by several technical factors, including: (i) sequencing depth, (ii) the number of cells in a given subcluster-region combination, and (iii) the number of genes tested (**Fig. S8B**). After correcting for these confounding factors, we found that MSNs showed the highest enrichment of aDEGs on average, followed by OPC-olig cells (**Fig. 4B**), while glutamatergic neurons and astrocytes showed comparable log fold enrichment in specific regions (**Fig. S8C**). Within cell classes shared among regions, dlPFC, M1 and CN consistently showed more aDEGs than average (z-score > 0), suggesting they might be hotspots of brain aging (**Fig. 4C**; **Fig. S8C**). These three regions are involved in cognition (dlPFC, CN) and motor control (M1, CN) (*37*–*39*), potentially linking transcriptional changes to the cognitive and mobility declines that accompany aging. aDEG effect sizes, which should be unbiased by the number of cells per test, were similarly distributed across most cell classes, with slightly higher effect sizes in microglia, vascular and ependymal cells, hinting at some consistent and programmed age-associated differences in expression across genes in distinct cells and regions (**Fig. 4A**). Additionally, the effect sizes of age up-regulated and age down-regulated genes were similar in most cell classes, except in microglia which had stronger effects in upregulated aDEG (**Fig. 4A**).

Most aDEGs were restricted to only one or a few cell classes (27% of aDEGs were cell-class-specific), consistent with aging predominantly driving cell-class-specific transcriptional changes (**Fig. 4D**). Seventy-five percent of these cell-class-specific aDEGs were detectably expressed in ≥7 cell classes, suggesting such age-dependent changes are due to cell-class-specific regulation rather than a lack of expression in other cell classes (**Fig. S8D**). Altogether, we identified 2510 cell-class-specific aDEGs, ranging from 3%-18% of aDEGs detected in each cell class (**Fig. 4D**). For example, *P2RY12*, a gene important for microglial homeostasis, is expressed in 9 cell classes, but declines with age only in microglia. This expression pattern mirrors that seen in disease-associated and aged microglia (*40*) (**Fig. S8E**), further supporting the relevance of cell-class-specific aDEGs to functional decline in brain aging.

Only a handful of aDEGs were shared broadly across multiple cell classes. *FKBP5*, a regulator of glucocorticoid receptor sensitivity whose increase has been linked to depression (*41*), schizophrenia (*42*) and AD pathology (*43*), showed brain-wide upregulation in 4 neuronal cell classes, consistent with ours and others’ findings (*6*, *8*, *44*). In addition, we detected age effects in *FKBP5* in non-neuronal cells, including upregulation in OPC-olig cells (brain-wide) and downregulation in vascular cells (in lCb and M1) (**Fig. S8F**). This suggests there might be divergent age effects on key age-associated genes dependent on the cell class. Notably, two aDEGs were identified across all cell classes, suggesting a shared mechanism across cell classes (**Fig. 4D**). One of these was *YEATS2*, which increases in expression with age in all neuronal cell classes and most glia (except for microglia where expression is lower in older age). Notably, *YEATS2* is a paralog of the *Drosophila* ENL/AF9 domain-containing protein that can regulate lifespan in flies, and, when knocked down in neurons, extends lifespan (*45*). As a selective histone crotonylation reader, we speculate that *YEATS2* plays a key role in age-associated epigenetic remodeling (**Fig. S8G**).

### Cellular and regional heterogeneity are linked to aging vulnerability

To take further advantage of the regional and cellular breadth of our dataset, we asked whether some cell classes or regions exhibit more similar age effects on gene expression than others. In particular, towards assessing whether aging is governed by global vs. local regulatory programs, we sought to identify shared and distinct signatures of the aging transcriptome that span cell classes and anatomical structures.

To do so, we assessed the similarity of age effects on all detectably expressed genes across cell classes and regions. This allowed us to evaluate how broadly age-related transcriptional differences are shared, or diverge, across the brain’s cellular and spatial landscape. We found that age effects on gene expression were primarily structured by cell identity: age effects were more similar within the same cell class across regions than between different cell classes within the same region (**Fig. 4E-F**). This suggests that aging impacts the transcriptome predominantly through cell-class-intrinsic programs. Among neurons, excitatory and inhibitory classes show particularly strong age-associated transcriptional similarity with one another across regions. Neurons also share age-associated patterns with OPC-olig cells and astrocytes, suggesting partially coordinated aging patterns among electrically active or metabolically coupled cell classes. In contrast, other glial populations, such as microglia, vascular cells, and ependymal cells, exhibit highly distinct and specialized age patterns, consistent with their divergent functional roles in homeostasis, immunity, and barrier integrity.

Within cell classes, regional context variably shapes age effects on the transcriptome. OPC-olig cells display consistent age signatures across neuroanatomical regions, pointing to a systemic aging trajectory linked to myelin maintenance and progenitor pool exhaustion (**Fig. 3B**; **Fig. S6E**). Neurons, in contrast, show strong regional specificity. Glutamatergic and GABAergic neurons in particular share similar age signatures within cortical regions (**Fig. S9A-B**), while astrocytes demonstrate distinct age transcriptomes that are contingent upon region—potentially reflecting regional activity demands and metabolic constraints (**Fig. S9C**). For example, Bergmann glia—a cerebellum-specific astrocyte subtype—exhibits a unique suite of cerebellum-restricted aDEGs that is shared with basket cell and cerebellar neurons (**Fig. S9D-E**).

In contrast, cell-class-specific and regionally restricted aDEGs may highlight specialized aging vulnerabilities. Cortical neurons display unique aDEGs enriched for synaptic signaling and oxidative stress pathways, which are not observed in their counterparts outside the cortex (**Fig. S9F**). These cortex-specific aDEGs show limited overlap between excitatory and inhibitory neurons (Jaccard index = 0.04), yet may contribute to regional susceptibility. Similarly, microglia exhibit both global and localized age signatures. While their aDEG profiles are largely cell-class specific, some genes had anatomically restricted age effects with functional implications. For instance, microglia in the entorhinal cortex exhibit age-associated upregulation of RhoGTPase (*ARHGAP27*) involved in clathrin-mediated endocytosis—a pathway implicated in amyloid-β clearance and neuroinflammation—potentially linking regional microglial aging to mechanisms of neurodegeneration (**Fig. S9G**) (*46*).

To understand the biological functions associated with these pathways, we performed gene set enrichment analysis (GSEA) on aDEGs for each cell subcluster-region combination. We identified both broadly conserved and regionally specialized pathways, which cluster by cell class and reflect both shared and distinct mechanisms of age-related variation (**Fig. S10A**). Among the most consistently downregulated pathways across neuronal populations, astrocytes, and OPC-olig cells were those involved in synaptic signaling. This synaptic decline is also present in rodents (*47*, *48*), non-human primates (*49*, *50*) and humans (*51*–*54*), consistent with a strong evolutionary conservation that may underlie reduced plasticity and connectivity in the aging brain (**Fig. 4G**). Other broadly downregulated programs include those related to neuronal development, axon structure, cell adhesion, and cell signaling–differences consistent with known reductions in dendritic complexity and glutamate receptor activity during aging (*55*).

However, region and cell class context modulate these trends. In microglia, age-associated downregulation of synaptic signaling is restricted to the EC and dlPFC, while in the cerebellum, it is restricted to basket cells. Even among glutamatergic neurons, synaptic membrane-related genes decline more rapidly in neocortical regions compared to the hippocampus or EC (**Fig. S11**). This observation is consistent with stereological studies showing pronounced synapse loss in prefrontal cortex but relatively preserved synapses in hippocampus during normal aging (*56*, *57*). Furthermore, this pattern in healthy brains contrasts with AD, where early loss occurs in EC and hippocampus (*58*). In vascular and ependymal cells, the downregulation of genes involved in cell junctions suggests potential impairment of the blood-brain barrier with age (**Fig. 4G**) (*59*), further reflecting tissue-specific vulnerability to functional decline.

We observed cell-class-specific enrichment of brain aging hallmarks (*60*–*62*). For example, astrocytes show age-associated upregulation of genes involved in the DNA damage response, while glutamatergic neurons and GABAergic neurons upregulated genes in both DNA damage and repair (**Fig. S10B**). Astrocytes exhibit a unique signature of cell senescence which is not enriched in other cell classes (**Fig. S10B**), consistent with their decrease in abundance with age in multiple regions (**Fig. 3B**). Given the current literature revealing important roles of astrocytes in neuromodulation (*63*–*65*), loss of astrocytes with age might be connected to disruption of the neuronal network and synaptic functions. Upregulated aDEGs were enriched for immune activation and the inflammatory response in astrocytes, microglia, vascular cells and ependymal cells across a few regions, while GABAergic neurons and cerebellar neurons show an age-associated downregulation of the inflammatory response (**Fig. 4H**; **Fig. S10B**).

Elimination of senescent cells (“senolysis”) was recently reported to improve normal and pathological changes associated with aging in mice and monkeys (*66*–*69*). Thus, the discovery of region-specific neuronal subclusters with strong age-associated senescence markers could identify cells that might be most responsive to senolytic therapies. Genes related to senescence-associated secretory phenotype (SASP) were upregulated (p-value < 0.05, signal > 20%) in subsets of neurons in specific regions, including vasoactive intestinal peptide (*VIP+*) GABAergic neurons in EC and dlPFC, parvalbumin (*PVALB+*) GABAergic neurons in IPP, and several D1 and D2 medium spiny neuron subclusters, particularly in CN. In contrast, these genes were downregulated with age in the glutamatergic neurons of the subiculum (**Fig. 4I**).

### Age-associated genes overlap with transcriptional hallmarks of Alzheimer’s disease

After identifying age-associated transcriptional programs, we next asked whether neurodegenerative diseases like AD represent an amplification of these same age-related pathways, or instead reflect distinct, disease-specific mechanisms. To do so, we compared aDEGs from our dataset to AD-differentially expressed genes (AD-DEGs) from three independent age-matched, case-control single-cell studies (**Fig. S12A-B, 5A-B**) (*18*, *70*, *71*). Enrichment analyses revealed significant directionally concordant overlap in aDEGs with up- and down-regulated AD-DEGs (**Fig. 5C**; >55% of aDEGs overlap with AD-DEGs to the same cell class, and 11-85% of aDEGs overlap with AD-DEGs from other cell classes), indicating highly convergent transcriptional responses. Notably, microglia exhibited directionally consistent convergence with microglia-specific AD-DEGs, implicating this cell class in a common aging-neurodegeneration axis (**Fig. 5A-B, S12A-B**). Within the DEGs that comprise the overlap between aDEGs and AD-DEGs, 9% were microglia-specific (one-sided binomial p-value: 0.07; **Fig. 5D**). Microglia-specific convergence was enriched for adaptive and innate immunity, suggesting similar immune responses in both aging and neurodegeneration (**Fig. 5E**). In contrast, convergent transcriptional signatures in astrocytes, OPC-olig, and neurons reflected more global aging and AD programs across cell classes. The convergence of these cell classes are mostly (>97%) driven by pan-cellular DEGs (aDEGs shared by ≥ 2 cell classes) and are enriched for synaptic functions and neuron development (**Fig. 5F**).

**Figure 5.**
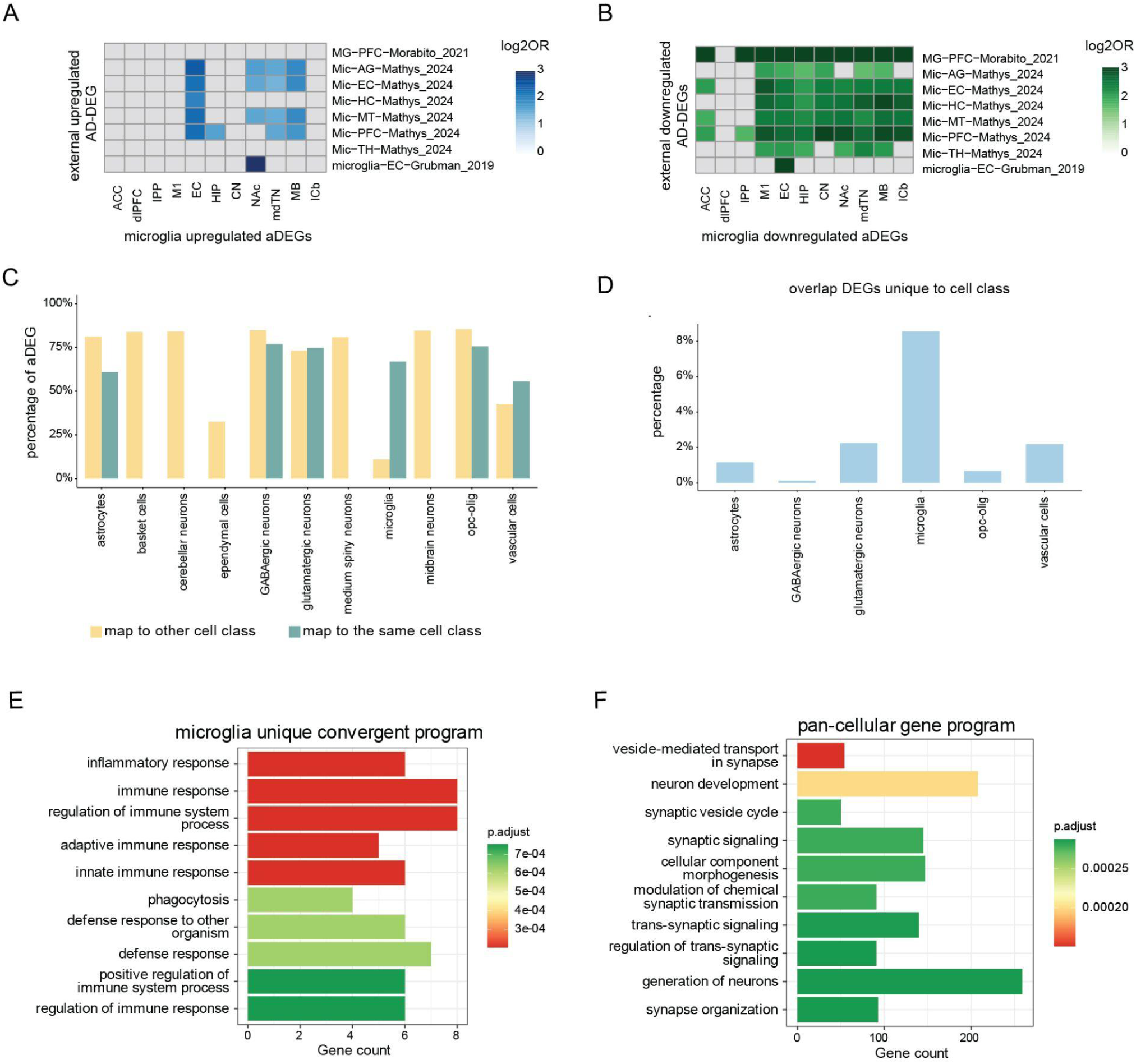
Similarities between transcriptional changes associated with aging and Alzheimer’s disease. **(A-B)** Heatmap of log_2_ odds ratio (OR) of enrichment in microglia performed by a Fisher’s exact test of overlaps between: **A)** upregulated aDEGs and AD DEGs, or **B)** downregulated aDEGs and AD DEGs (p adjusted < 0.001). **C)** Proportion of aDEGs overlapping AD DEGs with the same cell class (blue) or other cell class (yellow) within aDEGs of each cell class. **D)** Proportion of aDEGs that are uniquely overlapping AD DEGs within one cell class. **E)** GO term enrichment of genes uniquely detected in microglia and overlap between aDEGs and AD DEGs. **F)** GO term enrichment of DEGs detected in >1 cell class in our dataset or external AD datasets.

Having identified strong overlap between aDEGs and AD-DEGs, we next asked when in the lifespan these directionally concordant genes–hereafter referred to as AD-convergent genes–undergo their most pronounced expression shifts. To address this, we applied k-means clustering to the average expression of AD-convergent genes across individuals grouped into six age intervals: 0-3, 3-6, 6-9, 9-12, 12-15, and 15-21 years. These genes exhibited diverse temporal dynamics across glutamatergic neuronal subclusters, with strong regional specificity (**Fig. S13**). In cortical L2/3 IT neurons, most AD-convergent genes showed gradual, directional shifts—either increasing or decreasing steadily from early to late life (**Fig. S13A-B**). In contrast, entorhinal cortex specific L2/3 neurons exhibited more abrupt upregulation of AD-convergent genes in late adulthood (**Fig. S13C-D**). Within the hippocampus, CA1, CA3, and dentate gyrus (DG) neurons showed a distinct pattern: most upregulated genes increased in early adulthood, while downregulated genes sharply declined in late life (**Fig. S13E-J**).

### Cell-class- and region-specific sex differences in the primate brain transcriptome

Sex differences in aging are well-documented in humans and nonhuman animals (*72*, *73*), including in the brain (*74*–*76*). Both the additive and interactive effects of sex and chronological age may contribute to sex differences in vulnerability to neurodegenerative diseases and help explain inter-individual heterogeneity in aging. In line with recent findings in humans (*51*), we found no significant sex differences in brain cellular composition (*i.e.*, the relative proportions of cell classes or cell subclusters) after multiple hypothesis test correction (**Table S7**; all FDR > 0.2). We therefore sought to explore sex differences in gene expression within cell classes, as well as any overlap or interactions between sex differences and age-related differences.

Applying the same negative binomial linear mixed model framework and multivariate adaptive shrinkage (MASH) analysis described above to data from adult macaques (age > 3 years), we identified pervasive effects of sex on gene expression across regions and cell classes. More than half (57%) of all genes tested were differentially expressed (LFSR < 0.05) between males and females in at least one cell subcluster and region (8,687/15,117). Similar to aDEGs, most sex-differentially expressed genes (sDEGs) were cell-class-specific or minimally shared: 81% of sDEGs occurred in only 1 to 3 cell classes (**Fig. S14**; **Table S8**). Also similar to aDEGs, this was not driven by genes with highly restricted expression, as 75% of cell-class-specific sDEGs were detectably expressed in ≥ 9 cell classes, suggesting cell-class-specific regulatory differences even in genes that are broadly expressed across the brain.

sDEGs were not evenly distributed across cell classes or brain regions. On average, oligodendrocytes had 4.5-fold more sDEGs than expected after controlling for the number of cells and total transcripts per subcluster (**Fig. 6A**; **Methods**). The number of sDEGs for oligodendrocytes was particularly pronounced following our application of MASH (∼58-fold increase, compared to 31-fold in GABAergic neurons, the cell class with the next highest increase), suggesting an unusually high degree of sharing of sDEGs across oligodendrocyte subclusters and regions. While characterizing signatures of sharing, we found that sDEGs shared across oligodendrocytes were enriched for genes that are differentially expressed in response to androgens or estrogens (**Table S9**), suggesting that the oligodendrocyte sDEGs may be partly attributable to sex-biased steroid hormones (**Supplementary Note 4**).

**Figure 6.**
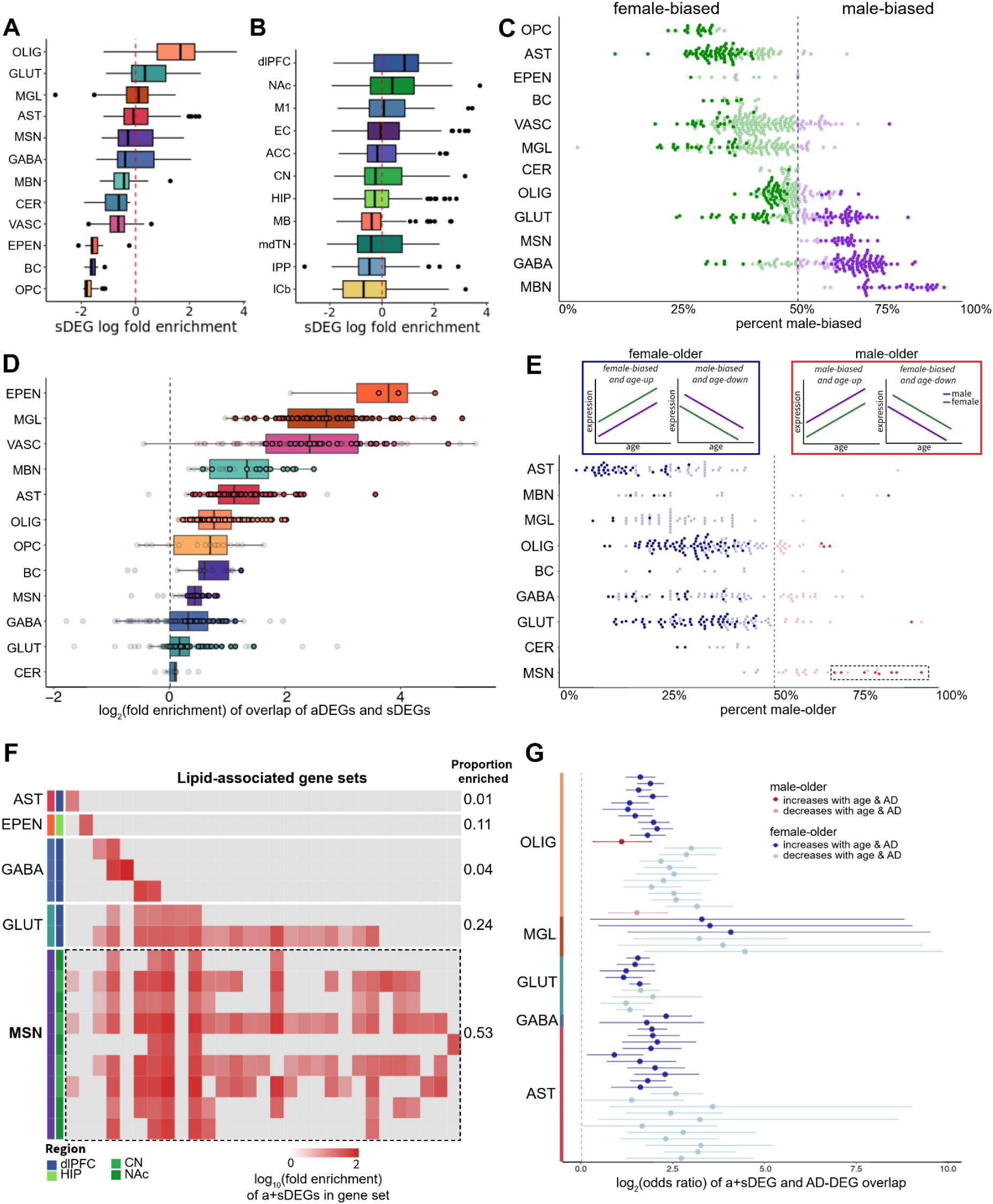
Convergence of sex and age on the brain transcriptome. **(A-B)** Enrichment of sDEGs by cell class **(A)** or cell type **(B)**, relative to expectation based on cluster size and nUMIs (Methods). **C)** sDEGs are significantly (FDR < 0.05; darker colored points) male-biased (purple) or female-biased (green), rather than balanced, in most subcluster-regions. The dotted line represents the delineation between male or female bias in sDEGs. **D)** Enrichment of a+sDEGs within subclusters by cell class. Darker colored points represent an enriched subcluster (FDR < 0.05). **E)** a+sDEGs show significant (FDR < 0.05; darker colored points) directional bias for many subcluster-region combinations. The majority of subcluster-regions show significant female-older enrichment (blue), but medium spiny neuron subclusters in the NAc and CN are exceptions, instead exhibiting male-older enrichment (red). **F)** Gene set enrichment of male-older a+sDEGs among gene sets associated with lipid metabolism. Here we are showing all subclusters that showed a significant male-older bias in their age-increasing a+sDEGs, and were enriched for at least one lipid-related gene set. Dashed box highlights the enrichment of lipid metabolism genes in medium spiny neuron male-older genes. **G)** Enrichment of AD-DEGs in a+sDEGs.

At the regional level, the dlPFC and NAc showed the strongest enrichment for sDEGs (**Fig. 6B**). Both GABAergic and glutamatergic neurons showed the strongest enrichment of sDEGs in the dlPFC (3.6- and 3.3-fold, respectively). Glutamatergic neurons also showed enrichment of sex-biased gene expression in the hippocampus, EC, M1, and ACC. These findings suggest that sex differences in excitatory neuron gene regulation are strongest in prefrontal and medial temporal regions.

We next tested whether sDEGs exhibited directional sex bias (*i.e.*, whether more sDEGs were male-biased or female-biased than expected by chance). Overall, sDEGs were evenly distributed: 47% showed higher expression in females and 53% showed higher expression in males. However, when we examined individual subcluster-region combinations, clear patterns of directional bias emerged. Of 844 subcluster-region combinations analyzed, 404 (48%) exhibited significant bias toward one sex or the other (two-sided binomial test, FDR < 0.05; **Fig. 6C**). This bias varied by cell class, *e.g.* 94% (32/34) of midbrain neuron subclusters, 83% (24/29) of medium spiny neuron subclusters, and 63% (80/127) GABAergic neuron subclusters, had significantly more male-biased than female-biased sDEGs. In contrast, 88-100% of subclusters of astrocytes, OPCs, microglia, ependymal cells, vascular cells, oligodendrocytes, and basket cells had significantly more female-biased than male-biased sDEGs. The picture was more mixed for glutamatergic neurons: 68% of subclusters (75/110) were biased, and among those, 60% male-biased and 40% female biased, while no cerebellar neuron subclusters exhibited directional sDEG bias.

Despite finding no significant sex differences in cell composition or abundance of cell subtypes, we found that cell-class-specific sDEGs are pervasive in the macaque brain. Moreover, within most cell classes, these sDEGs are directionally biased, with glia showing more female-bias in gene expression and some neuronal classes showing more male-bias. This suggests that even when subtype composition is balanced between male and female cells, the gene expression within those cells shows significant differences. These phenotypic differences may contribute to sex-specific patterns of disease vulnerability in aging even within the same subtype.

### Sex and age alter expression of similar genes across the brain

Because sex differences in gene regulation may intersect with aging to shape cell- and region-specific vulnerabilities to brain disease, we next asked whether age and sex influenced expression of the same genes. Specifically, we focused on genes showing both age- and sex-associated differential expression (a+sDEGs), which represented 30% of all tested genes (4,492/15,117). Within subcluster-region combinations containing at least 10 age DEGs or 10 sex DEGs (n = 842), we then tested for significant overlap between age and sex effects.

We observed substantial enrichment overall: 43% of all subcluster-region combinations showed significant overlap of aDEGs and sDEGs (358/842, FDR < 0.05; **Fig. 6D**). These overlaps were particularly strong for glial cell classes, including oligodendrocytes (88%, 112/127), astrocytes (72%, 68/94), and microglia (70%, 59/84). Among neurons, MSNs exhibited the highest overlap (55%, 16/29), followed by midbrain neurons (44%, 14/32), glutamatergic neurons (28%, 30/109), and GABAergic neurons (16%, 19/121). Cerebellar neurons and basket cells showed limited enrichment at 10% (1/10) and 8% (1/13), respectively. These results suggest that glial cells, particularly oligodendrocytes and astrocytes, are key sites of convergent age- and sex-related transcriptional regulation, while neuronal convergence is more variable and subtype-specific. This is consistent with findings from human studies, which have reported that glia show higher interindividual variability in gene expression than neurons (*77*, *78*)

To evaluate whether age-associated transcriptional differences more closely resemble male- or female-biased expression patterns, we examined directional relationships among a+sDEGs. Genes were classified as “male-older” when the effects of male sex and older age were aligned (*e.g.,* male-biased and upregulated with age or female-biased and downregulated with age), and “female-older” when the effects of female sex and older age aligned. We then tested enrichment of these directional patterns within each subcluster-region that showed significant aDEG and sDEG overlap. Almost half (45%) showed significant directional bias towards male-older or female-older (binomial test FDR < 0.05; **Fig. 6E**). Strikingly, 93% of these (150/162) were skewed toward female-older expression, suggesting that, at any given age, females were transcriptionally “older” than males in these cells. This bias appeared across 6 of 10 cell classes, with astrocytes, oligodendrocytes, and glutamatergic neurons each showing 60-80% of their subcluster-regions enriched for female-older a+sDEGs. In astrocytes, these female-older genes were enriched for synapse-related functions, which showed lower expression in older and female individuals. These trends suggest potential sex-specific vulnerabilities in synaptic maintenance and plasticity, processes central to cognitive aging and neurodegenerative disease risk.

MSNs were a notable exception to the broad female-older bias in a+sDEGs. MSNs exhibited significantly more male-older profiles in 7 of 16 subclusters (**Fig. 6E**). This pattern is particularly intriguing given MSNs’ role in Parkinson’s disease (PD), an age-related neurodegenerative disorder with strong male bias (*79*–*81*). These MSN male-older genes, which primarily showed higher expression in males and higher expression in older animals, were significantly enriched for lipid and cholesterol metabolism pathways (**Fig. 6F**; FDR < 0.05).

It is possible that these additive effects of sex and age on cell-specific gene regulation contribute to sex differences in vulnerability to neurodegenerative disease, particularly AD. Indeed, a+sDEGs were significantly enriched for AD-DEGs (*18*) in a cell-class-specific manner (**Fig. 6G**; **Fig. S15**). Overall, 86% of a+sDEGs tested at the cell-class-region level showed significant enrichment for AD-DEGs (60/70, FDR < 0.05). The majority (58/60, 97%) of these tests showed female-older AD-DEG overlap – due to the overall bias towards female-older transcriptional patterns across cell classes (**Fig. 6E**). Microglia, in particular, showed strong cell-class-specific enrichment for female-older a+sDEGs (**Fig. S15**). Together, this suggests that females show more AD-associated gene expression patterns than males at any given age.

### Sex modifies cell-specific age effects across the brain

Having established that age and sex each exert broad independent influences on brain gene expression, we next asked if sex modulates age-related gene expression–that is, whether age-associated transcriptional differences vary between males and females. To reduce our multiple testing burden, we focused on genes significantly associated with either age or sex (aDEGs or sDEGs; LFSR < 0.05), and modeled age-by-sex interactions in only those genes. We identified age-by-sex differentially expressed genes (a*sDEGs) in all cell classes and regions (**Fig. 7A & 7B**). In total, 1,382 unique genes (13% of 10,592 tested) exhibited significant interaction effects (LFSR < 0.05) in at least one subcluster-region, and more than half of subcluster-regions tested (60%, n = 508) contained at least one such gene. Thus, while not ubiquitous, sex-by-age interactions appear to be a pervasive feature of transcriptional aging across the primate brain. Of note, the majority of a*sDEGs (81%, n=1,117) were cell-class-specific, suggesting that sex-dependent age-related expression patterns are highly specialized to individual cell classes (**Table S8**, **Fig. S14**).

**Figure 7.**
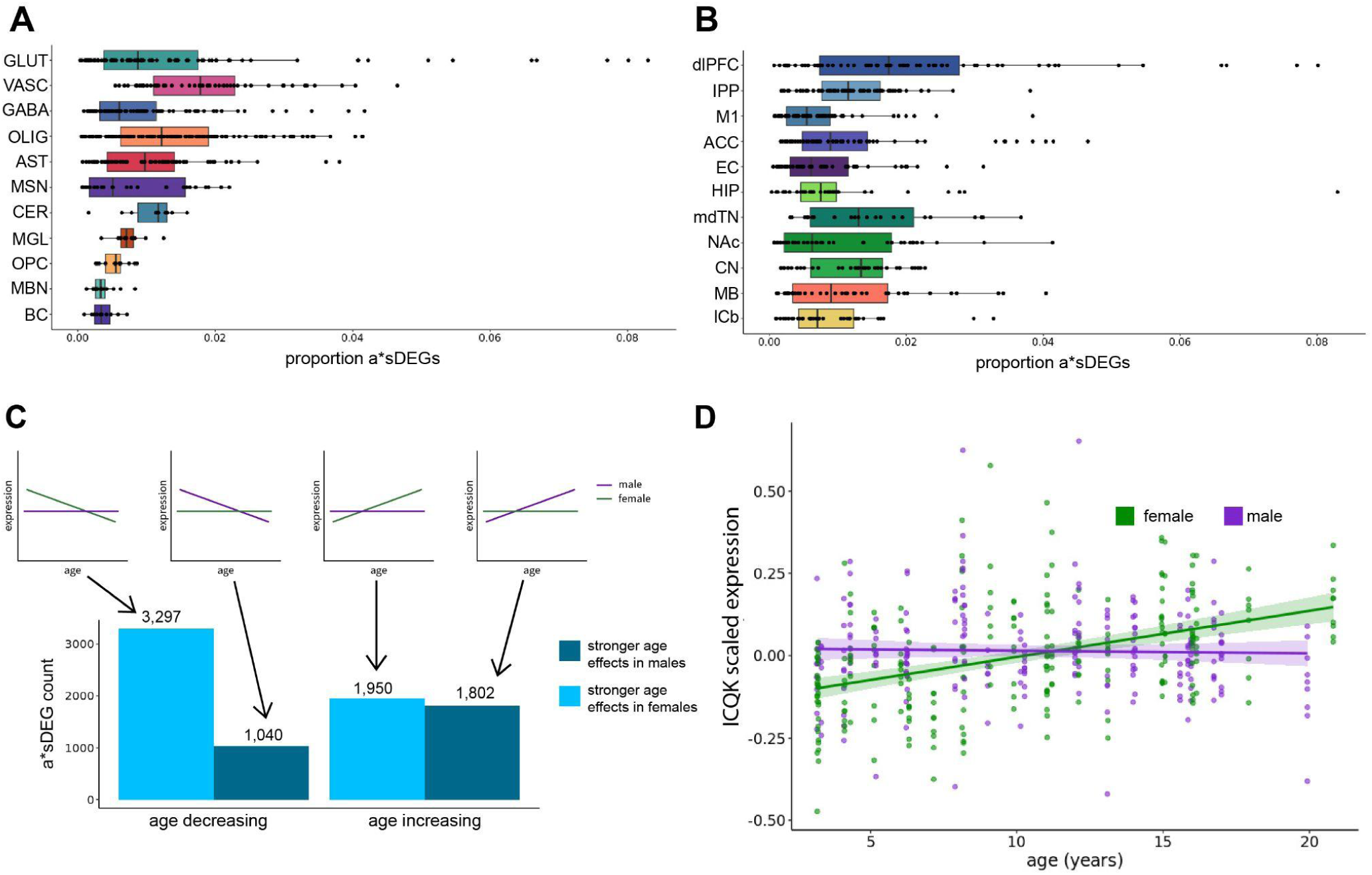
Sex-specific age effects on gene expression. **A)** Proportion of a*sDEGs out of total genes tested in each subcluster-region combination, plotted by cell class. **B)** The same as A) but plotted by brain region. **C)** A summary showing the number of a*sDEGs in each direction. **D)** Plot showing the gene expression of *IQCK* in astrocyte subclusters in which we identified an interaction of age and sex (LFSR < 0.05).

To identify cell classes most implicated in sex-specific aging, we then tested if certain cell subclusters showed an overrepresentation of a*sDEGs relative to chance. Oligodendrocytes and glutamatergic neurons showed the strongest such enrichment, with 16% of subcluster-region combinations in each of these cell classes exhibiting significant overrepresentation of a*sDEGs (20/127 for oligodendrocytes, enrichment range: 1.5- to 3.1-fold; 18/110 for glutamatergic neurons, range: 1.4- to 6.1-fold; FDR < 0.05, **Fig. 7A**). Regionally, a*sDEGs were most prevalent in the dlPFC, which contained 3-times as many enriched subclusters than any other region (27% vs. 9% in the mdTN, the next most enriched region, **Fig. 7B**).

We then assessed whether these interactions disproportionately reflected stronger age effects in males or females. This revealed a striking asymmetry: among genes downregulated with age, 3.2-fold more showed stronger effects in females (3,297) than in males (1,040). In contrast, age-associated upregulation was more balanced (1,950 stronger in females vs. 1,802 in males). Overall, 65% (5,247) of a*sDEGs exhibited stronger age effects in females, suggesting that female-biased transcriptional remodeling as a dominant axis of sex-modified aging (**Fig. 7C**).

Although highly cell-class-specific (**Fig. S14D**), some a*sDEGs showed consistent effects across multiple subclusters within the same cell class. One gene, *IQCK*, was identified as an a*sDEG in 11 astrocyte subcluster-regions, where *IQCK* expression increased more strongly with age in females than males (**Fig. 7D**). Given that *IQCK* is an Alzheimer’s risk gene whose expression correlates with amyloid and tau pathology (*18*, *82*–*84*), these female-biased age-related increases may contribute to sex differences in AD vulnerability. Together, these results highlight that sex not only additively impacts the aging process, but also interacts to potentially modify the rate and pace of aging in a cell-class- and region-specific manner.

## Discussion

Here, we generated a comprehensive single-cell atlas across 11 brain regions in both male and female rhesus macaques spanning the adult lifespan, providing a high-resolution view of cell-class- and region-specific patterns influenced by age and sex. This resource enables systematic investigation of gene regulatory programs underlying the aging process. Using this dataset, we characterized inter-regional and inter-cellular heterogeneity in aging dynamics, explored their relevance to human neurodegenerative diseases, and uncovered sex-specific differences in brain aging.

We observed substantial variability in age-related differences across brain cell classes and regions. Differences in cell abundance were more pronounced in glia and broadly observed across regions, whereas such differences in neurons were restricted to specific anatomical structures (*e.g.* neocortex, cerebellum). Our findings align with previous studies in mice, supporting the conclusion that brain aging is primarily driven by cell identity rather than regional location (*48*, *85*). Neurons show regionally stratified age-related differences (*e.g.* separation of neocortex and other brain regions). Neurons and astrocytes in the cerebellum display distinct transcriptional signatures of age, suggesting region-specific cellular activity and aging processes. Other glia, such as microglia, vascular cells and ependymal cells, also display highly distinct and specialized age relationships between regions, consistent with their spatial specialization in the brain. This inter-region and inter-cell class heterogeneity was reflected in the restriction of aging hallmarks to specific combinations of regions and cell classes. We identified astrocytes in most brain aging pathways, which have recently been implicated in neuromodulation and resilience to Alzheimer’s disease (*63*–*65*). Our findings support a role for astrocytes in age-related cognitive decline and point to several vulnerable pathways. We further observed specific neuronal subtypes displaying pathological age phenotypes (*i.e.* senescence-associated secretory phenotype). Overall, our findings highlight aging heterogeneity at the cell class, region, and subtype level. These findings include some cell types implicated in neurodegenerative diseases, thus providing potential targets for interventions to delay, prevent, or treat age-related diseases of the brain.

Our study also situates human neurodegeneration within the broader context of healthy aging in nonhuman primates. We identified age-related changes in cell populations relevant to AD, including declines in L2/3 IT glutamatergic neurons and SST+ GABAergic neurons ((*15*, *18*) and increases in neurodegenerative disease-associated oligodendrocyte and microglial subtypes (*32*, *33*). Lipid droplet-associated microglial state shifts mirrored those linked to humans with the APOEε4/ε4 genotype (*35*). While all nonhuman primates are homozygous for APOEε4 by human classification criteria (*86*, *87*), their APOE protein’s lipid-binding properties are more similar to the human APOEε3 variant. Thus, the age-linked microglial state transitions in macaque aging may capture conserved, ancestral aspects of APOEε4-related neuroinflammatory processes that precede neurodegeneration.

Across cells, we observed strong overlap between gene expression programs in normal primate aging and those implicated in human AD, implying that AD may, in part, represent an “extreme” extension of normal aging. Some transcriptional patterns emerged early in life, while others accumulated gradually, consistent with both developmental origins and lifelong progression towards AD vulnerability. Although macaques rarely exhibit the extensive amyloid and tau pathology characteristic of human AD (*88*–*91*), their brains activate similar molecular pathways with increasing age, highlighting that early, conserved mechanisms of neuroinflammatory and neuronal decline may underlie both healthy and pathological aging.

We found that sex contributes substantially to heterogeneity in brain aging, primarily through cell type-specific gene expression, rather than through changes in cell composition. More than half of all expressed genes showed sex-biased expression in at least one cell class or region, with glial populations – particularly astrocytes and oligodendrocytes – exhibiting the most extensive sex differences. Strikingly, many sDEGs were also differentially expressed with age and many cell types showed clear biases in terms of which sex was transcriptionally “older”. Across most cell types and regions, we observed a consistent “female-older” transcriptional pattern, whereby female-biased genes tended to increase with age, while male-biased genes tended to decrease. This may reflect female-biased activation of cellular stress or maintenance pathways, potentially contributing to the sex differences in some age-associated neurodegenerative diseases, including AD (*93*). Conversely, MSNs – key to dopaminergic signaling and motor control – consistently exhibited a “male-older” pattern, revealing an age-associated increase in lipid-related functions. This pattern may contribute to the higher prevalence of Parkinson’s disease in men (*79*–*81*) and reflect the involvement of lipid-related functions in PD, as shown in observational and postmortem human studies (*94*, *95*) and in mouse models (*96*–*100*).

In addition to these additive effects of sex and age, we found that sex also modifies age-related gene expression in a cell-type- and region-specific manner. Although not ubiquitous, a*sDEGs were broadly distributed across the brain and disproportionately enriched in oligodendrocytes and glutamatergic neurons, and in the dlPFC. The additive and interactive effects of age and sex on gene expression identified here are concordant with human studies, which found considerable additive and limited interactive effects of age and sex on gene expression (*92*). Notably, most a*sDEGs showed stronger age effects in females, suggesting that transcriptional remodeling with age is often accelerated in female brains. One illustrative example is *IQCK*, an AD risk gene whose expression increased more strongly with age in astrocytes in females compared to males. These findings indicate that sex-by-age interactions can amplify molecular differences across the lifespan, potentially contributing to sex-biased vulnerability to disorders such as AD.

There are several limitations to this study: 1) Although we used a Poisson log-normal model to estimate differences in cell abundance and mitigate the effects of compositionality, some compositional bias may persist due to undersampling. In this context, large shifts in the abundance of one cell type could artifactually inflate or deplete the inferred abundance of other cell types; 2) We applied an isotropic significance threshold across our statistical tests, which may limit detection of differential abundant cell types or differentially expressed genes/pathways in conditions that are underpowered. 3) The inference of non-linear dynamics is constrained by the variability observed across samples, particularly during mid-life. Increasing the density of sampling and including additional individuals will strengthen future analyses of non-linear trajectories.

In summary, our multi-region single-cell transcriptomic atlas of the macaque brain reveals how age shapes cell abundance and gene regulation. While many differences are shared across regions, we identify cellular and molecular dynamics that are differentially regulated in specific cell classes, brain regions, and in a sex-dependent manner. This work offers a novel resource for understanding the molecular basis of vulnerability to neurodegenerative diseases and informing therapeutic strategies. We highlight vulnerable cell subclusters, gene programs that converge with neurodegenerative disease signatures, and sex-specific regulatory effects in regions and cell types implicated in disease. We anticipate that this atlas will serve as a useful reference for advancing our understanding of brain aging and its intersection with neurodegeneration.

## Methods

### Study population and sample collection

We analyzed postmortem brain samples from 55 rhesus macaques (*Macaca mulatta*; 29 females, 26 males) ranging in age from 5 months to 21 years. All animals belonged to the semi-free-ranging colony on Cayo Santiago, Puerto Rico (18°09′N, 65°44′W), a long-term research population maintained by the Caribbean Primate Research Center (CPRC) of the University of Puerto Rico. The colony was founded in 1938 and has since been continuously monitored (*10*). Animals live in social groups under naturalistic conditions, with daily provisioning and *ad libitum* access to water, and without medical intervention aside from tetanus inoculations during scheduled trapping events. No contraceptives are used, and animals reproduce freely (*101*).

As part of routine colony management and population control, the CPRC periodically removes animals (*9*). The Cayo Biobank Research Unit (CBRU) coordinates tissue collection from these removal events (*8*, *13*, *102*, *103*). All animals included in this study were euthanized for colony management purposes by CPRC veterinary staff in accordance with approved protocols (University of Puerto Rico IACUC #338300). Immediately following euthanasia, brains were removed, hemisected, and coronally sectioned into 11 blocks using custom molds. Each block was flash-frozen in liquid nitrogen vapor and stored at −80°C and regions dissected as described in (*13*). All of the samples for this manuscript were dissected from the right brain hemisphere.

### snRNA-seq data generation

To minimize within-region batch effects, nuclei isolation and snRNA-seq library preparation were performed in 11 batches, one per region. Nuclei were isolated and fixed using a modified version of the optimized sci-RNA-seq3 protocol (*12*). Briefly, tissue was manually pulverized on dry ice, transferred to pre-chilled tubes, and stored at −80°C. For lysis, pulverized tissue was suspended in hypotonic lysis buffer A (0.3 M sucrose, 0.1% Triton X-100, 3 mM MgCl₂, 0.025% IGEPAL CA-630, 1% DEPC in PBS) for 10 min, filtered (40-μm), pelleted, and resuspended in SPBSTM buffer (0.3 M sucrose, 1X PBS, 0.1% TritonX-100, 3 mM MgCl₂, and 1% DEPC). Nuclei were fixed for 10 min in 1.5 mM dithiobis(succinimidyl propionate) (DSP), diluted from a 123 mM stock in DMSO into methanol. After washing in 0.3 M SPBSTM, nuclei were layered onto a 10:1 volume of 1.4 M SPBSTM, centrifuged (3,000 x g, 20 min, 4°C), and resuspended. Nuclei were filtered (20-μm), counted, aliquoted, and cryopreserved in 90% 0.3 M SPBSTM with 10% anhydrous DMSO at -80°C.

Aliquots were shipped on dry ice to the Brotman Baty Institute Advanced Technology Lab (BAT-Lab) at the University of Washington for library preparation (*12*, *104*). One library per region was generated using the optimized sci-RNA-seq3 protocol with the following modifications: nuclei were counted using the ImageExpress Pico System (Molecular Devices), and 15,000-20,000 nuclei were loaded per reverse transcription well. Tagmentation used N7 adaptor-loaded Tn5 transposase (Diagenode, C01070010-20), with 6-9 μL of Tn5 per 550 μL of 2X TD buffer (20 mM Tris-HCl pH 7.6, 10 mM MgCl₂, 20% dimethylformamide), distributed at 5uL per well.

Libraries were cleaned using a two-step bead purification: first with 0.8X AMPure XP beads to remove lower molecular weight fragments, followed by a 0.55X bead-based supernatant step to remove larger fragments. Final purification was performed using Qiagene GeneRead Size Selection columns. Libraries were sequenced on Illumina NovaSeq S4 flow cells at the University of Washington Northwest Genomics Center, using the following settings: Read 1 - 34 cycles, Read 2 - 100 cycles, Index 1 - 10 cycles, Index 2 - 10 cycles.

### snRNA-seq preprocessing

Read alignment and gene count matrix generation was performed using the BAT-Lab pipelines for sci-RNA-seq3 (https://github.com/bbi-lab/bbi-dmux; https://github.com/bbi-lab/bbi-sci). Briefly the pipelines include the following steps: (i) converts base calls to fastq files with bcl2fastq/ v.2.20 (RRID:SCR_015058) (Illumina), (ii) removes polyA tails using Trim Galore/v.0.6.7 (RRID:SCR_011847) (99), (iii) aligns trimmed reads to a reference genome with STAR/v.2.7.6 (RRID:SCR_004463) (*105*), (iv) extracts mapped reads, (v) removes duplicates, and (vi) generates UMI counts for exonic and intronic regions of each gene, tabulated according to the unique three-level barcode design in sci-RNA-seq3. The rhesus macaque reference genome (Mmul_10) (*106*) and Ensembl (version 101) (RRID:SCR_002344) annotation were used as our reference. The 3’ untranslated region annotations of genes and transcripts were extended by 500 bp to avoid misclassifying genic reads as intergenic.

After generating the count matrix, we removed nuclei of low quality or ambient RNA with the following standard: (i) UMI counts < 100, (ii) number of gene detected ≤ 100, (iii) percentage of mitochondria ≥ 10%, (iv) percentage of reads mapping to intronic region ≥ 20%. For each sample, we then imported the filtered gene-by-nucleus count matrices into the AnnData/v.0.8.0 (RRID:SCR_018209) (*107*) framework and then ran Scrublet/v.0.2.3 (RRID:SCR_018098) (*107*, *108*)(expected_doublet_rate = 0.05) to calculate doublet scores. We marked nuclei as Scrublet-inferred doublets if they had Scrublet doublet scores > 0.20.

To identify doublets with the scrublet output, we used an iterative clustering strategy using Scanpy/v.1.9.1 (RRID:SCR_018139)(*109*) implemented by bioalpha(*110*) on the ASU Research Compute Cluster (*111*). First, we combined the count matrices of all samples from the same brain region into a single AnnData object. Second, we normalized the data to the total UMI per nucleus, logarithmized the data, and subsetted the data to the 10,000 most variable genes. For each cell, we regressed out total UMI counts per nucleus and then mean-centered and scaled the data. The dimensionality of the data was then reduced by principal components analysis (PCA) (50 components). We then constructed a neighborhood graph using 30 nearest neighbors (n_neighbors=30) and 30 principal components (n_pcs=30), and performed Louvain clustering at a high resolution (resolution = 6) to partition the nuclei into small clusters. Finally, we calculated the proportion of Scrublet-inferred doublets in each cluster and observed a bimodal distribution across all clusters, with a clear separation of the two peaks at 40%. Therefore, we filtered all nuclei in clusters with ≥ 40% of Scrublet-inferred doublets.

To further filter out nuclei of low quality or doublets that are remaining, we implemented a distribution based approach. For putative non-neuronal and neuronal nuclei, we modeled their UMI count distributions separately and computed the mean and standard deviation (SD) for each group. Nuclei with UMI counts ≤ mean + 1 SD and ≥ mean + 2 SD were excluded from further analysis.

After the filtering steps as described above, we repeated the normalization and dimensionality reduction. We then visualized the filtered dataset by UMAP (min_dist=0.25,spread=1.0, n_components=2). We confirmed adequate removal of doublets by observing the clean separation of distinct cell types and the absence of clusters expressing obviously ambiguous marker gene profiles.

### snRNA-seq cell class and cell subcluster annotation

#### Cell classes

For each region, we annotated the cell class by using a combination of label transfer and majority ruling. First, we annotated each cell to its putative cell class by an optimal transport based label transfer method implemented by TACCO (*112*). We performed label transfer with all cell classes present in the corresponding region in the BICCN taxonomy (*13*) as the reference. Due to substantial increase in UMI counts in the current dataset compared to the BICCN reference, we normalized our data to conform to the reference (normalize_to=’reference’). After annotation with TACCO, we applied a majority ruling approach on the Louvain clusters (resolution = 6) generated above. We annotated a cluster to a particular cell class if >70% of the nuclei in the cluster belonged to the same class, or the proportion of the top annotated cell class was 3x higher than the second annotated one. Clusters that were not annotated with the majority ruling were subjected to manual annotation based on marker gene expression and subcluster annotation described below (1-15% of nuclei in each region). Unknown clusters with high mean doublet scores (>0.15) and inconsistent annotation between cell class and subcluster were marked as doublets and excluded (0-8% of nuclei in each region). We also identified two cell classes that were not expected in a certain region (i.e. basket cells in midbrain, medium spiny neurons in hippocampus), likely coming from dissection errors. They were kept for cell abundance analysis at the cell class level to account for unidentified cells from dissection errors in these regions, but excluded in other analyses.

#### Cell subclusters

After finalizing the annotation of the cell class in each region, we combined nuclei of the same cell class among regions to generate subclusters under each cell class. We performed normalization and dimensionality reduction as above except subsetting to 3,000 most variable genes. We then constructed a neighborhood graph as above and performed Louvain clustering (resolution = 1) for each cell class. For each cell class, we then annotated the cell subclusters by label transfer implemented by TACCO as described above. We used BICCN (*13*) and human brain datasets (*14*, *15*) as reference. We also compared the Allen mouse brain atlas by MapMyCell(*113*). For microglia and astrocytes subclusters, we have performed additional functional annotation by comparing their transcriptomes to external datasets (*22*) and Gene Ontology with msigDB (*114*, *115*). After generation and annotation of cell subclusters, we combined all nuclei into a global AnnData object (∼5.3M nuclei). We then inspected the localization of subclusters annotated to the same cell class. For subclusters that were not localizing closer to their annotated cell class, we reassigned their cell class and subcluster labels based on marker gene expression (1% of all subclusters).

### TF Regulon analysis

To identify TF regulons that are enriched in cell subclusters, we have performed single-cell regulatory network inference and clustering (SCENIC) analysis with pySCENIC (*116*). We filtered macaque genes in our dataset to retain only those with one-to-one human orthologs, and then mapped the macaque genes to their corresponding human orthologs using orthology annotations from Ensembl BioMart (release 113). To accelerate the analysis, we filtered out genes expressed in <100 cells and subset cell class objects to 100k cells. We followed the pipeline of SCENIC, which in briefly: i) generated co-expression modules based on positive correlation between transcription factors (annotated in hg38 human genome) and candidate target genes with GRNBoost2; ii) each co-expressed module was analyzed with cisTarget using v10 human motif databases and annotations to identify enriched motifs: modules and targets for which the motif of TF is enriched retained; iii) the activity of each regulon in each cell was evaluated using AUCell, which calculated the Area Under the recovery Curve. We evaluated the activity of a particular regulon in a subcluster by averaging the regulon activities across all cells.

### Cell composition analysis

#### Poisson Lognormal models (Hooke)

To perform differential analysis of cell state abundances, we modeled cell abundance change with Poisson Lognormal (PLN) models implemented in Hooke (*25*). We retained only adult animals (age>3 years) for modeling linear abundance change. For each region, we first converted our single-cell dataset into a matrix containing the number of cells of each cell class/subcluster observed in each sample, along with metadata for each sample. Then Hooke estimates per-sample “size factors” that will control for sample-to-sample variation in cell capture levels. Next, it initialized a PLN model that estimates proportions of each cell state as a function of age and sex by new_cell_count_model(vhat_method = ‘bootstrap’, num_bootstraps = 100). We extracted the coefficients and standard errors from the full model and calculated two-sided p-values under the null hypothesis that each coefficient equals zero, using the Student’s t-distribution. We then performed false discovery rate (FDR) correction on the resulting p-values and considered effects significant at an adjusted p-value < 0.05. To maintain sufficient statistical power, we excluded region-cell subcluster combinations with fewer than 100 cells from the results of differential analysis within cell class. To visualize non-linear abundance change, we input cell count in all animals into the PLN models and extract the ‘normalized counts’ after normalization with size factor.

#### Multivariate adaptive shrinkage (MASH)

To further boost sensitivity, especially in regions with low cell counts, we applied multivariate adaptive shrinkage (MASH) through the R package mashR (*29*). We imported the coefficients output and standard errors from Hooke and applied the data-driven approach: (i) selected strong signals (local false discovery rate, LFSR<0.05) from running a condition-by-condition analysis on all the data; (ii) performed PCA and compute initial covariance matrices based on the top 5 PCs of the strong signals; (iii) applied the Extreme Deconvolution algorithm from the initialized matrices to the strong signals; (iv) fitted mash to all tests.

### Pseudotime analysis

The pseudotime analysis was performed with Monocle3 (*104*). We followed the instruction to (i) perform normalization and dimensionality reduction of the full dataset with default parameters; (ii) cluster cells into partitions using a statistical test (*117*); (iii) for each partition, fit a principal graph and set the root node to the subclusters with abundance decrease over age; (iv) calculate pseudotime values for each cell using order_cells.

### Differential expression analysis

#### Linear mixed model (NEBULA)

To assess differential gene expression by age and sex across combinations of cell subclusters and brain regions, we applied the NEBULA framework, which fits a negative binomial gamma mixed model (NBGMM) using a large-sample approximation. Following NEBULA guidelines, we excluded subcluster-region combinations with fewer than 30 animals. Within each combination, we filtered out lowly expressed genes using a threshold of counts per cell (cpc) < 0.05 (i.e., expressed in fewer than 5% of cells). The model included age, sex, and UMI counts as fixed effects, and animal ID as a random effect. We performed false discovery rate (FDR) correction on the resulting p-values and considered genes significant at an adjusted p-value < 0.05. To further improve statistical robustness, we applied a post-hoc heuristic filter, excluding combinations with fewer than 300 cells, resulting in 820 unique combinations of region and cell subclusters. This threshold was based on the minimum cell count at which a linear relationship between cell number and the number of detected differentially expressed genes became evident. To model differential gene expression as an effect of the interaction between age and sex, we filtered the inputs to test genes that were either an aDEG or sDEG (LFSR < 0.05).

#### Multivariate adaptive shrinkage (MASH)

To further boost sensitivity, especially in combinations of cell subclusters and regions with low cell counts, we applied multivariate adaptive shrinkage (MASH) as above among all combinations of cell subclusters and regions within each cell class. For sex-specific analysis, sDEGs on the autosomes and sex chromosomes were input into mashR separately.

### Gene set analysis

#### Gene set enrichment analysis

All age-associated differentially expressed genes (aDEGs) identified by NEBULA were ranked in ascending order based on their local false discovery rate (LFSR) values (-fold change sign x log10(LFSR+1e-43)) obtained from MASH. This ranked gene list was then used for gene set enrichment analysis (GSEA) using the GSEA function from the clusterProfiler package(*118*). Enrichment analysis was conducted using gene sets from msigDB (*114*, *115*), KEGG (*119*, *120*) and synGO (*119*). The significance score was calculated by -NES sign x log10(p-value of a given pathway). For significance score at the cell class level, we then summarized their mean significance score by cell class and region.

#### Gene set over-representation analysis

Before performing gene set ORA, we classified a+sDEGs as “female older” or “male older.” Female-older genes are those that show a male-biased gene expression and decrease with age, or a female-biased gene expression and increase with age, and vice-versa for male-older genes. In other words, genes were grouped into male-older or female-older based on the overall trend of their age-associated expression and whether males or females show stronger expression in that direction. For example, if a male-biased gene increases in expression with age, then the overall trend is that the gene is more highly expressed in older animals and males, thus males are transcriptionally older. If the gene is more highly expressed in females but decreases in expression with age, then the overall trend is decreased gene expression in older animals and thus males, having lower expression across the lifespan, are transcriptionally older in their expression.

We used the “enricher” function of the clusterProfiler R package (*118*) to perform a hypergeometric test on all genes in a given subcluster-region that showed a significant effect of age and sex on their expression (a+sDEGs). The background set of genes for this analysis was all the genes included in the NEBULA DEG analysis for that subcluster-region combination. This test was performed on synGO gene sets (*119*), as well as the Hallmark, KEGG Medicus, BioCarta, GO:BP, GO:MF, GO:CC, PID, Reactome, and WikiPathways collections from msigDB (*115*).

For analysis, results were filtered for FDR-adjusted p-value < 0.05 and number of overlapping genes > 1. Gene sets passing these filters (n = 2475) were sorted into broader groups (e.g., lipid metabolism, synaptic function) based on their description with assistance from Claude Sonnet 4.5 (Anthropic, 2025) (*121*), with manual verification and modification.

## Acknowledgments

We thank the management and staff of the CPRC (particularly A. Ruiz-Lambides, C. Sariol, and A. Burgos-Rodríguez) for maintaining the Cayo Santiago and Sabana Seca field stations. We also thank members of the CPRC (particularly S. Bauman, O. Gonzalez, N. Compo, C. Pacheco, and S. Gascot Maldonado), the CBRU biobanking team (C. Walker and J. Stylli), the Snyder-Mackler Lab (B. Slikas, S. Ford, A. Greenier, M. Koperska, C. Adjangba, S. van Djik, L. Brassington, and T. Zintel), and the Platt Lab (L. Assi) for assistance with sample collection and/or logistics. We are grateful to Troy McDiarmid, Embla Størdal, Tony Li, Xiaoyi Li, Xingfan Huang, Riddhiman Garge, H. Pliner, B. Ewing, and A. Buckley for assistance and/or feedback through various stages of this manuscript and Gil Speyer and the ASU Research Computing team for computational resources and support. The data reported here were generated via the single-cell platform of the Brotman Baty Institute (BBI) Advanced Technology Lab (BAT-Lab) at the University of Washington. This publication was supported by and coordinated through the Brain Initiative Cell Atlas Network (BICAN).

## Author contributions

N.S.-M., J.S., M.L.P., K.L.C., M.J.M., L.M.S., W.Y and K.L.W conceived the study. K.L.C., M.J.M., and N.S.-M. collected samples, with logistical support from Cayo Biobank Research Unit and M.I.M. M.O.B. performed neuroanatomical dissections, assisted by K.L.C., M.J.M., and N.S.-M. K.L.C., K.L.W, M.O,-N, D.R.O. and A.M performed lab work. A.V managed data and provided bioinformatic support. W.Y, K.L.W, K.L.C. and N.S.-M. analyzed the data, with input from A.R.D, M.O,-N, L.M.S., M.J.M., M.L.P., and J.S. W.Y, K.L.W, K.L.C., M.O.B., A.R.D., M.J.M., M.L.P., J.S., and N.S.-M. wrote the paper. All authors edited and approved the manuscript. **Consortium authors:** The members of the Cayo Biobank Research Unit are S. C. Antón, L. J. N. Brent, J. P. Higham, M.I.M., A. D. Melin, M.J.M., M.L.P., J. Sallet, and N.S.-M.

## Data availability

Data will be publicly available on GEO upon peer reviewed publication.

## Code availability

All code for this study is accessible through the following GitHub repositories: https://github.com/CayoBiobankResearchUnit/macaque-aging-series (data analysis), https://github.com/bbi-lab/bbi-dmux (sci-RNA-seq3 data demultiplexing), and https://github.com/bbi-lab/bbi-sci (sci-RNA-seq3 data preprocessing).

## SUPPLEMENTARY TABLES

[Extended data table](https://docs.google.com/spreadsheets/d/1e0wAI2ZAiBqRiY3bzETTQqvpWymoViHRHky5JYQpHps/edit?usp=sharing)

## SUPPLEMENTARY FIGURES

**Figure S1.**
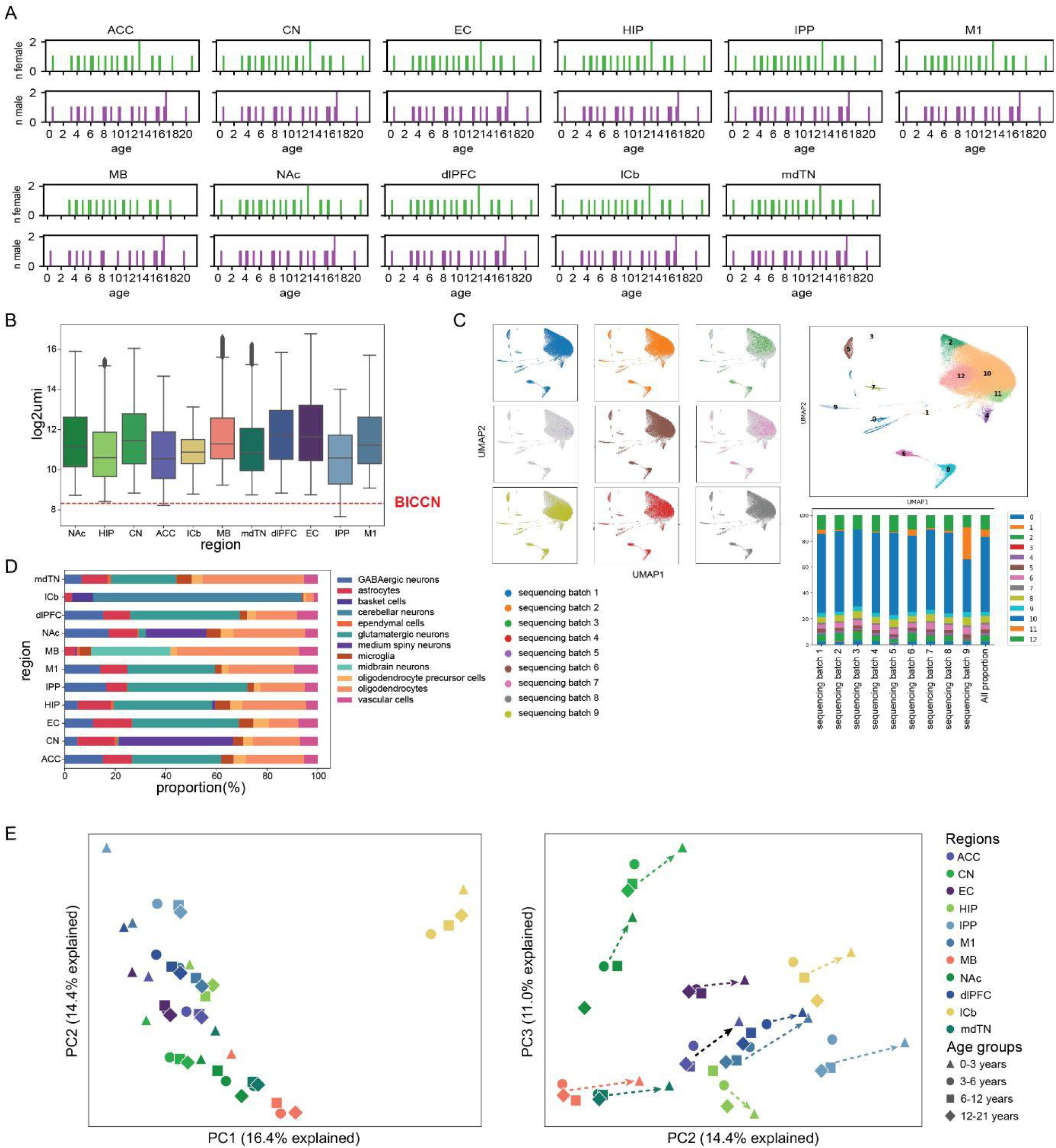
Quality control and principal component analysis. **A)** Age and sex distribution of samples collected for each brain region. **B)** Boxplot showing distribution of number of unique molecular identifiers (UMIs, log2-transformed) recovered in each cell collected by region. The red dotted line shows median UMI recovered in previous BICCN atlas(*13*) **C)** Joint UMAP projections of control samples (cerebellum) from different sequencing runs, colored by the sequencing run ids where they were loaded. Unsupervised clustering was performed on the jointed dataset. Barplots showing proportion of control samples from different sequencing runs in each Louvain cluster and all cells. **D)** Barplot showing proportion of cell class across regions. **E)** For each region, transcriptional profiles were aggregated to create pseudobulk transcriptomes of 4 age groups (i.e. infants: 0-3 years, adolescence: 3-6 years, adulthood: 6-12 years, aged: 12-21 years). Principal component analysis (PCA) was performed on the resulting pseudobulk transcriptomes. PCA plots showing PC1 and PC2 (upper), PC2 and PC3 showing (lower). PC1 captures differences between cerebellum and all other brain regions. PC2 and PC3 capture differences among regions and age trends within each region. The dotted line indicates huge gaps between transcriptomes of infant samples (0-3 years) and transcriptomes of adult samples (>3 years).

**Figure S2.**
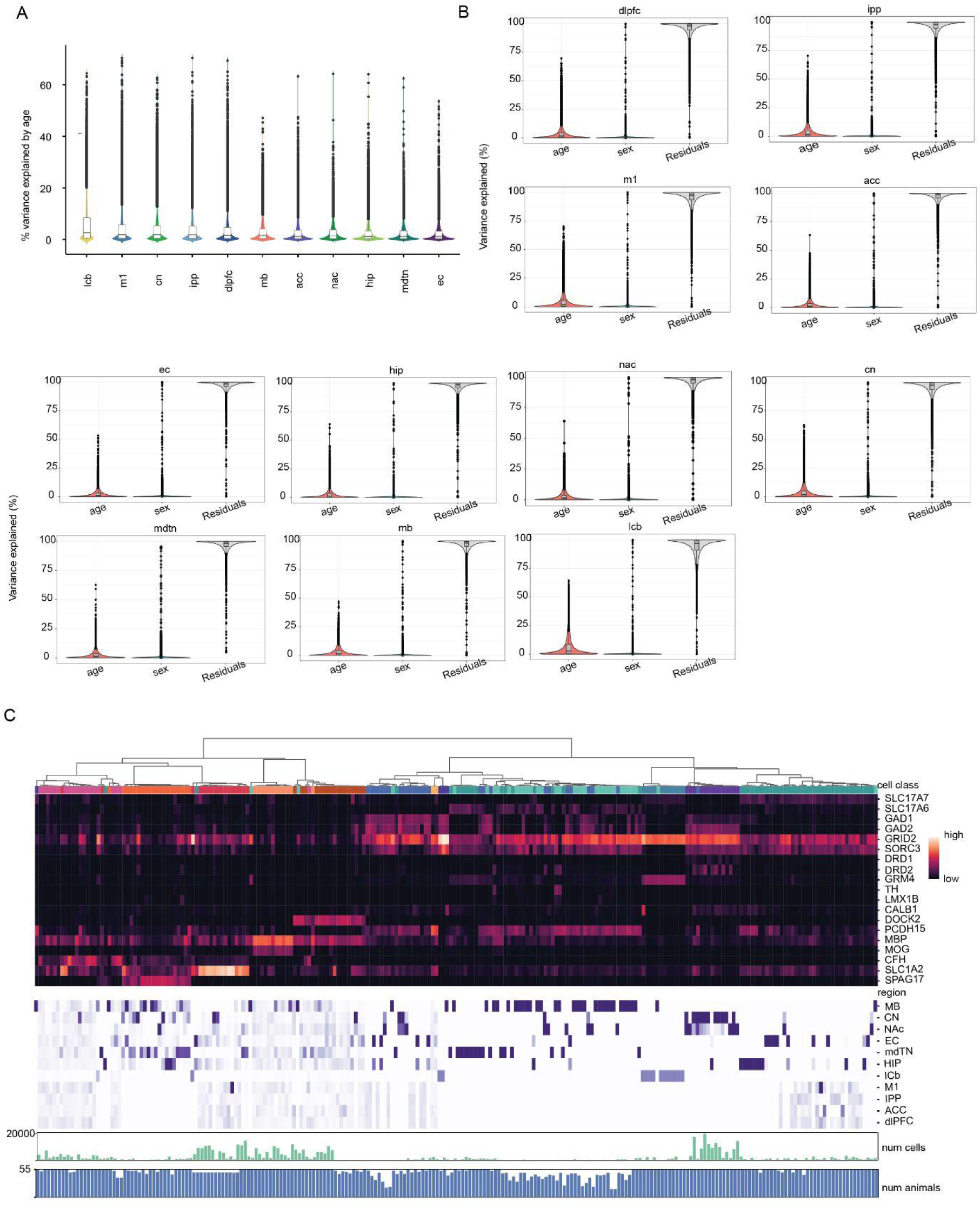
Variance partitioning and sub-clustering statistics. **A)** For each region, transcriptional profiles were aggregated to create pseudobulk transcriptomes for each individual. Violin plot showing percentage variance explained by age for each gene in individual regions, ordered by highest to lowest median percentage. **B)** For each region, transcriptional profiles were aggregated to create pseudobulk transcriptomes for each individual. Violin plot showing percentage variance explained by age and sex for each region for each gene. **C)** Dendrograms showing hierarchical clustering of subclusters labeled by cell class (colors are the same as Fig 1B). Heatmap showing cell class marker gene expression among subclusters and distribution among regions. Barplots showing number of cells in each subcluster and number of donors with this subcluster in at least one region.

**Figure S3.**
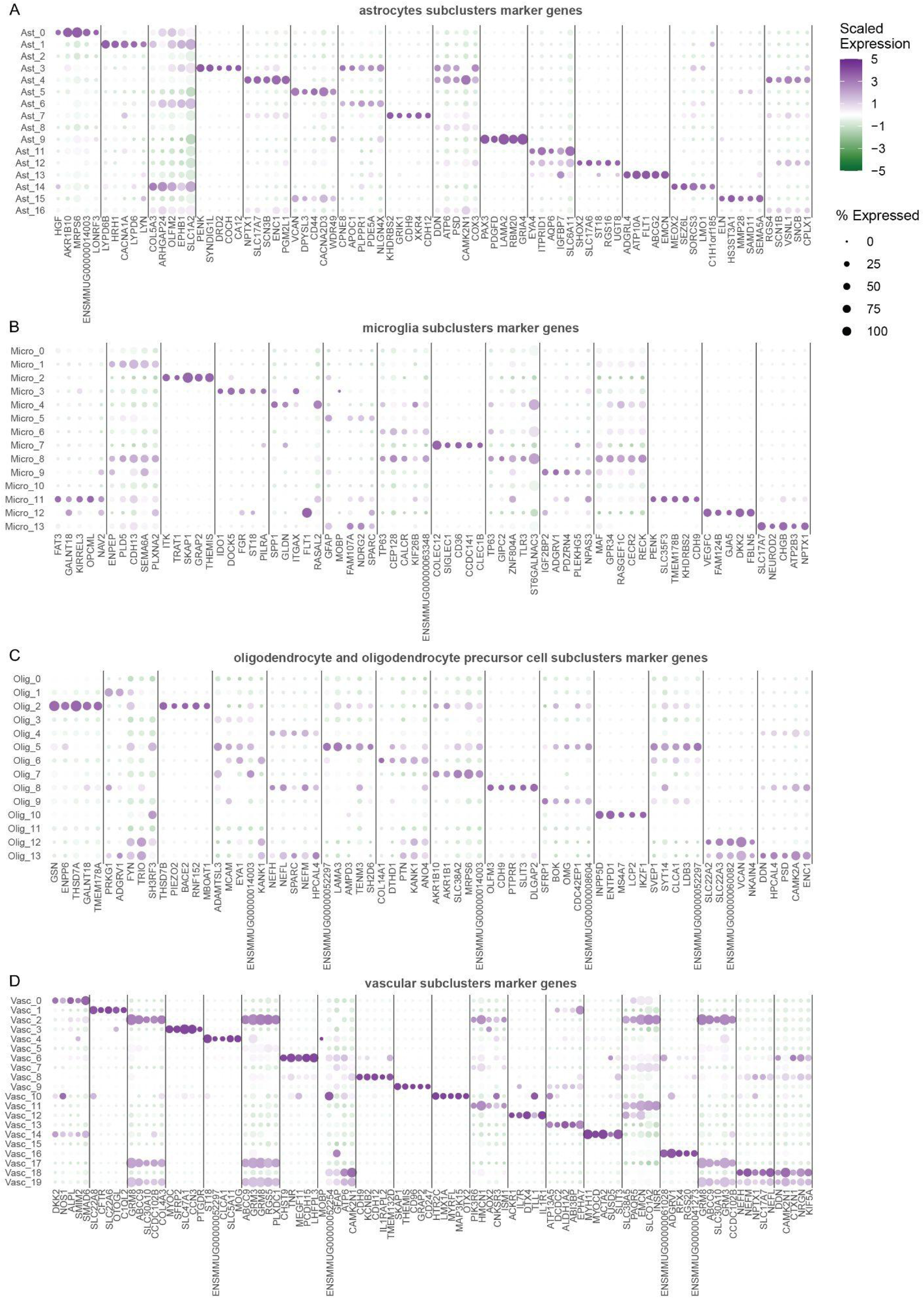
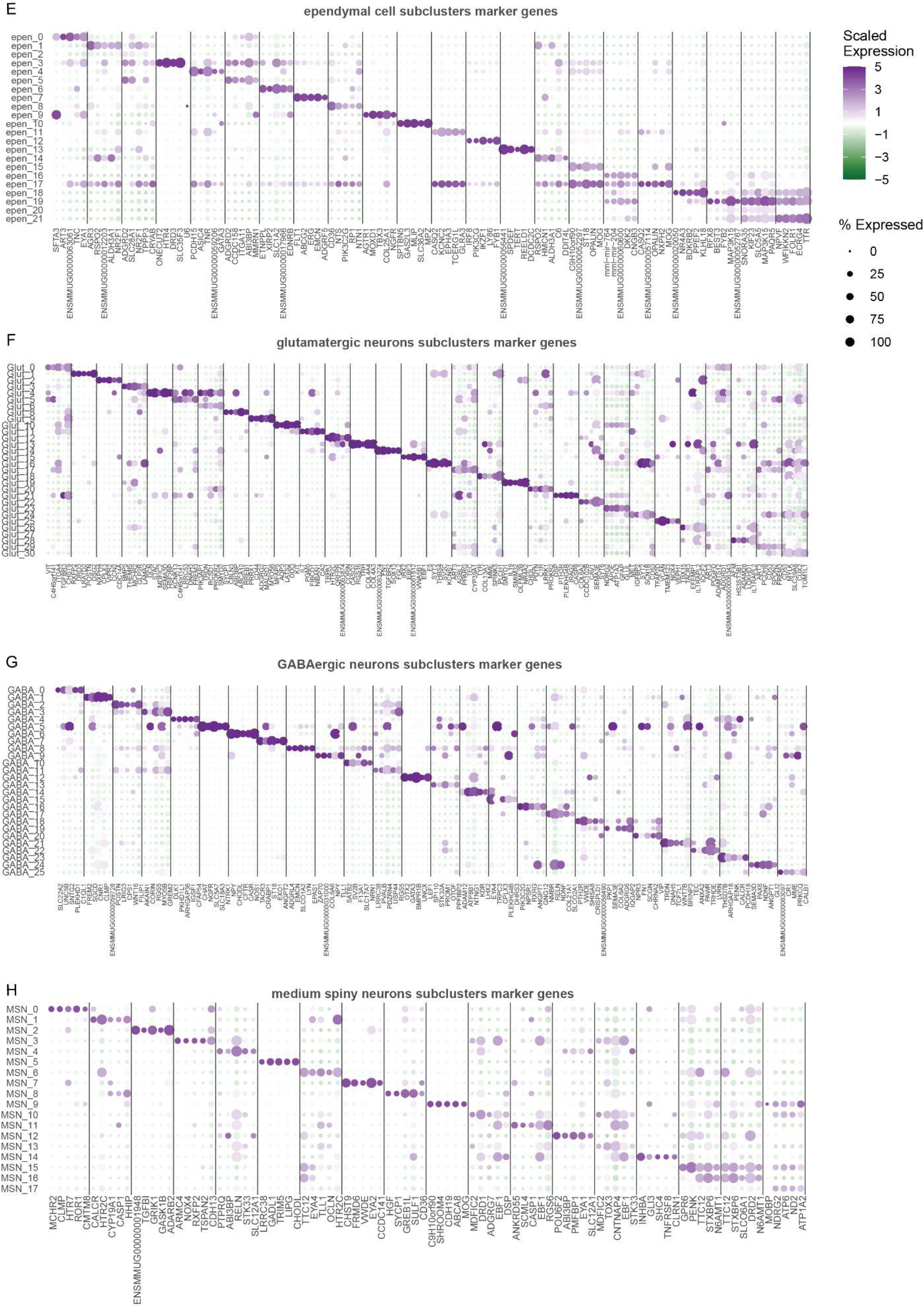
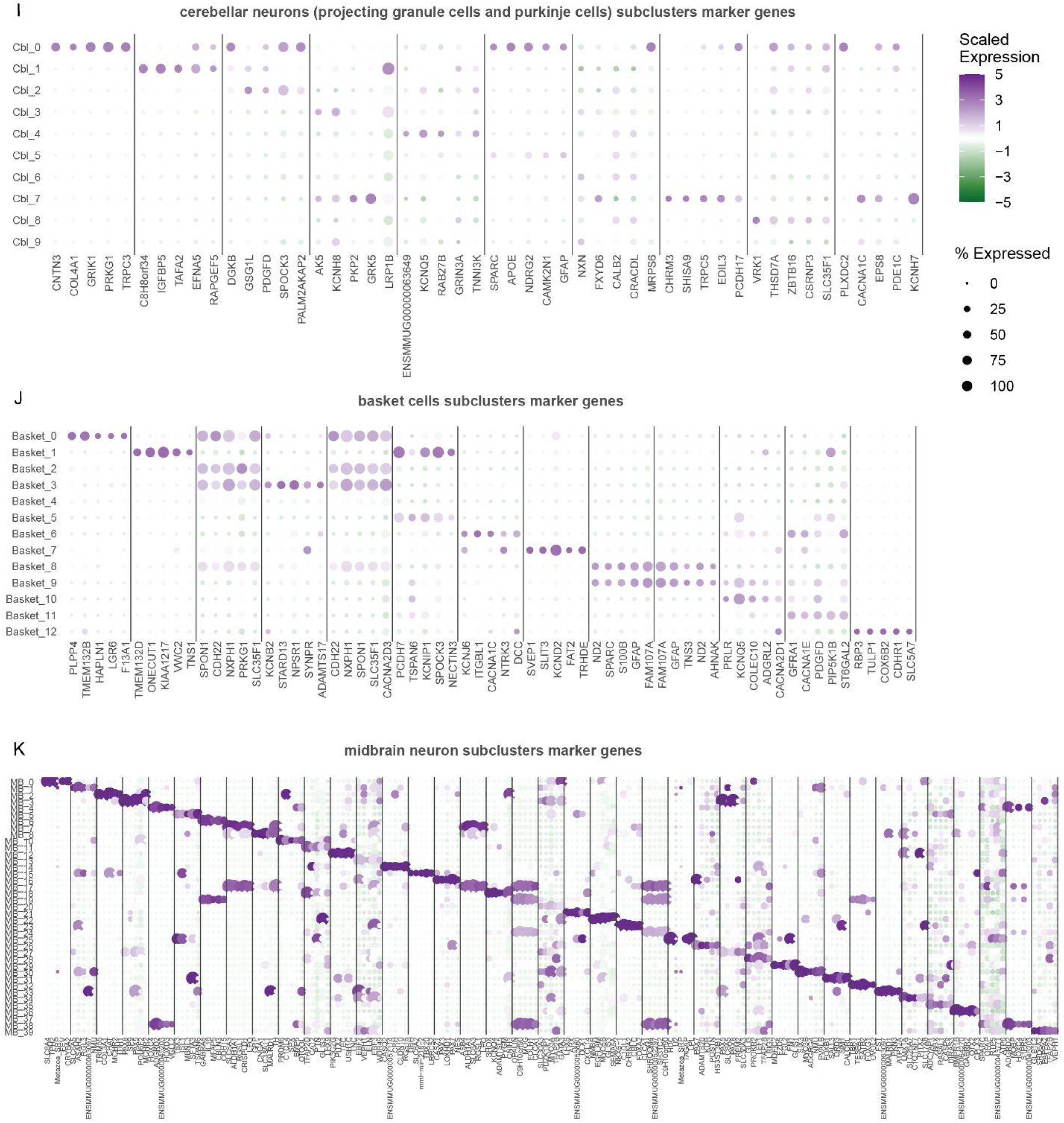
Marker genes of cell subclusters. Subclusters were identified by unsupervised clustering for each non-neuron cell class (OPC and oligodendrocytes were analyzed together) and for neurons from each of the 6 anatomical regions (table S4). The top 5 marker genes with the highest z-score and log-fold change from a Wilcoxon test were selected as the marker for each subcluster.

**Figure S4.**
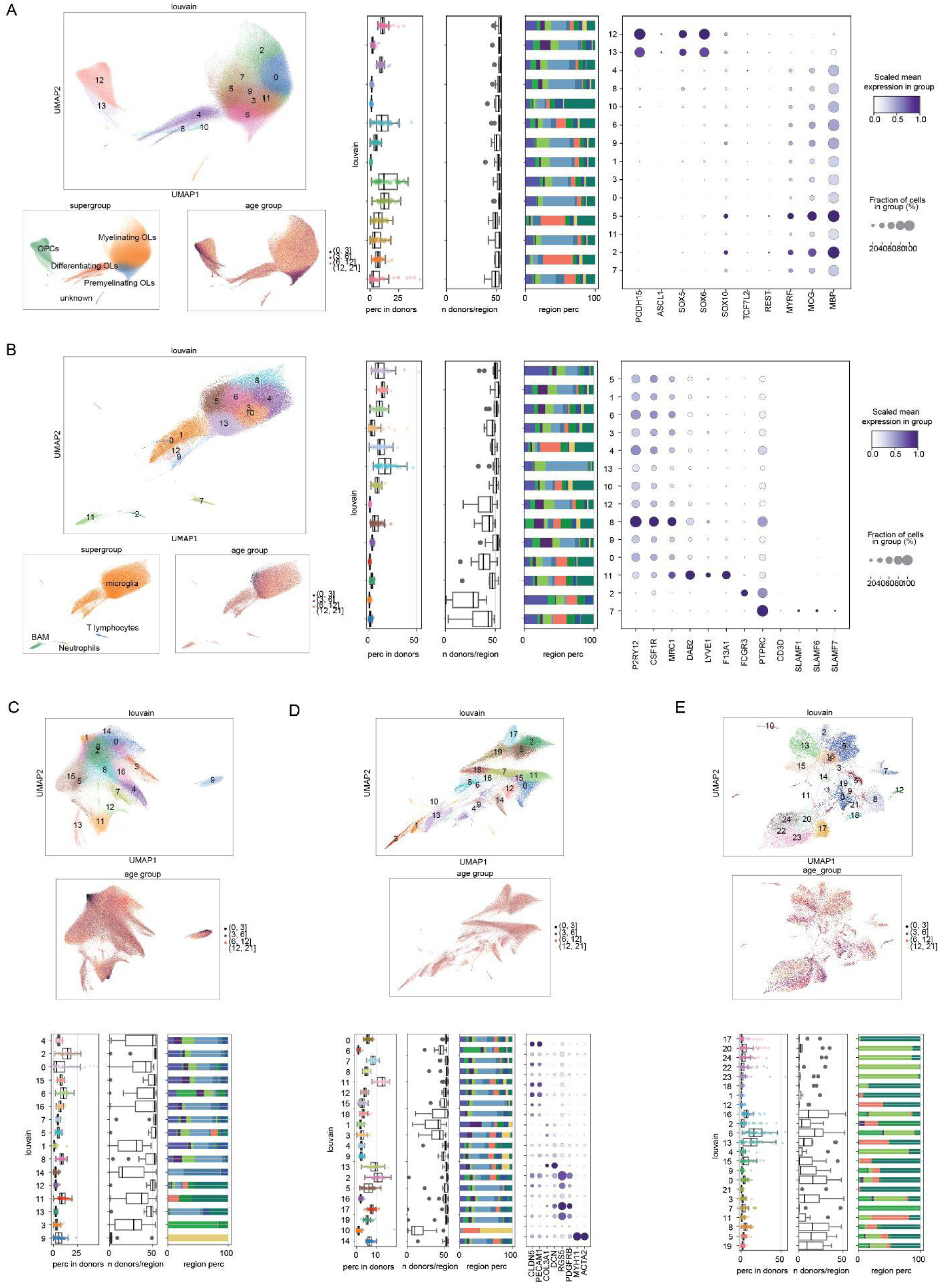
Donor specificity, region representation and known cell subtype marker genes. UMAP visualization of oligodendrocytes **(A)**, microglia **(B)**, astrocytes **(C)**, vascular cells **(D)**, ependymal cells **(E)** subclusters identified by Louvain unsupervised clustering, annotation of supergroups, and age groups. Boxplot showing proportion of each subcluster in the total population of glutamatergic neurons in each donor. Boxplots showing the number of donors with this subcluster in a specific region. Subclusters showing high differences in donor numbers among regions are region-specific subclusters. Barplots showing proportion of subclusters coming from each region. Dotplot of scaled mean expression of marker genes for known cell subtypes in each subcluster.

**Figure S5.**
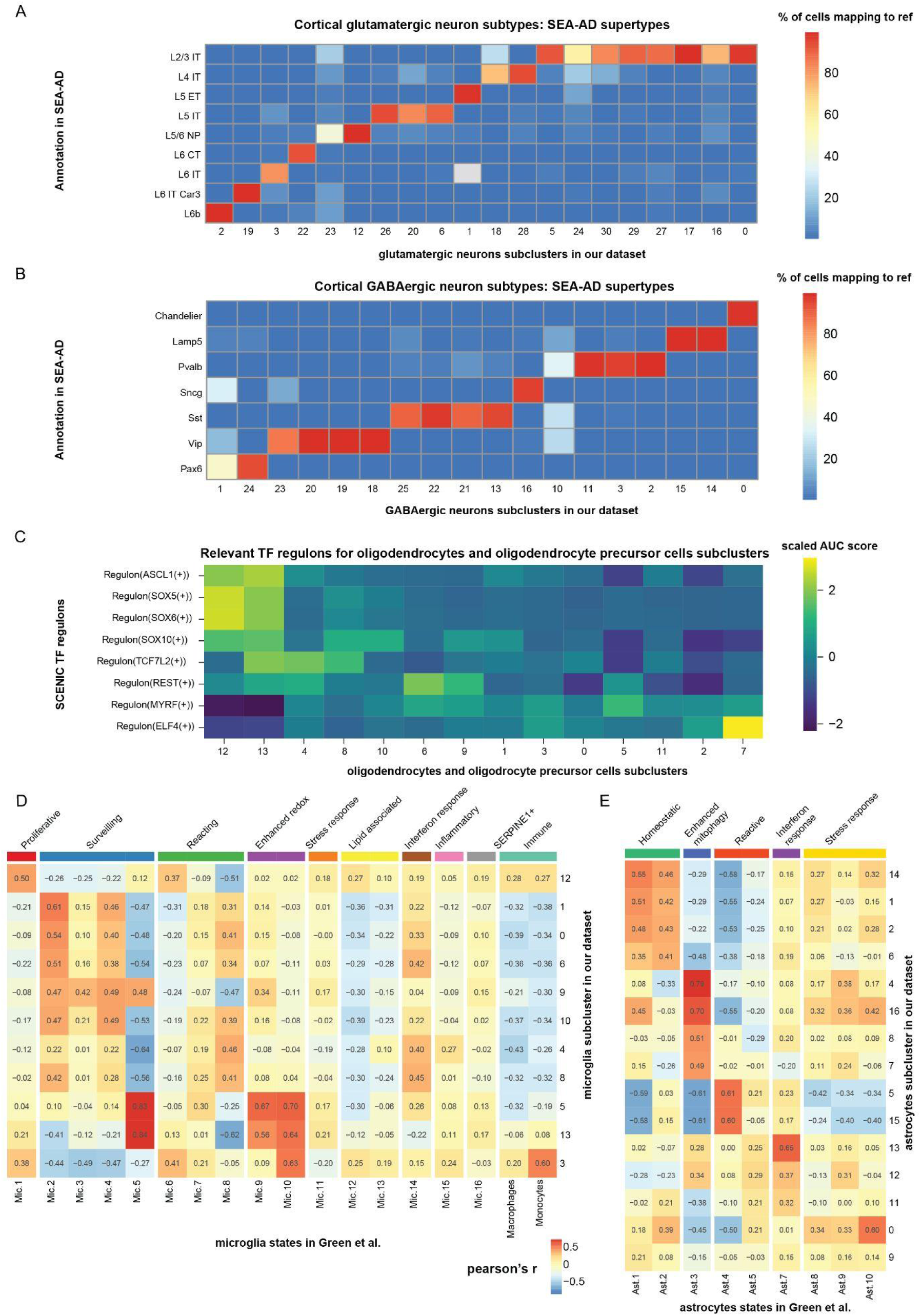
Mapping of subclusters to external datasets. **A)** Heatmap showing percentage of cells in our glutamatergic neuron subclusters mapping to SEA-AD excitatory cortical subtypes with label transfer by TACCO (method). **B)** Heatmap showing percentage of cells in our GABAergic neuron subclusters mapping to SEA-AD inhibitory cortical subtypes. **C)** TF regulons and their AUC scores computed by SCENIC for each subcluster in oligodendrocyte precursors and oligodendrocytes, scaled by row. **D)** Heatmap of pearson’s correlation of average log2FC of marker genes in our microglia subclusters and those in the distinct microglia states reported by Green et al. (*22*). **E)** Same as panel (D) but showing correlations with different astrocyte states.

**Figure S6.**
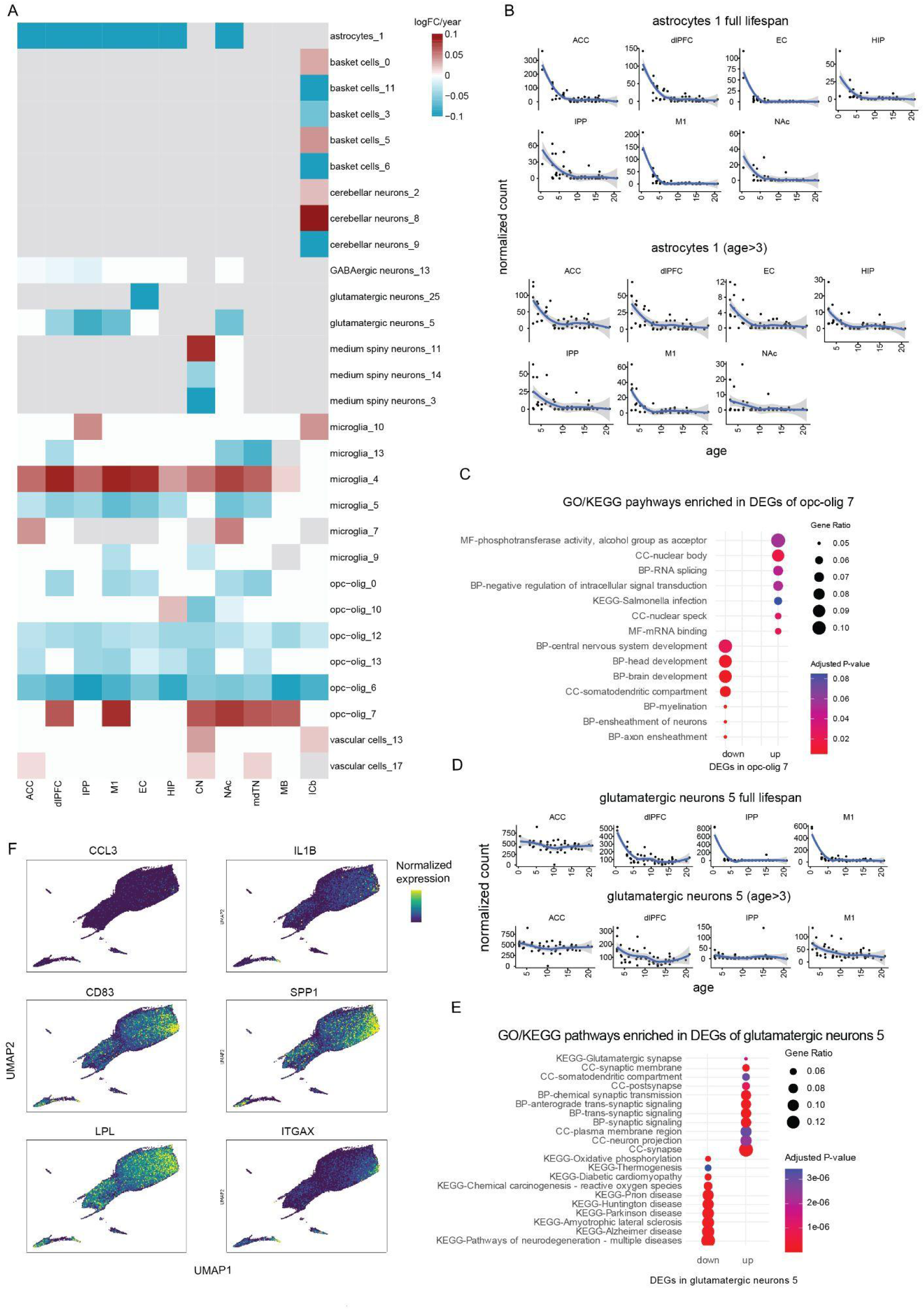
Cell abundance change for subclusters and GO term enrichment. **A)** Heatmap showing cell abundance change across different cell subtypes and regions. Value represents log-fold differences per year in the latent abundance from the poisson log-normal model implemented by Hooke(*25*). **B)** Loess fit of the latent abundance of astrocytes 1 across macaques of all ages (upper), and of age > 3 (lower). **C)** GO term and KEGG pathway enrichment of up and down regulated genes of OPC-olig 7. **D)** Loess fit of the latent abundance of glutamatergic neurons 5 (an L2/3 IT neuron subcluster) across macaques of all ages (upper), and of age > 3 (lower). **E)** GO term and KEGG pathway enrichment of up and down regulated genes of glutamatergic neurons 5 against all other L2/3 IT neuron subclusters. **F)** UMAP showing expression of proinflammatory cytokine response genes.

**Figure S7.**
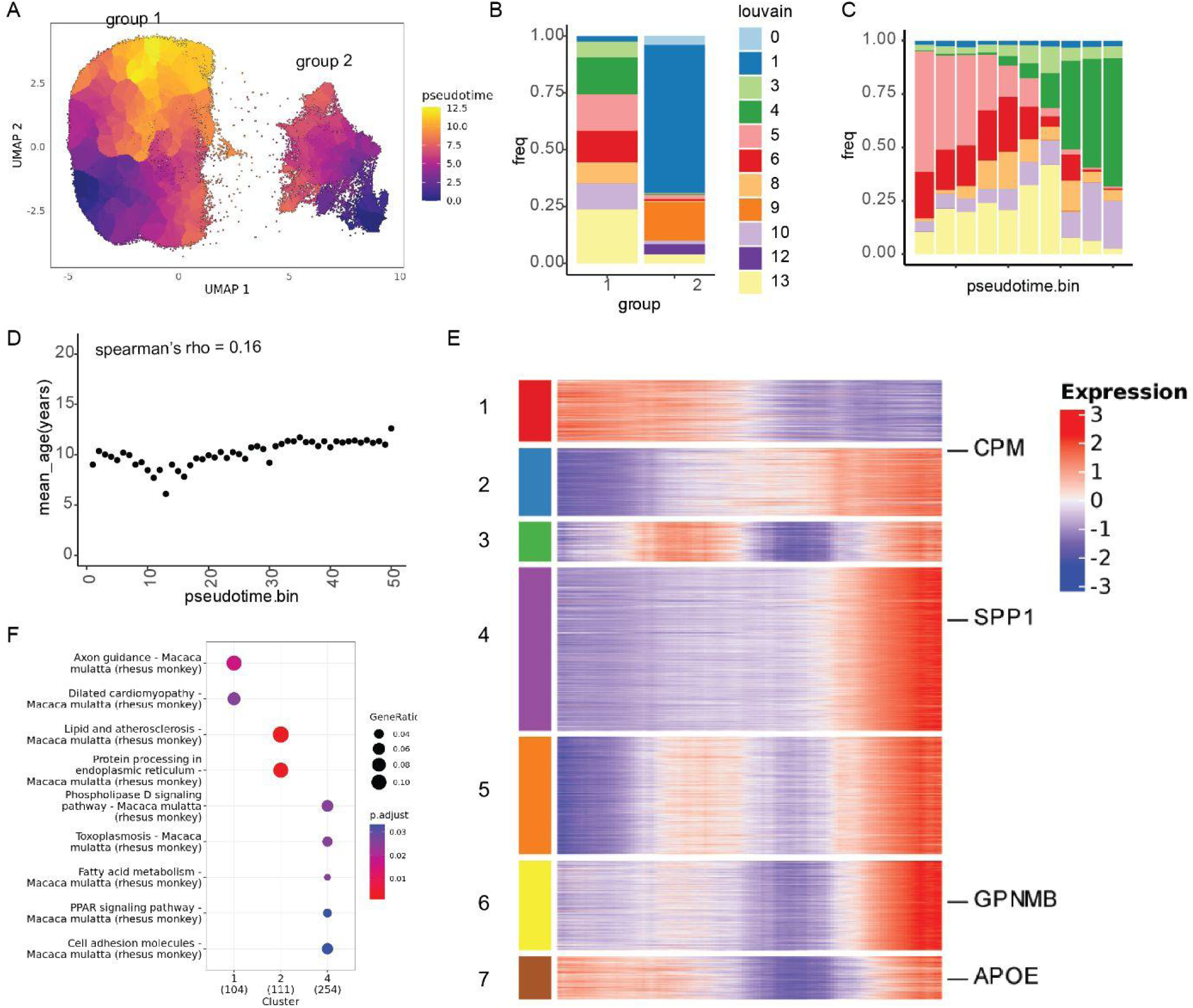
Pseudotime trajectory of age-related transition of microglial state. **A)** UMAP colored by pseudotime output from Monocle3. **B)** Proportion of cells from each subcluster in the graph partitions identified by Monocle 3. **C)** Proportion of cells in group 1 from each subcluster in each pseudotime bin (10 bins). **D)** Dotplot showing correlation of pseudotime bins (50 bins) and mean age in each pseudotime bin for partition 1. **E)** Gene expression heatmap across pseudotime. Expression values are averaged within defined pseudotime bins (50 bins). Expression patterns are clustered into 7 clusters by k-means clustering. **F)** GO term enrichment of genes in each expression trajectory clusters.

**Figure S8.**
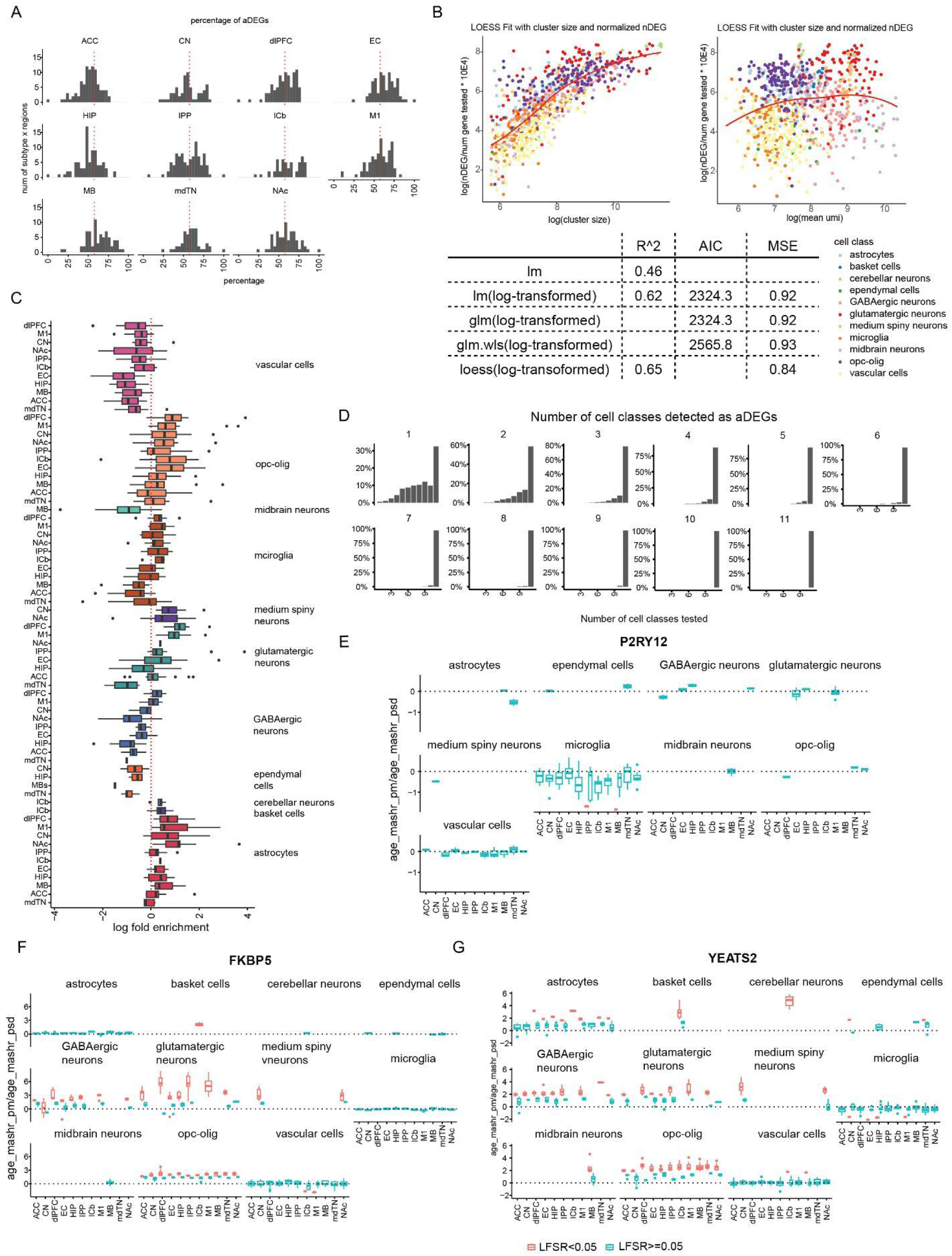
Transcriptional differences and GSEA pathway enrichments of aDEGs. **A)** Percentage of upregulated aDEGs in each combination of subcluster and region. **B)** Loess fit of normalized nDEGs (total number of DEG/number of tests) and cluster size (left) and number of UMI (right). Goodness-of-fit metrics for different models. **C)** Residuals from loess model for each subcluster and region combinations. **D)** Barplots showing the percentage of genes tested across 1 to 11 regions per cell class, stratified by the number of regions in which they were detected as significant aDEGs. Most genes were tested in all 11 regions.E,F. Boxplot showing standardized effect size for P2RY12 **(E),** FKBP5 **(F)** and YEATS2**(G)** showing significant association with age (LFSR<0.05) and coefficients of all other combinations (LFSR>=0.05).

**Figure S9.**
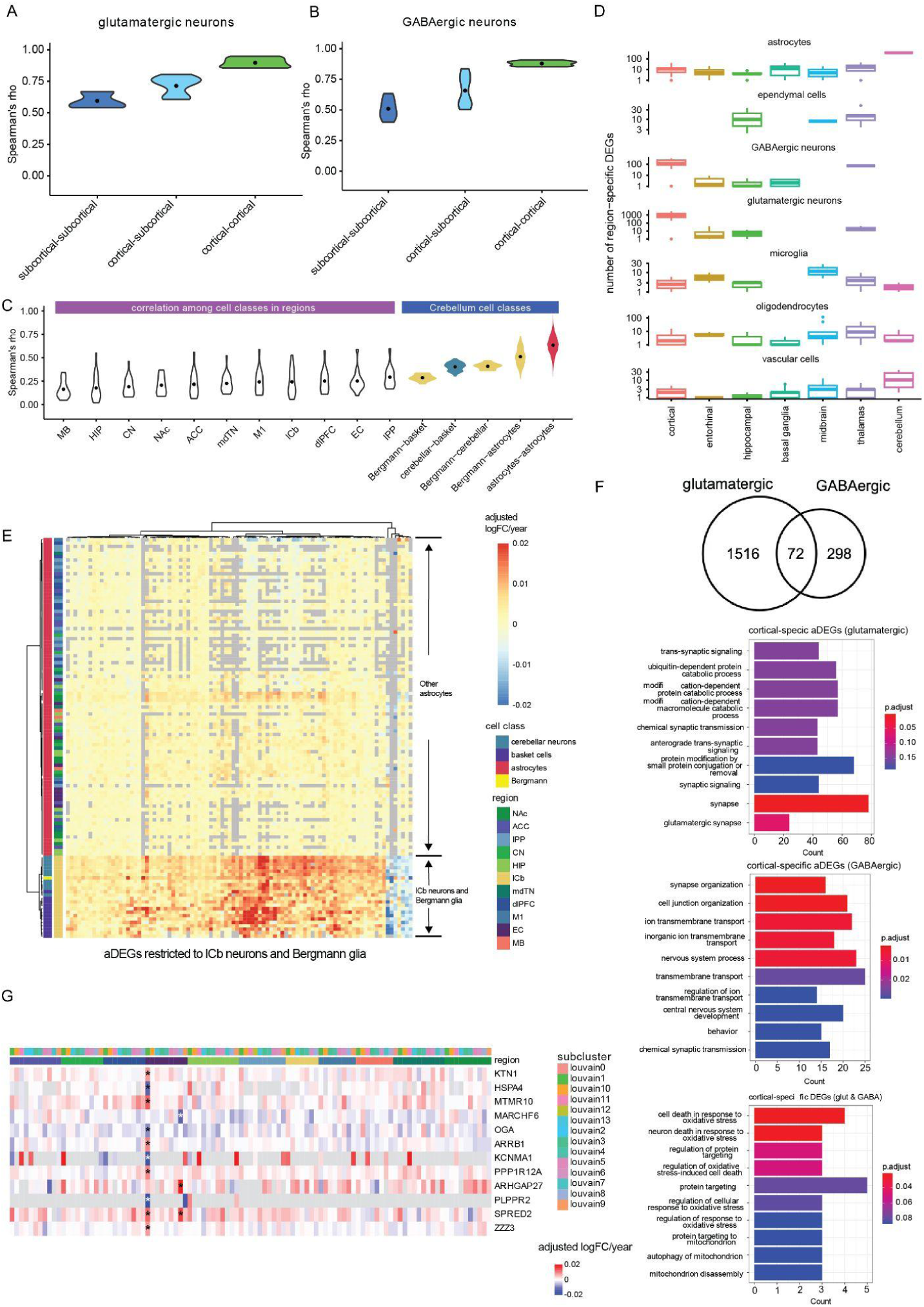
Regional specificity of aDEGs. **A-B**) Violin plots of spearman’s rho of age coefficients of all genes from the linear mixed model after MASHR correction with correlation across these three categories (1) subcortical vs subcortical, (2) subcortical vs cortical, and (3) cortical vs cortical in glutamatergic neurons (A) and GABAergic neurons (B). **C)** Violin plots illustrate spearman’s rho across (1) different cell classes within the same brain region, (2) different pairs of cell classes within cerebellum, and (3) different regions within astrocytes. **D)** Number of region-specific aDEGs (LFSR<0.05), defined as those specifically detected in all regions within each anatomical structure and present in at least 3 out of 4 the cortical structure. Cortical: ACC, dlPFC, M1, IPP; entorhinal: microglia; hippocampal: HIP; basal ganglia: CN, NAc; midbrain: MB; thalamus: mdTN; cerebellum: lCb. **E)** Heatmap showing adjusted logFC (posterior mean after MASHR) per year for aDEGs specific to and shared by cerebellum neurons and Bergmann glia across combinations of subcluster and region. **F)** GO pathways enriched in cortical specific aDEGs in glutamatergic neurons (upper left) or GABAergic neurons (upper right) or both (lower right). Venn plot showing intersection of cortical specific aDEGs in glutamatergic neurons and GABAergic neurons (lower right). **G)** Heatmap showing adjusted logFC (posterior mean after MASHR) per year for microglial aDEGs specific to the entorhinal cortex across combinations of subcluster and region.

**Figure S10.**
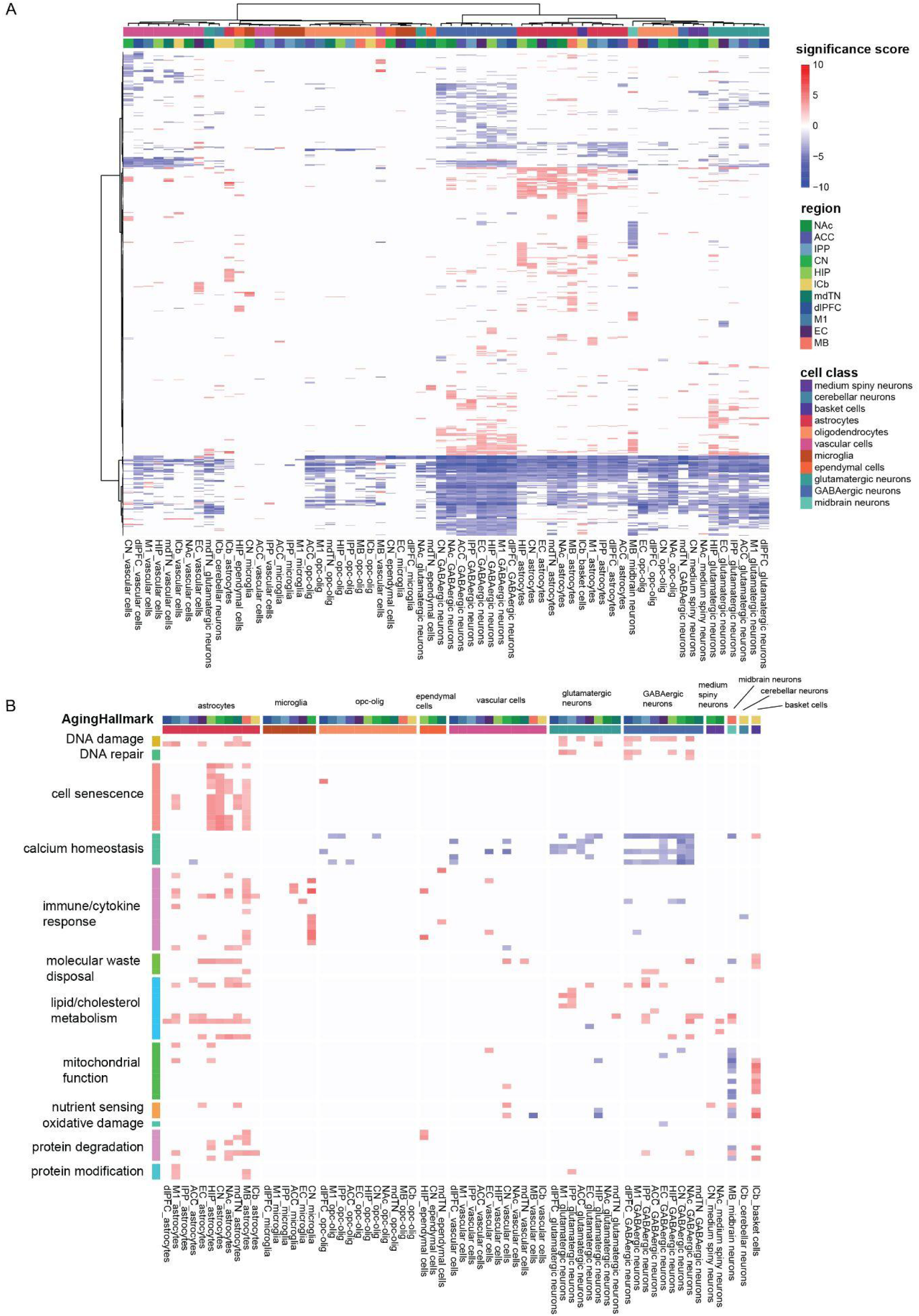
Gene set enrichment analysis of all genes with detected age-related changes. GSEA enrichment of GO pathways of all GO pathways **(A)** and **(B)** on brain aging hallmark sets(*62*) among combinations of subclusters and regions (significance score = - NES sign x log10(p-value)).

**Figure S11.**
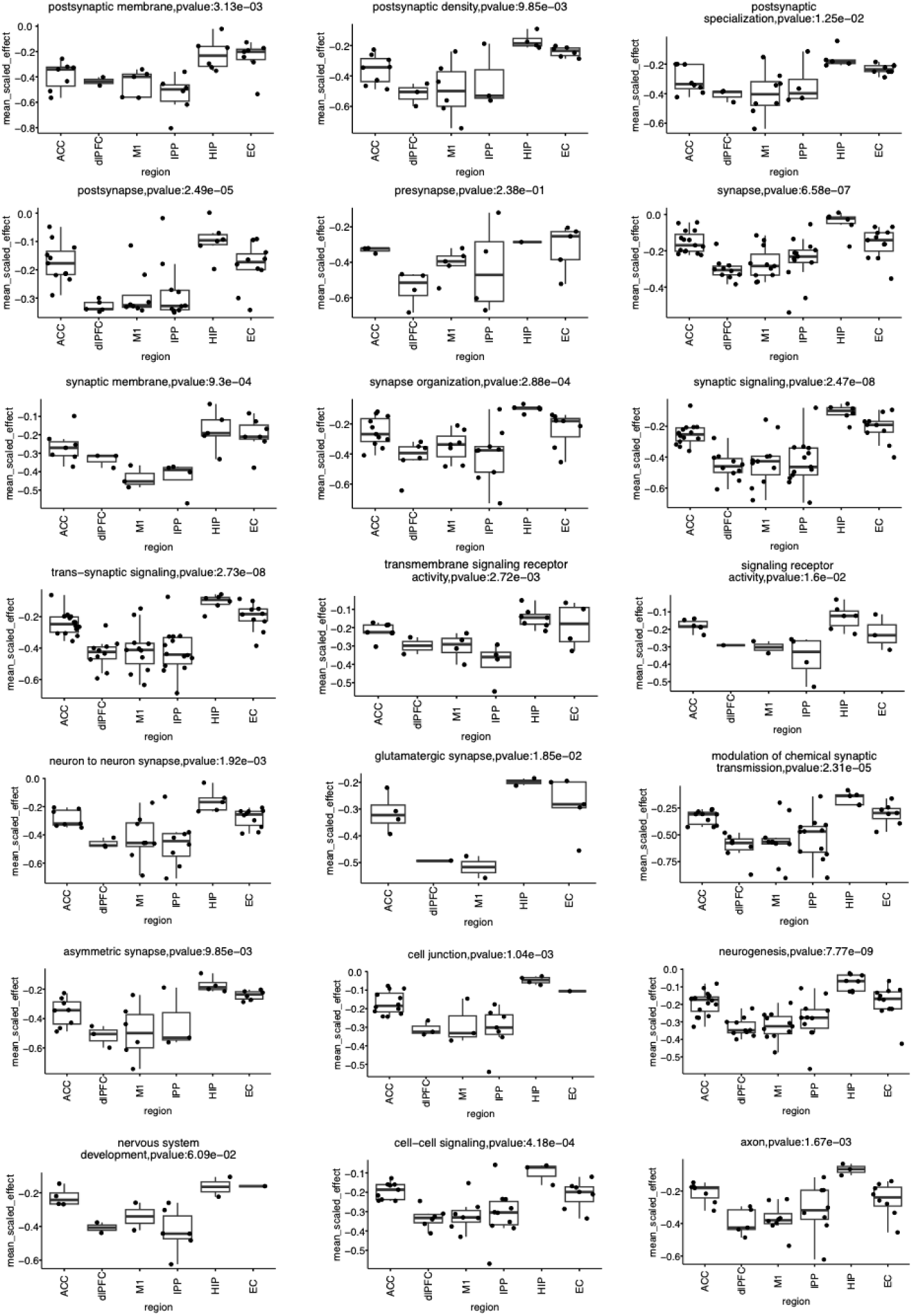
Mean scaled effects of GSEA gene sets on synapse, neural development and cell-cell signaling of glutamatergic neurons. Dot plots showing scaled effects (posterior mean/posterior standard deviation) averaged across genes in a given pathway. Each dot represents a subcluster x region combination with enrichment in cortical regions and the hippocampal formation. For each pathway, an anova model was fit to model the effects of origin of region and external subtype annotation (scaled effect ∼ region + subtype). We show the p-value without correction for region effect for each pathway.

**Figure S12.**
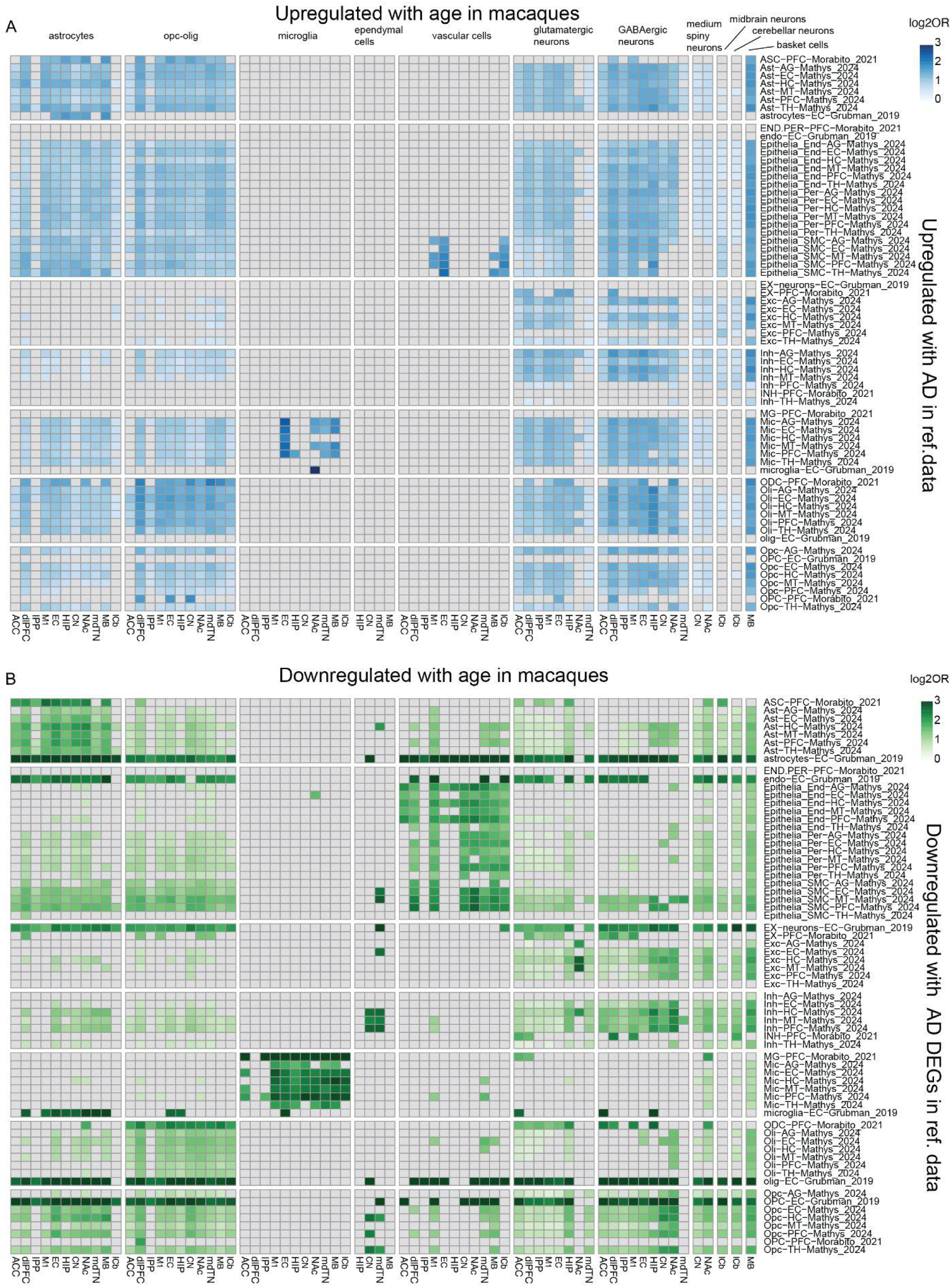
Overlapping gene programs between aDEGs and AD DEGs. Heatmap of log2OR of enrichment performed by a fisher’s exact test of overlaps between **A)** upregulated aDEGs or **B)** downregulated aDEGs and AD DEGs (p adjusted < 0.001).

**Figure S13.**
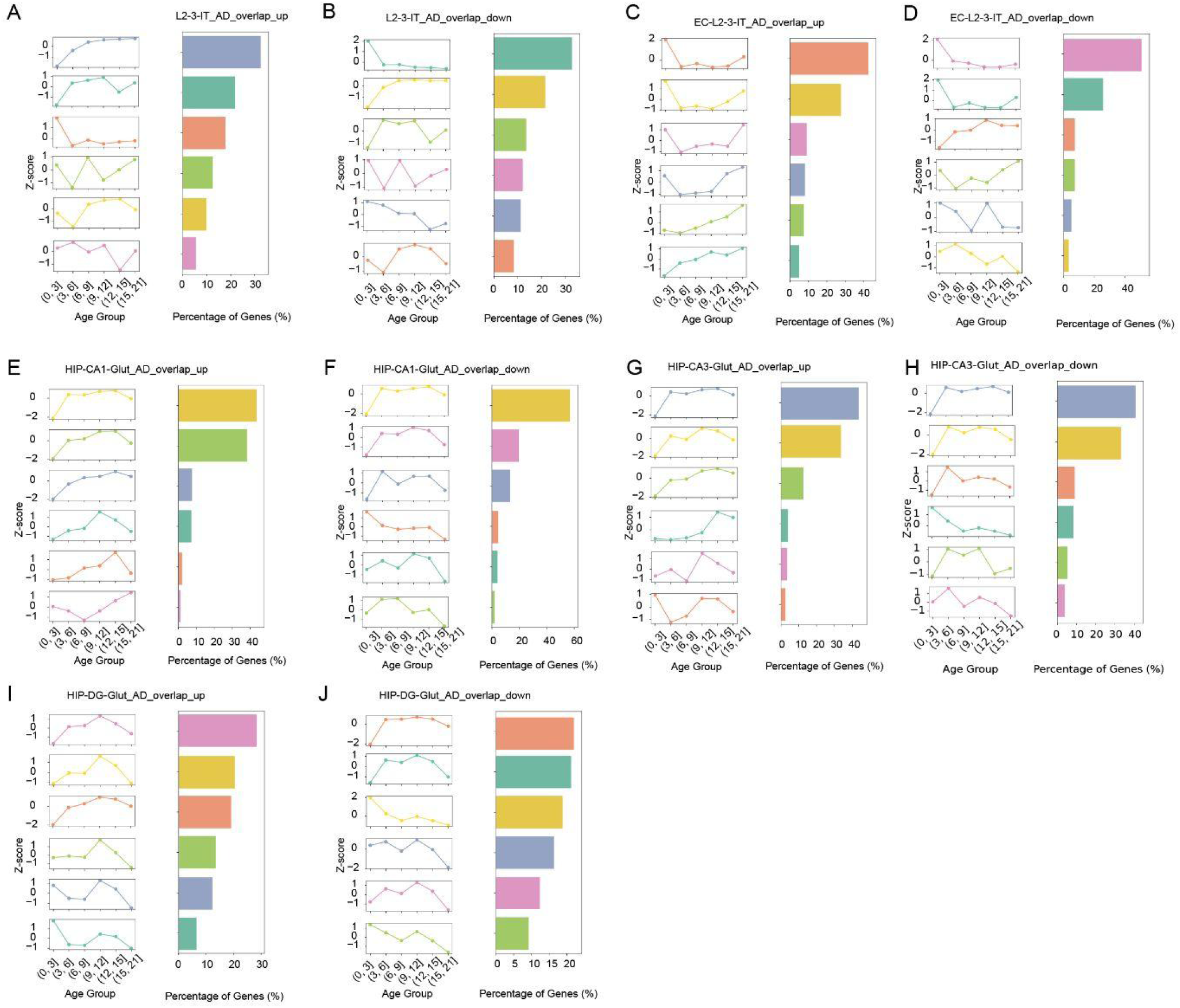
Non-linear temporal dynamics of AD DEGs in aging. Nonlinear temporal trajectory of convergent DEGs (genes overlap between aDEGs and AD DEGs) in each glial cell class or neuronal subtypes. In each panel, from left to right, (1) we showed a lineplot showing average z-score values within each kmeans-cluster and across age bins; (2) a barplot showing proportion of genes bellowing to each k-cluster. The panel orders are **A-B)** neocortical L2/3 IT glutamatergic neurons: upregulated in both aDEG and AD DEG **(A)**, downregulated in both aDEG and AD DEG **(B)**. **C-D)** EC L2/3 IT glutamatergic neurons: upregulated in both aDEG and AD DEG**(C)**, downregulated in both aDEG and AD DEG **(D)**. **E-F)** HIP CA1 glutamatergic neurons: upregulated in both aDEG and AD DEG **(E)**, downregulated in both aDEG and AD DEG **(F)**. **G-H)** HIP CA3 glutamatergic neurons: upregulated in both aDEG and AD DEG **(G)**, downregulated in both aDEG and AD DEG **(H)**. **I-J)** HIP DG glutamatergic neurons: upregulated in both aDEG and AD DEG **(I)**, downregulated in both aDEG and AD DEG **(J)**.

**Figure S14.**
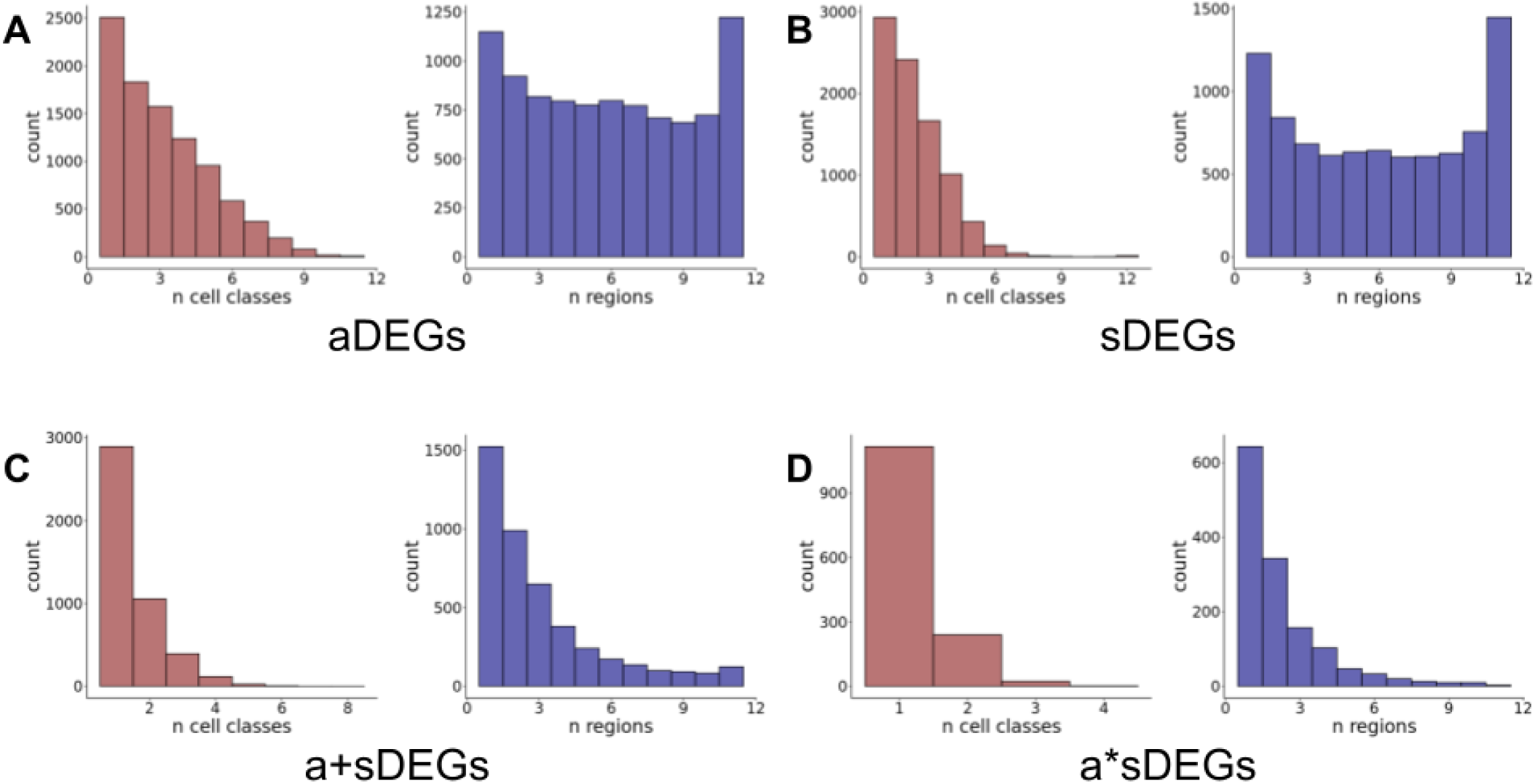
Histograms showing counts of DEGs by number of cell classes (red) or regions (blue) in which they appear.

**Figure S15.**
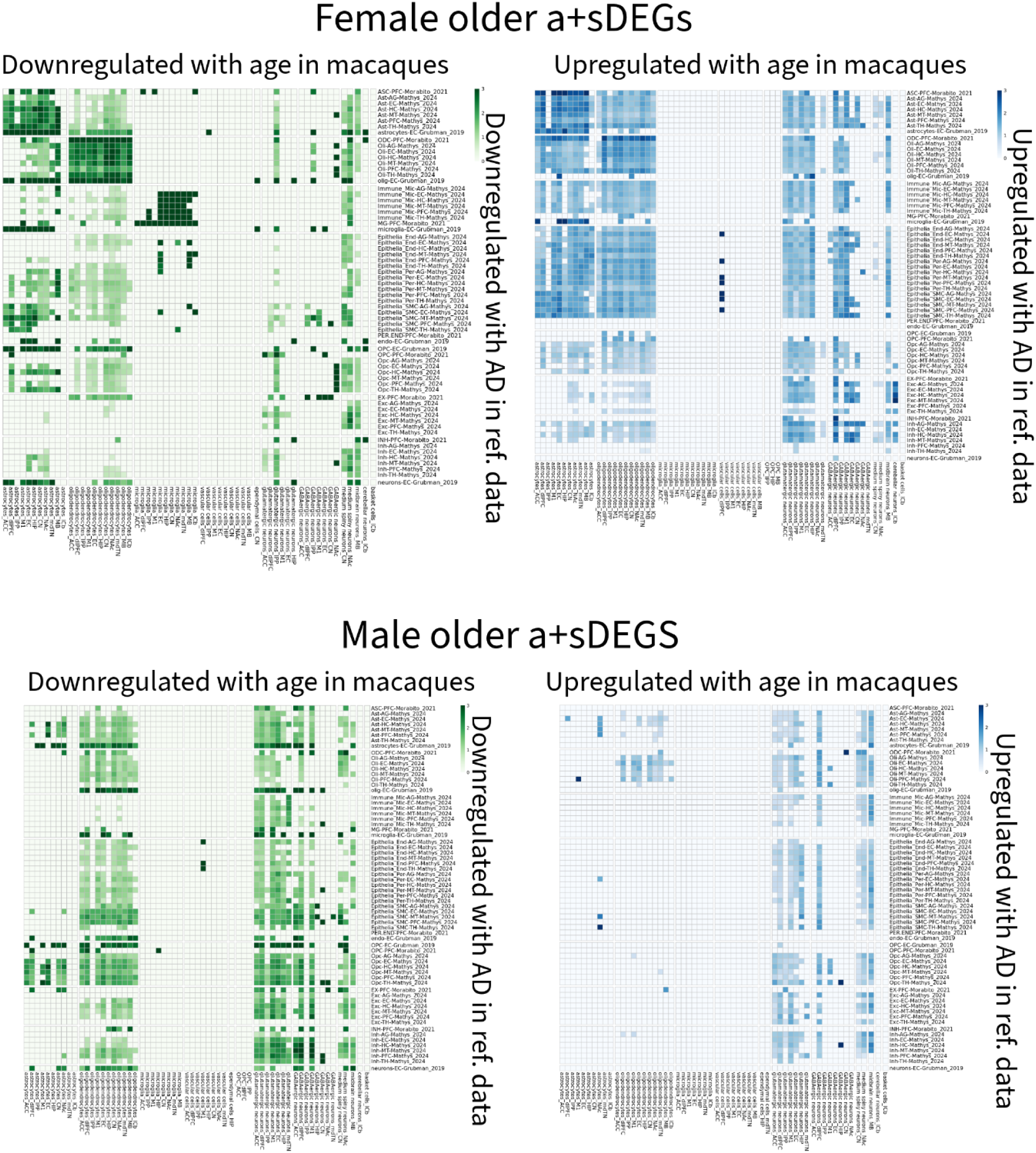
Results of Fisher’s exact tests between a+sDEGs and AD-DEGs. DEGs from this manuscript that were up- or down-regulated with age were tested against AD-DEGs from reference datasets that were up- or down-regulated in AD, respectively. Values shown are log_2_ odds ratios.

## SUPPLEMENTARY NOTES

### Supplementary Note 1

We first used unsupervised Louvain clustering and label transfer from two reference atlases (*13*, *14*) to identify 12 molecularly distinct cell classes across the 11 regions (**Fig 1B**). Based on known cell -type-specific marker genes gene markers and expected regional representation, these cell classes were annotated as either (i) neuronal cells, including glutamatergic neurons (*SLC17A7, SLC17A6*), GABAergic neurons (*GAD1, GAD2*), medium spiny neurons (*DRD1, DRD2*), midbrain neurons (neurons residing in midbrain), cerebellar basket cells (*SORCS3, KIT*) and other cerebellar neurons (granule cells and Purkinje cells*, GRM4*); or (ii) non-neuronal cells, including oligodendrocyte precursor cells (OPCs; *PCDH15*), oligodendrocytes (*MBP, MOG*), astrocytes (*SLC1A2, GFAP*), microglia (*DOCK2, P2RY12*), vascular cells (*CFH*), and ependymal cells (*SPAG17*) (**Methods**; Fig. 1E).

What is the coverage of these major cell classes in our dataset? The major cell classes were captured across the lifespan with no dropouts (Fig. 1C). As expected, the distribution of major cell classes differed extensively across regions, reflecting the cellular makeup underlying known region-specific functions (Fig. 1D; **S1D**) (*13*). Most of the major cell classes recovered are as expected, except for basket cells in midbrain and medium spiny neurons in hippocampus. The presence of these unexpected cell classes likely indicates minor dissection errors into nearby regions: basket cells from cerebellum and medium spiny neurons from the basal ganglia. To account for the proportion of cells from dissection errors, we retain these unexpected cell classes for differential abundance study at the cell class level. Within a region, the expected major cell classes were present in >80% individuals, except for ependymal cells (**Table S2**). Ependymal cells are a rare cell class (0.4% of the full dataset), so their recovery might be more variable in each individual.

### Supplementary Note 2

We first aimed to characterize how different covariates (i.e. region, age, sex) pattern transcriptional variance. To do so, we divided the macaque lifespan to different age groups and created pseudobulk transcriptomes among individuals in each age group for a given region (**Fig S1E**). The pseudobulk transcriptomes were then projected to a principal component analysis (PCA) space. As visualized in the PCA space, transcriptomes were largely separated by regions rather than by age. Therefore, it is important to dissect age-associated change within individual regions. We did observe progressive differences of the transcriptome in each region, with a considerable gap between infancy (age >3 years) and the other age groups (ages 3 - 21). By analyzing the genes most highly correlated with age, we found that the infant samples were transcriptionally distinct from the rest of the donors in the major cell classes (**Fig S1E**). Indeed, the brain tissue volumes of macaques continues to grow throughout infancy, only stabilizing or decelerating at the onset of adolescence, which occurs around 3 or 4 years of age in macaques (*122*, *123*).

We further looked at variance explained by different covariates (i.e. age and sex) within each region. Age explained a small proportion of variance in gene expression (median = 1.1-2.6%), which is consistent with previous studies using bulk RNA-seq (median = 0.6-6.4%) and human brain transcriptomic data from the Genotype–Tissue Expression (GTEx) Consortium V8 dataset (median = 0.3-4.2%) (*8*). Among regions, age explained the most differences in the cerebellum, followed by the primary motor cortex and caudate nucleus (**Fig. S2A**). Sex explained a much smaller proportion of variance, explaining variance of 15-21% of all genes detected (median = 1.4-2.2% among those explained by sex) (**Fig S2B**; **Table S3**).

### Supplementary Note 3

To examine the heterogeneity within each cell class, we then performed unsupervised sub-clustering analysis for each cell class (oligodendrocyte precursor and oligodendrocytes were analyzed together to capture transition states). Overall, we identified a total of 225 subclusters, with 193-195,742 193–195,742 cells in each subcluster (Fig. 1E). Hierarchical clustering of all cell subclusters reflected ontogenic relationships, with two clades of neuronal classes and non-neuronal classes (except OPCs were clustering with neurons potentially due to expression of similar receptors such as *GRID2*). The subclusters show consistent marker gene expression as their corresponding cell class and are defined by a distinct set of marker genes (*e.g.* GABAergic neurons subclusters are marked by *GAD1* and *GAD2*; Figs. 1E, S2C, S3).

The majority of the GABAergic neuron subclusters were observed in cortical and subcortical regions (19 out of 26 subclusters), while some subclusters have regional bias (Fig. 2B). Most of these specializations were reported in a human study (*18*). We identified the four major neocortical types of GABAergic neurons in our dataset (*SST^+^*, *PVALB^+^*, *VIP^+^*, or *LAMP5^+^*), including the *LAMP5^+^LHX6^+^* subtype specific to the primate cortex (*124*) (**Fig. S5B**). A subpopulation of *SST^+^*neurons were enriched in the entorhinal cortex and hippocampus, including an *NPY^+^* subtype reported. A unique GABAergic neuron subtype (*OTX2^+^ GATA3^+^*) was specific to thalamus, which show corresponding markers (*MEIS2*, *FOXP2*) with the counterpart of humans, also enriched in genes involved in neurite outgrowth (*SEMA3C*, *SPON1*), and receptors for serotonin (*HTR2A*), acetylcholine(*CHRM2*, *CHRNA3*), and glutamate (*GRM3*). We also discovered multiple subclusters that are specific to basal ganglia, including LHX6 (*TACR3^+^ CRABP1^+^*), LHX8 (*DLK1^+^ PKHD1L1^+^*), SST(*NPY^+^ CHODL^+^*) and cholinergic (*CHAT^+^ NGFR^+^*) neurons.

We have identified 14-25 subclusters for each non-neuronal cell class and further characterized them by distinct expression profiles, marker genes, and enriched pathways. The oligodendrocyte precursors and oligodendrocytes were partitioned into 14 subclusters. All oligodendrocytes subclusters are found in all regions, showing minimal regional specialization. These subclusters mapped to 4 distinct states along oligodendrocyte differentiation and maturation with differential activities of transcription factor regulons (*125*, *126*): oligodendrocyte precursor cells (olig-OPC 12 and 13; regulons of *PDGFRA, PCDH15*), pre-myelinating oligodendrocytes (olig-OPC 4 and 8, regulons of *ASCL1, SOX5|6|10*), differentiating oligodendrocytes (olig-OPC 6, regulons of *REST*) and myelinating oligodendrocytes (olig-OPC 0, 1, 2, 3, 5, 7, 9, 11, regulons of *MYRF*) (**Figs. S4A, S5C**).

We have observed both microglia and peripheral immune subclusters in our dataset, including border associated macrophage (microglia 11; *DAB2^+^ LYVE1^+^ F13A1^+^*), neutrophils (microglia 2; *FCGR3*+), T lymphocytes (microglia 7; CD3D*^+^* SLAMF*^+^*). Similar to oligodendrocytes, the microglial and immune cells show minimal regional specificity. The microglia subclusters show differential enrichment for signature genes of different microglial states: homeostatic, surveilling, stress response, lipid metabolism, neuroinflammation, etc (**Figs. S4B, S5D**).

The majority of astrocytes subclusters are distributed across the brain while a subset of them (5 out of 16) exhibited region specificity, including lCb-specific Bergmann glia cells (astrocytes 9; *GABRA2^+^*), a subcluster that is specific to the basal ganglia (astrocytes 3, *PENK^+^ SYNDIG1L^+^*), and 3 subclusters (astrocytes 1, 12, 13) enriched in the thalamus. The astrocyte subclusters show differential enrichment for signature genes of homeostatic, reactive, mitophagic, and stress responsive astrocyte states (**Figs. S4C, S5E**).

For rarer cell classes, vascular cells and ependymal cells, we have captured their distinct subtypes in our dataset. Vascular cells subclusters show exclusive enrichment of markers of endothelial cells, pericytes, vascular leptomeningeal cells (VLMC), and vascular smooth muscle cells (VSMCs), most of which distribute broadly across the brain except subcluster 10 (*HTR2C^+^ LMX1A^+^*) was specific to neurons of the midbrain and cerebellum (**Fig. S4D**). Ependymal subclusters show more regional variations, with bias for hippocampus, basal ganglia, midbrain and thalamus, respectively. This potentially reflects distinct molecular signatures of different ventricle structures present in different regions (**Fig. S4E**).

### Supplementary Note 4

We observed a 57.9-fold increase in the number of sex differentially expressed genes (sDEGs) associated with our use of MASH. This magnitude of increase was approximately four times greater than the median fold increase across cell class (14.4-fold) and nearly two times greater than that of the next highest increase (30.9-fold in GABAergic neurons). We hypothesized that the high degree of power in oligodendrocytes could reflect an unusually high degree of covariance among sex-associated effects across many genes in oligodendrocytes. We further hypothesized that this high degree of sharing may be related to fundamental gene regulatory responses to sex hormones that are broadly recapitulated across all or many oligodendrocyte subtypes.

To evaluate our hypothesis, we tested the prediction that sDEGs that are highly shared across oligodendrocytes would be enriched for genes known to be differentially expressed in response to estrogen and androgens. To do so, we first downloaded the androgen response (M5908), estrogen response early (M5906), and estrogen response late (M5907) gene sets from the Hallmark dataset from MSigDB. We then used ENSEMBL orthology information downloaded through biomaRt to identify all one-to-one orthologous macaque genes, which we combined into a single non redundant gene set comprising 360 macaque orthologs.

We defined highly shared sDEGs as those passing a threshold of LFSR < 0.01 in at least 50% of region/Louvain clusters in oligodendrocytes. We identified 113 genes that were highly shared in oligodendrocyte region/Louvain clusters by these thresholds. We then used Fisher’s exact test to test the prediction that genes in the sex hormone response gene set are overrepresented among highly shared oligodendrocyte sDEGs. We found that, indeed, sex hormone response genes were 4.43x more likely to be among highly shared oligodendrocyte sDEGs relative to chance (P = 0.0002). We found that this result was broadly robust to differing LFSR and sharedness thresholds.

To evaluate whether this pattern was specific to oligodendrocytes, we repeated our enrichment analysis across all other cell classes. This analysis was complicated by the differing number of shared genes across cell classes and our use of thresholding. Therefore, we used an iterative strategy to relax our thresholds in an attempt to equalize the number of genes considered shared across cell classes. First, we implemented the same thresholds as we used for oligodendrocytes (LFSR < 0.01 in >= 50% of region/Louvain clusters). If the number of genes was less than 100, we iteratively reduced the percentage by 1% in a stepwise fashion until either the number of passing genes surpassed 100 or the percentage of Louvain/clusters reached 10%. If the number of passing genes still did not exceed 100, we repeated the iterative process with a relaxed threshold of 0.05. If the number of passing genes still did not surpass 100, we continued to relax the LFSR threshold in 0.05 increments, each time repeating the iterative process of relaxing the sharedness threshold from a starting point of 50%. The maximum LFSR threshold reached was 0.3 (OPCs).

After roughly equalizing the number of broadly shared genes across cell classes, we tested for enrichment of sex hormone response genes using Fisher’s exact tests and adjusted for multiple hypothesis testing using a Benjamini-Hochberg correction. We found that, of all cell classes, only oligodendrocytes (Padj = 0.0025) and vascular cells (Padj = 0.0152) showed enrichment of sex steroid hormone response genes. Results are shown in **Table S9**.

## References

1. E. Colita, V. O. Mateescu, D.-G. Olaru, A. Popa-Wagner, Cognitive decline in ageing and disease: Risk factors, genetics and treatments. Curr. Health Sci. J. 50, 170–180 (2024).

2. V. A. Pandya, R. Patani, Region-specific vulnerability in neurodegeneration: lessons from normal ageing. Ageing Res. Rev. 67, 101311 (2021).

3. H. Fu, J. Hardy, K. E. Duff, Selective vulnerability in neurodegenerative diseases. Nat. Neurosci. 21, 1350–1358 (2018).

4. M. T. Ferretti, A. S. Dimech, A. S. Chadha, Sex and Gender Differences in Alzheimer’s Disease (Academic Press, 2021).

5. K. L. Chiou, M. J. Montague, E. A. Goldman, M. M. Watowich, S. N. Sams, J. Song, J. E. Horvath, K. N. Sterner, A. V. Ruiz-Lambides, M. I. Martínez, J. P. Higham, L. J. N. Brent, M. L. Platt, N. Snyder-Mackler, Rhesus macaques as a tractable physiological model of human ageing. Philos Trans R Soc Lond B Biol Sci 375, 20190612 (2020).

6. A. S. Fröhlich, N. Gerstner, M. Gagliardi, M. Ködel, N. Yusupov, N. Matosin, D. Czamara, S. Sauer, S. Roeh, V. Murek, C. Chatzinakos, N. P. Daskalakis, J. Knauer-Arloth, M. J. Ziller, E. B. Binder, Single-nucleus transcriptomic profiling of human orbitofrontal cortex reveals convergent effects of aging and psychiatric disease. Nat. Neurosci. 27, 2021–2032 (2024).

7. D. Dumitriu, J. Hao, Y. Hara, J. Kaufmann, W. G. M. Janssen, W. Lou, P. R. Rapp, J. H. Morrison, Selective changes in thin spine density and morphology in monkey prefrontal cortex correlate with aging-related cognitive impairment. J. Neurosci. 30, 7507–7515 (2010).

8. K. L. Chiou, A. R. DeCasien, K. P. Rees, C. Testard, C. H. Spurrell, A. A. Gogate, H. A. Pliner, S. Tremblay, A. Mercer, C. J. Whalen, J. E. Negrón-Del Valle, M. C. Janiak, S. E. Bauman Surratt, O. González, N. R. Compo, M. K. Stock, A. V. Ruiz-Lambides, M. I. Martínez, Cayo Biobank Research Unit, M. A. Wilson, A. D. Melin, S. C. Antón, C. S. Walker, J. Sallet, J. M. Newbern, L. M. Starita, J. Shendure, J. P. Higham, L. J. N. Brent, M. J. Montague, M. L. Platt, N. Snyder-Mackler, Multiregion transcriptomic profiling of the primate brain reveals signatures of aging and the social environment. Nat. Neurosci. 25, 1714–1723 (2022).

9. R. Hernandez-Pacheco, D. L. Delgado, R. G. Rawlins, M. J. Kessler, A. V. Ruiz-Lambides, E. Maldonado, A. M. Sabat, Managing the Cayo Santiago rhesus macaque population: The role of density. Am. J. Primatol. 78, 167–181 (2016).

10. M. J. Kessler, R. G. Rawlins, A 75-Year Pictorial History of the Cayo Santiago Rhesus Monkey Colony. American journal of primatology 78, 6 (2015).

11. B. Thierry, M. Singh, W. Kaumanns, Macaque Societies: A Model for the Study of Social Organization (Cambridge University Press, 2004).

12. B. K. Martin, C. Qiu, E. Nichols, M. Phung, R. Green-Gladden, S. Srivatsan, R. Blecher-Gonen, B. J. Beliveau, C. Trapnell, J. Cao, J. Shendure, Optimized single-nucleus transcriptional profiling by combinatorial indexing. Nat. Protoc. 18 (2023).

13. K. L. Chiou, X. Huang, M. O. Bohlen, S. Tremblay, A. R. DeCasien, D. R. O’Day, C. H. Spurrell, A. A. Gogate, T. M. Zintel, Cayo Biobank Research Unit, M. G. Andrews, M. I. Martínez, L. M. Starita, M. J. Montague, M. L. Platt, J. Shendure, N. Snyder-Mackler, A single-cell multi-omic atlas spanning the adult rhesus macaque brain. Science Advances, doi: 10.1126/sciadv.adh1914 (2023).

14. K. Siletti, R. Hodge, A. Mossi Albiach, K. W. Lee, S.-L. Ding, L. Hu, P. Lönnerberg, T. Bakken, T. Casper, M. Clark, N. Dee, J. Gloe, D. Hirschstein, N. V. Shapovalova, C. D. Keene, J. Nyhus, H. Tung, A. M. Yanny, E. Arenas, E. S. Lein, S. Linnarsson, Transcriptomic diversity of cell types across the adult human brain. Science 382, eadd7046 (2023).

15. M. I. Gabitto, K. J. Travaglini, V. M. Rachleff, E. S. Kaplan, B. Long, J. Ariza, Y. Ding, J. T. Mahoney, N. Dee, J. Goldy, E. J. Melief, A. Agrawal, O. Kana, X. Zhen, S. T. Barlow, K. Brouner, J. Campos, J. Campos, A. J. Carr, T. Casper, R. Chakrabarty, M. Clark, J. Cool, R. Dalley, M. Darvas, S.-L. Ding, T. Dolbeare, T. Egdorf, L. Esposito, R. Ferrer, L. E. Fleckenstein, R. Gala, A. Gary, E. Gelfand, J. Gloe, N. Guilford, J. Guzman, D. Hirschstein, W. Ho, M. Hupp, T. Jarsky, N. Johansen, B. E. Kalmbach, L. M. Keene, S. Khawand, M. D. Kilgore, A. Kirkland, M. Kunst, B. R. Lee, M. Leytze, C. L. Mac Donald, J. Malone, Z. Maltzer, N. Martin, R. McCue, D. McMillen, G. Mena, E. Meyerdierks, K. P. Meyers, T. Mollenkopf, M. Montine, A. L. Nolan, J. K. Nyhus, P. A. Olsen, M. Pacleb, C. M. Pagan, N. Peña, T. Pham, C. A. Pom, N. Postupna, C. Rimorin, A. Ruiz, G. A. Saldi, A. M. Schantz, N. V. Shapovalova, S. A. Sorensen, B. Staats, M. Sullivan, S. M. Sunkin, C. Thompson, M. Tieu, J. T. Ting, A. Torkelson, T. Tran, N. J. Valera Cuevas, S. Walling-Bell, M.-Q. Wang, J. Waters, A. M. Wilson, M. Xiao, D. Haynor, N. M. Gatto, S. Jayadev, S. Mufti, L. Ng, S. Mukherjee, P. K. Crane, C. S. Latimer, B. P. Levi, K. A. Smith, J. L. Close, J. A. Miller, R. D. Hodge, E. B. Larson, T. J. Grabowski, M. Hawrylycz, C. D. Keene, E. S. Lein, Integrated multimodal cell atlas of Alzheimer’s disease. Nat Neurosci, doi: 10.1038/s41593-024-01774-5 (2024).

16. Z. Yao, C. T. J. van Velthoven, M. Kunst, M. Zhang, D. McMillen, C. Lee, W. Jung, J. Goldy, A. Abdelhak, M. Aitken, K. Baker, P. Baker, E. Barkan, D. Bertagnolli, A. Bhandiwad, C. Bielstein, P. Bishwakarma, J. Campos, D. Carey, T. Casper, A. B. Chakka, R. Chakrabarty, S. Chavan, M. Chen, M. Clark, J. Close, K. Crichton, S. Daniel, P. DiValentin, T. Dolbeare, L. Ellingwood, E. Fiabane, T. Fliss, J. Gee, J. Gerstenberger, A. Glandon, J. Gloe, J. Gould, J. Gray, N. Guilford, J. Guzman, D. Hirschstein, W. Ho, M. Hooper, M. Huang, M. Hupp, K. Jin, M. Kroll, K. Lathia, A. Leon, S. Li, B. Long, Z. Madigan, J. Malloy, J. Malone, Z. Maltzer, N. Martin, R. McCue, R. McGinty, N. Mei, J. Melchor, E. Meyerdierks, T. Mollenkopf, S. Moonsman, T. N. Nguyen, S. Otto, T. Pham, C. Rimorin, A. Ruiz, R. Sanchez, L. Sawyer, N. Shapovalova, N. Shepard, C. Slaughterbeck, J. Sulc, M. Tieu, A. Torkelson, H. Tung, N. Valera Cuevas, S. Vance, K. Wadhwani, K. Ward, B. Levi, C. Farrell, R. Young, B. Staats, M.-Q. M. Wang, C. L. Thompson, S. Mufti, C. M. Pagan, L. Kruse, N. Dee, S. M. Sunkin, L. Esposito, M. J. Hawrylycz, J. Waters, L. Ng, K. Smith, B. Tasic, X. Zhuang, H. Zeng, A high-resolution transcriptomic and spatial atlas of cell types in the whole mouse brain. Nature 624, 317–332 (2023).

17. GTEx Consortium, The GTEx Consortium atlas of genetic regulatory effects across human tissues. Science 369, 1318–1330 (2020).

18. H. Mathys, C. A. Boix, L. A. Akay, Z. Xia, J. Davila-Velderrain, A. P. Ng, X. Jiang, G. Abdelhady, K. Galani, J. Mantero, N. Band, B. T. James, S. Babu, F. Galiana-Melendez, K. Louderback, D. Prokopenko, R. E. Tanzi, D. A. Bennett, L.-H. Tsai, M. Kellis, Single-cell multiregion dissection of Alzheimer’s disease. Nature 632, 858–868 (2024).

19. MapMyCells. https://portal.brain-map.org/atlases-and-data/bkp/mapmycells.

20. Ao Chen, Yidi Sun, Ying Lei, Chao Li, Sha Liao, Juan Meng, Yiqin Bai, Zhen Liu, Zhifeng Liang, Zhiyong Zhu, Nini Yuan, Hao Yang, Zihan Wu, Feng Lin, Kexin Wang, Mei Li, Shuzhen Zhang, Meisong Yang, Tianyi Fei, Zhenkun Zhuang, Yiming Huang, Yong Zhang, Yuanfang Xu, Luman Cui, Ruiyi Zhang, Lei Han, Xing Sun, Bichao Chen, Wenjiao Li, Baoqian Huangfu, Kailong Ma, Jianyun Ma, Zhao Li, Yikun Lin, He Wang, Yanqing Zhong, Huifang Zhang, Qian Yu, Yaqian Wang, Xing Liu, Jian Peng, Chuanyu Liu, Wei Chen, Wentao Pan, Yingjie An, Shihui Xia, Yanbing Lu, Mingli Wang, Xinxiang Song, Shuai Liu, Zhifeng Wang, Chun Gong, Xin Huang, Yue Yuan, Yun Zhao, Qinwen Chai, Xing Tan, Jianfeng Liu, Mingyuan Zheng, Shengkang Li, Yaling Huang, Yan Hong, Zirui Huang, Min Li, Mengmeng Jin, Yan Li, Hui Zhang, Suhong Sun, Li Gao, Yinqi Bai, Mengnan Cheng, Guohai Hu, Shiping Liu, Bo Wang, Bin Xiang, Shuting Li, Huanhuan Li, Mengni Chen, Shiwen Wang, Minglong Li, Weibin Liu, Xin Liu, Qian Zhao, Michael Lisby, Jing Wang, Jiao Fang, Yun Lin, Qing Xie, Zhen Liu, Jie He, Huatai Xu, Wei Huang, Jan Mulder, Huanming Yang, Yangang Sun, Mathias Uhlen, Muming Poo, Jian Wang, Jianhua Yao, Wu Wei, Yuxiang Li, Zhiming Shen, Longqi Liu, Zhiyong Liu, Xun Xu, Chengyu Li, Single-cell spatial transcriptome reveals cell-type organization in the macaque cortex. Cell 186, 3726–3743.e24 (2023).

21. S. Hao, X. Zhu, Z. Huang, Q. Yang, H. Liu, Y. Wu, Y. Zhan, Y. Dong, C. Li, H. Wang, E. Haasdijk, Z. Wu, S. Li, H. Yan, L. Zhu, S. Guo, Z. Wang, A. Ye, Y. Lin, L. Cui, X. Tan, H. Liu, M. Wang, J. Chen, Y. Zhong, W. Du, G. Wang, T. Lai, M. Cao, T. Yang, Y. Xu, L. Li, Q. Yu, Z. Zhuang, Y. Xia, Y. Lei, Y. An, M. Cheng, Y. Zhao, L. Han, Y. Yuan, X. Song, Y. Song, L. Gu, C. Liu, X. Lin, R. Wang, Z. Wang, Y. Wang, S. Li, H. Li, J. Song, M. Chen, W. Zhou, N. Yuan, S. Sun, S. Wang, Y. Chen, M. Zheng, J. Fang, R. Zhang, S. Zhang, Q. Chai, J. Liu, W. Wei, J. He, H. Zhou, Y. Sun, Z. Liu, C. Liu, J. Yao, Z. Liang, X. Xu, M. Poo, C. Li, C. I. De Zeeuw, Z. Shen, Z. Liu, L. Liu, S. Liu, Y. Sun, C. Liu, Cross-species single-cell spatial transcriptomic atlases of the cerebellar cortex. Science 385, eado3927 (2024).

22. G. S. Green, M. Fujita, H.-S. Yang, M. Taga, A. Cain, C. McCabe, N. Comandante-Lou, C. C. White, A. K. Schmidtner, L. Zeng, A. Sigalov, Y. Wang, A. Regev, H.-U. Klein, V. Menon, D. A. Bennett, N. Habib, P. L. De Jager, Cellular communities reveal trajectories of brain ageing and Alzheimer’s disease. Nature 633, 634–645 (2024).

23. Website.

24. M. A. Nascimento, S. Biagiotti, V. Herranz-Pérez, S. Santiago, R. Bueno, C. J. Ye, T. J. Abel, Z. Zhang, J. S. Rubio-Moll, A. R. Kriegstein, Z. Yang, J. M. Garcia-Verdugo, E. J. Huang, A. Alvarez-Buylla, S. F. Sorrells, Protracted neuronal recruitment in the temporal lobes of young children. Nature 626, 1056–1065 (2024).

25. M. Duran, E. R. Barkan, A. Tresenrider, H. Lee, R. Z. Friedman, N. Lammers, M. Colon, J. Franks, B. Ewing, D. Kimelman, C. Trapnell, A statistical framework for inferring genetic requirements from embryo-scale single-cell sequencing experiments, bioRxiv (2025)p. 2025.04.03.646654.

26. Luis E.B. Bettio, Luckshi Rajendran, Joana Gil-Mohapel, The effects of aging in the hippocampus and cognitive decline. Neuroscience & Biobehavioral Reviews 79, 66–86 (2017).

27. E. C. Sams, Oligodendrocytes in the aging brain. Neuronal Signaling 5 (2021).

28. J. Luebke, H. Barbas, A. Peters, Effects of normal aging on prefrontal area 46 in the rhesus monkey. Brain Res. Rev. 62, 212–232 (2010).

29. S. M. Urbut, G. Wang, P. Carbonetto, M. Stephens, Flexible statistical methods for estimating and testing effects in genomic studies with multiple conditions. Nat Genet 51, 187–195 (2019).

30. S. M. Urbut, S. Koyama, W. Hornsby, R. Bhukar, S. Kheterpal, B. Truong, M. S. Selvaraj, B. Neale, C. J. O’Donnell, G. M. Peloso, P. Natarajan, Bayesian multivariate genetic analysis improves translational insights. iScience 26, 107854 (2023).

31. W. E. Allen, T. R. Blosser, Z. A. Sullivan, C. Dulac, X. Zhuang, Molecular and spatial signatures of mouse brain aging at single-cell resolution. Cell 186, 194–208.e18 (2023).

32. S. Sun, Y. Sun, S.-C. Ling, L. Ferraiuolo, M. McAlonis-Downes, Y. Zou, K. Drenner, Y. Wang, D. Ditsworth, S. Tokunaga, A. Kopelevich, B. K. Kaspar, C. Lagier-Tourenne, D. W. Cleveland, Translational profiling identifies a cascade of damage initiated in motor neurons and spreading to glia in mutant SOD1-mediated ALS. Proceedings of the National Academy of Sciences 112, E6993–E7002 (2015).

33. R. Mancuso, N. Fattorelli, A. Martinez-Muriana, E. Davis, L. Wolfs, J. Van Den Daele, I. Geric, J. Premereur, P. Polanco, B. Bijnens, P. Preman, L. Serneels, S. Poovathingal, S. Balusu, C. Verfaillie, M. Fiers, B. De Strooper, Xenografted human microglia display diverse transcriptomic states in response to Alzheimer’s disease-related amyloid-β pathology. Nature Neuroscience 27, 886–900 (2024).

34. L. J. M. Mastenbroek, S. M. Kooistra, B. J. L. Eggen, J. R. Prins, The role of microglia in early neurodevelopment and the effects of maternal immune activation. Semin Immunopathol 46, 1 (2024).

35. M. S. Haney, R. Pálovics, C. N. Munson, C. Long, P. K. Johansson, O. Yip, W. Dong, E. Rawat, E. West, J. C. M. Schlachetzki, A. Tsai, I. H. Guldner, B. S. Lamichhane, A. Smith, N. Schaum, K. Calcuttawala, A. Shin, Y.-H. Wang, C. Wang, N. Koutsodendris, G. E. Serrano, T. G. Beach, E. M. Reiman, C. K. Glass, M. Abu-Remaileh, A. Enejder, Y. Huang, T. Wyss-Coray, APOE4/4 is linked to damaging lipid droplets in Alzheimer’s disease microglia. Nature 628, 154–161 (2024).

36. C. G. Silva-García, Devo-aging: Intersections between development and aging. GeroScience 45, 2145–2159 (2023).

37. M. Bigliassi, E. Filho, Functional significance of the dorsolateral prefrontal cortex during exhaustive exercise. Biol. Psychol. 175, 108442 (2022).

38. B. Tomasino, M. Gremese, The Cognitive Side of M1. Front Hum Neurosci 10, 298 (2016).

39. M. E. Driscoll, P. C. Bollu, P. Tadi, “Neuroanatomy, Nucleus Caudate” in StatPearls (StatPearls Publishing, Treasure Island (FL), 2023).

40. L. Muzio, A. Viotti, G. Martino, Microglia in Neuroinflammation and Neurodegeneration: From Understanding to Therapy. Front. Neurosci. 15, 742065 (2021).

41. E. B. Binder, D. Salyakina, P. Lichtner, G. M. Wochnik, M. Ising, B. Pütz, S. Papiol, S. Seaman, S. Lucae, M. A. Kohli, T. Nickel, H. E. Künzel, B. Fuchs, M. Majer, A. Pfennig, N. Kern, J. Brunner, S. Modell, T. Baghai, T. Deiml, P. Zill, B. Bondy, R. Rupprecht, T. Messer, O. Köhnlein, H. Dabitz, T. Brückl, N. Müller, H. Pfister, R. Lieb, J. C. Mueller, E. Lõhmussaar, T. M. Strom, T. Bettecken, T. Meitinger, M. Uhr, T. Rein, F. Holsboer, B. Muller-Myhsok, Polymorphisms in FKBP5 are associated with increased recurrence of depressive episodes and rapid response to antidepressant treatment. Nat. Genet. 36, 1319–1325 (2004).

42. D. Sinclair, S. G. Fillman, M. J. Webster, C. S. Weickert, Dysregulation of glucocorticoid receptor co-factors FKBP5, BAG1 and PTGES3 in prefrontal cortex in psychotic illness. Sci. Rep. 3, 3539 (2013).

43. L. J. Blair, B. A. Nordhues, S. E. Hill, K. M. Scaglione, J. C. O’Leary 3rd, S. N. Fontaine, L. Breydo, B. Zhang, P. Li, L. Wang, C. Cotman, H. L. Paulson, M. Muschol, V. N. Uversky, T. Klengel, E. B. Binder, R. Kayed, T. E. Golde, N. Berchtold, C. A. Dickey, Accelerated neurodegeneration through chaperone-mediated oligomerization of tau. J. Clin. Invest. 123, 4158–4169 (2013).

44. N. Matosin, J. Arloth, D. Czamara, K. Z. Edmond, M. Maitra, A. S. Fröhlich, S. Martinelli, D. Kaul, R. Bartlett, A. R. Curry, N. C. Gassen, K. Hafner, N. S. Müller, K. Worf, G. Rehawi, C. Nagy, T. Halldorsdottir, C. Cruceanu, M. Gagliardi, N. Gerstner, M. Ködel, V. Murek, M. J. Ziller, E. Scarr, R. Tao, A. E. Jaffe, T. Arzberger, P. Falkai, J. E. Kleinmann, D. R. Weinberger, N. Mechawar, A. Schmitt, B. Dean, G. Turecki, T. M. Hyde, E. B. Binder, Associations of psychiatric disease and ageing with FKBP5 expression converge on superficial layer neurons of the neocortex. Acta Neuropathol. 145, 439–459 (2023).

45. R. Yeewa, S. Pohsa, T. Yamsri, W. Wongkummool, P. Jantaree, S. Potikanond, W. Nimlamool, V. Shotelersuk, L. Lo Piccolo, S. Jantrapirom, The histone acylation reader ENL/AF9 regulates aging in Drosophila melanogaster. Neurobiol Aging 144, 153–162 (2024).

46. S. Jaye, U. S. Sandau, J. A. Saugstad, Clathrin mediated endocytosis in Alzheimer’s disease: cell type specific involvement in amyloid beta pathology. Front. Aging Neurosci. 16, 1378576 (2024).

47. S. H. Wood, S. van Dam, T. Craig, R. Tacutu, A. O’Toole, B. J. Merry, J. P. de Magalhães, Transcriptome analysis in calorie-restricted rats implicates epigenetic and post-translational mechanisms in neuroprotection and aging. Genome Biol 16, 285 (2015).

48. K. Jin, Z. Yao, C. T. J. van Velthoven, E. S. Kaplan, K. Glattfelder, S. T. Barlow, G. Boyer, D. Carey, T. Casper, A. B. Chakka, R. Chakrabarty, M. Clark, M. Departee, M. Desierto, A. Gary, J. Gloe, J. Goldy, N. Guilford, J. Guzman, D. Hirschstein, C. Lee, E. Liang, T. Pham, M. Reding, K. Ronellenfitch, A. Ruiz, J. Sevigny, N. Shapovalova, L. Shulga, J. Sulc, A. Torkelson, H. Tung, B. Levi, S. M. Sunkin, N. Dee, L. Esposito, K. A. Smith, B. Tasic, H. Zeng, Brain-wide cell-type-specific transcriptomic signatures of healthy ageing in mice. Nature, 1–15 (2025).

49. H. Zhang, J. Li, J. Ren, S. Sun, S. Ma, W. Zhang, Y. Yu, Y. Cai, K. Yan, W. Li, B. Hu, P. Chan, G.-G. Zhao, J. C. I. Belmonte, Q. Zhou, J. Qu, S. Wang, G.-H. Liu, Single-nucleus transcriptomic landscape of primate hippocampal aging. Protein Cell 12, 695–716 (2021).

50. L. C. Graham, M. J. Naldrett, S. G. Kohama, C. Smith, D. J. Lamont, B. W. McColl, T. H. Gillingwater, P. Skehel, H. F. Urbanski, T. M. Wishart, Regional Molecular Mapping of Primate Synapses during Normal Healthy Aging. Cell Rep 27, 1018–1026.e4 (2019).

51. J.-F. Chien, H. Liu, B.-A. Wang, C. Luo, A. Bartlett, R. Castanon, N. D. Johnson, J. R. Nery, J. Osteen, J. Li, J. Altshul, M. Kenworthy, C. Valadon, M. Liem, N. Claffey, C. O’Connor, L. A. Seeker, J. R. Ecker, M. M. Behrens, E. A. Mukamel, Cell-type-specific effects of age and sex on human cortical neurons. Neuron 112, 2524–2539.e5 (2024).

52. A. A. Dillman, E. Majounie, J. Ding, J. R. Gibbs, D. Hernandez, S. Arepalli, B. J. Traynor, A. B. Singleton, D. Galter, M. R. Cookson, Transcriptomic profiling of the human brain reveals that altered synaptic gene expression is associated with chronological aging. Sci Rep 7, 16890 (2017).

53. T. Lu, Y. Pan, S.-Y. Kao, C. Li, I. Kohane, J. Chan, B. A. Yankner, Gene regulation and DNA damage in the ageing human brain. Nature 429, 883–891 (2004).

54. E. Ling, J. Nemesh, M. Goldman, N. Kamitaki, N. Reed, R. E. Handsaker, G. Genovese, J. S. Vogelgsang, S. Gerges, S. Kashin, S. Ghosh, J. M. Esposito, K. Morris, D. Meyer, A. Lutservitz, C. D. Mullally, A. Wysoker, L. Spina, A. Neumann, M. Hogan, K. Ichihara, S. Berretta, S. A. McCarroll, A concerted neuron-astrocyte program declines in ageing and schizophrenia. Nature 627, 604–611 (2024).

55. D. L. Dickstein, D. Kabaso, A. B. Rocher, J. I. Luebke, S. L. Wearne, P. R. Hof, Changes in the structural complexity of the aged brain. Aging Cell 6, 275–284 (2007).

56. A. Peters, C. Sethares, J. I. Luebke, SYNAPSES ARE LOST DURING AGING IN THE PRIMATE PREFRONTAL CORTEX. Neuroscience 152, 970 (2007).

57. M. E. Young, D. T. Ohm, D. Dumitriu, P. R. Rapp, J. H. Morrison, Differential effects of aging on dendritic spines in visual cortex and prefrontal cortex of the rhesus monkey. Neuroscience 274, 33 (2014).

58. J. H. Morrison, M. G. Baxter, The Aging Cortical Synapse: Hallmarks and Implications for Cognitive Decline. Nature reviews. Neuroscience 13, 240 (2012).

59. E. G. Knox, M. R. Aburto, G. Clarke, J. F. Cryan, C. M. O’Driscoll, The blood-brain barrier in aging and neurodegeneration. Molecular Psychiatry 27, 2659–2673 (2022).

60. S. Ham, S.-J. V. Lee, Advances in transcriptome analysis of human brain aging. Experimental & Molecular Medicine 52, 1787–1797 (2020).

61. J. Lee, H.-J. Kim, Normal Aging Induces Changes in the Brain and Neurodegeneration Progress: Review of the Structural, Biochemical, Metabolic, Cellular, and Molecular Changes. Front. Aging Neurosci. 14, 931536 (2022).

62. M. P. Mattson, T. V. Arumugam, Hallmarks of Brain Aging: Adaptive and Pathological Modification by Metabolic States. Cell metabolism 27, 1176 (2018).

63. C. Eroglu, Astrocytes, hidden puppet masters of the brain. Science (2025).

64. K. B. Lefton, Y. Wu, Y. Dai, T. Okuda, Y. Zhang, A. Yen, G. M. Rurak, S. Walsh, R. Manno, B.-E. Myagmar, J. D. Dougherty, V. K. Samineni, P. C. Simpson, T. Papouin, Norepinephrine signals through astrocytes to modulate synapses. Science (2025).

65. K. A. Guttenplan, I. Maxwell, E. Santos, L. A. Borchardt, E. Manzo, L. Abalde-Atristain, R. D. Kim, M. R. Freeman, GPCR signaling gates astrocyte responsiveness to neurotransmitters and control of neuronal activity. Science 388, 763–768 (2025).

66. D. J. Baker, B. G. Childs, M. Durik, M. E. Wijers, C. J. Sieben, J. Zhong, R. A. Saltness, K. B. Jeganathan, G. C. Verzosa, A. Pezeshki, K. Khazaie, J. D. Miller, J. M. van Deursen, Naturally occurring p16(Ink4a)-positive cells shorten healthy lifespan. Nature 530, 184–189 (2016).

67. M. Xu, T. Pirtskhalava, J. N. Farr, B. M. Weigand, A. K. Palmer, M. M. Weivoda, C. L. Inman, M. B. Ogrodnik, C. M. Hachfeld, D. G. Fraser, J. L. Onken, K. O. Johnson, G. C. Verzosa, L. G. P. Langhi, M. Weigl, N. Giorgadze, N. K. LeBrasseur, J. D. Miller, D. Jurk, R. J. Singh, D. B. Allison, K. Ejima, G. B. Hubbard, Y. Ikeno, H. Cubro, V. D. Garovic, X. Hou, S. J. Weroha, P. D. Robbins, L. J. Niedernhofer, S. Khosla, T. Tchkonia, J. L. Kirkland, Senolytics improve physical function and increase lifespan in old age. Nat Med 24, 1246–1256 (2018).

68. M. Suda, I. Shimizu, G. Katsuumi, Y. Yoshida, Y. Hayashi, R. Ikegami, N. Matsumoto, Y. Yoshida, R. Mikawa, A. Katayama, J. Wada, M. Seki, Y. Suzuki, A. Iwama, H. Nakagami, A. Nagasawa, R. Morishita, M. Sugimoto, S. Okuda, M. Tsuchida, K. Ozaki, M. Nakanishi-Matsui, T. Minamino, Senolytic vaccination improves normal and pathological age-related phenotypes and increases lifespan in progeroid mice. Nat Aging 1, 1117–1126 (2021).

69. J. Lei, Z. Xin, N. Liu, T. Ning, Y. Jing, Y. Qiao, Z. He, M. Jiang, Y. Yang, Z. Zhang, L. Zhao, J. Li, D. Lv, Y. Yan, H. Zhang, L. Xiao, B. Zhang, H. Huang, S. Sun, F. Zheng, X. Jiang, H. Lu, X. Dong, S. Yue, C. Ma, J. Shuai, Z. Ji, F. Liu, Y. Ye, K. Yan, Q. Hu, G. Xu, Q. Zhao, R. Wu, Y. Cai, Y. Fan, Y. Jing, Q. Wang, P. Reddy, X. Lu, Z. Zheng, B. Liu, A. Haghani, S. Ma, K. Suzuki, C. Rodriguez Esteban, J. Yang, M. Song, S. Horvath, W. Zhang, W. Li, A. P. Xiang, L. Zhu, X. Fu, G. Zhao, J. C. I. Belmonte, J. Qu, S. Wang, G.-H. Liu, Senescence-resistant human mesenchymal progenitor cells counter aging in primates. Cell 0 (2025).

70. A. Grubman, G. Chew, J. F. Ouyang, G. Sun, X. Y. Choo, C. McLean, R. K. Simmons, S. Buckberry, D. B. Vargas-Landin, D. Poppe, J. Pflueger, R. Lister, O. J. L. Rackham, E. Petretto, J. M. Polo, A single-cell atlas of entorhinal cortex from individuals with Alzheimer’s disease reveals cell-type-specific gene expression regulation. Nat Neurosci 22, 2087–2097 (2019).

71. S. Morabito, E. Miyoshi, N. Michael, S. Shahin, A. C. Martini, E. Head, J. Silva, K. Leavy, M. Perez-Rosendahl, V. Swarup, Single-nucleus chromatin accessibility and transcriptomic characterization of Alzheimer’s disease. Nat Genet 53, 1143–1155 (2021).

72. S. Hägg, J. Jylhävä, Sex differences in biological aging with a focus on human studies. Elife 10 (2021).

73. A. M. Bronikowski, R. P. Meisel, P. R. Biga, J. R. Walters, J. E. Mank, E. Larschan, G. S. Wilkinson, N. Valenzuela, A. M. Conard, J. P. de Magalhães, J. E. Duan, A. E. Elias, T. Gamble, R. M. Graze, K. E. Gribble, J. A. Kreiling, N. C. Riddle, Sex-specific aging in animals: Perspective and future directions. Aging Cell 21, e13542 (2022).

74. D. B. Dubal, C. T. Murphy, Y. Suh, B. A. Benayoun, Biological sex matters in brain aging. Neuron 113, 2–6 (2025).

75. N. M. Armstrong, Y. An, L. Beason-Held, J. Doshi, G. Erus, L. Ferrucci, C. Davatzikos, S. M. Resnick, Sex differences in brain aging and predictors of neurodegeneration in cognitively healthy older adults. Neurobiol. Aging 81, 146–156 (2019).

76. J. Yang, J. Qu, H. Ma, Recent developments in understanding brain aging: sex differences, mechanisms, and implications in diseases. OAE Publishing Inc. 2 (2022).

77. N. Johansen, S. Somasundaram, K. J. Travaglini, A. M. Yanny, M. Shumyatcher, T. Casper, C. Cobbs, N. Dee, R. Ellenbogen, M. Ferreira, J. Goldy, J. Guzman, R. Gwinn, D. Hirschstein, N. L. Jorstad, C. D. Keene, A. Ko, B. P. Levi, J. G. Ojemann, T. Pham, N. Shapovalova, D. Silbergeld, J. Sulc, A. Torkelson, H. Tung, K. Smith, E. S. Lein, T. E. Bakken, R. D. Hodge, J. A. Miller, Interindividual variation in human cortical cell type abundance and expression. Science 382, eadf2359 (2023).

78. A. R. DeCasien, P. Auluck, S. Liu, S. R. Marenco, M. Cookson, A. Raznahan, Sex effects on gene expression across the human cerebral cortex at single cell resolution, bioRxiv (2025). 10.1101/2025.06.30.661781.

79. A. Elbaz, J. H. Bower, D. M. Maraganore, S. K. McDonnell, B. J. Peterson, J. E. Ahlskog, D. J. Schaid, W. A. Rocca, Risk tables for parkinsonism and Parkinson’s disease. J. Clin. Epidemiol. 55, 25–31 (2002).

80. F.-M. Zhou, “The striatal medium spiny neurons: what they are and how they link with Parkinson’s disease” in Genetics, Neurology, Behavior, and Diet in Parkinson’s Disease, C. R. Martin, V. R. Preedy, Eds. (Elsevier, 2020), pp. 395–412.

81. M. Rodriguez, C. Rodriguez-Sabate, I. Morales, A. Sanchez, M. Sabate, Parkinson’s disease as a result of aging. Aging Cell 14, 293–308 (2015).

82. H. Wang, D. Devadoss, M. Nair, H. S. Chand, M. K. Lakshmana, Novel Alzheimer risk factor IQ motif containing protein K is abundantly expressed in the brain and is markedly increased in patients with Alzheimer’s disease. Front Cell Neurosci 16, 954071 (2022).

83. A. M. Smith, K. Davey, S. Tsartsalis, C. Khozoie, N. Fancy, S. S. Tang, E. Liaptsi, M. Weinert, A. McGarry, R. C. J. Muirhead, S. Gentleman, D. R. Owen, P. M. Matthews, Diverse human astrocyte and microglial transcriptional responses to Alzheimer’s pathology. Acta Neuropathol 143, 75–91 (2022).

84. B. W. Kunkle, B. Grenier-Boley, R. Sims, J. C. Bis, V. Damotte, A. C. Naj, A. Boland, M. Vronskaya, S. J. van der Lee, A. Amlie-Wolf, C. Bellenguez, A. Frizatti, V. Chouraki, E. R. Martin, K. Sleegers, N. Badarinarayan, J. Jakobsdottir, K. L. Hamilton-Nelson, S. Moreno-Grau, R. Olaso, R. Raybould, Y. Chen, A. B. Kuzma, M. Hiltunen, T. Morgan, S. Ahmad, B. N. Vardarajan, J. Epelbaum, P. Hoffmann, M. Boada, G. W. Beecham, J.-G. Garnier, D. Harold, A. L. Fitzpatrick, O. Valladares, M.-L. Moutet, A. Gerrish, A. V. Smith, L. Qu, D. Bacq, N. Denning, X. Jian, Y. Zhao, M. Del Zompo, N. C. Fox, S.-H. Choi, I. Mateo, J. T. Hughes, H. H. Adams, J. Malamon, F. Sanchez-Garcia, Y. Patel, J. A. Brody, B. A. Dombroski, M. C. D. Naranjo, M. Daniilidou, G. Eiriksdottir, S. Mukherjee, D. Wallon, J. Uphill, T. Aspelund, L. B. Cantwell, F. Garzia, D. Galimberti, E. Hofer, M. Butkiewicz, B. Fin, E. Scarpini, C. Sarnowski, W. S. Bush, S. Meslage, J. Kornhuber, C. C. White, Y. Song, R. C. Barber, S. Engelborghs, S. Sordon, D. Voijnovic, P. M. Adams, R. Vandenberghe, M. Mayhaus, L. A. Cupples, M. S. Albert, P. P. De Deyn, W. Gu, J. J. Himali, D. Beekly, A. Squassina, A. M. Hartmann, A. Orellana, D. Blacker, E. Rodriguez-Rodriguez, S. Lovestone, M. E. Garcia, R. S. Doody, C. Munoz-Fernadez, R. Sussams, H. Lin, T. J. Fairchild, Y. A. Benito, C. Holmes, H. Karamujić-Čomić, M. P. Frosch, H. Thonberg, W. Maier, G. Roshchupkin, B. Ghetti, V. Giedraitis, A. Kawalia, S. Li, R. M. Huebinger, L. Kilander, S. Moebus, I. Hernández, M. I. Kamboh, R. Brundin, J. Turton, Q. Yang, M. J. Katz, L. Concari, J. Lord, A. S. Beiser, C. D. Keene, S. Helisalmi, I. Kloszewska, W. A. Kukull, A. M. Koivisto, A. Lynch, L. Tarraga, E. B. Larson, A. Haapasalo, B. Lawlor, T. H. Mosley, R. B. Lipton, V. Solfrizzi, M. Gill, W. T. Longstreth, T. J. Montine, V. Frisardi, M. Diez-Fairen, F. Rivadeneira, R. C. Petersen, V. Deramecourt, I. Alvarez, F. Salani, A. Ciaramella, E. Boerwinkle, E. M. Reiman, N. Fievet, J. I. Rotter, J. S. Reisch, O. Hanon, C. Cupidi, A. G. Andre Uitterlinden, D. R. Royall, C. Dufouil, R. G. Maletta, I. de Rojas, M. Sano, A. Brice, R. Cecchetti, P. S. George-Hyslop, K. Ritchie, M. Tsolaki, D. W. Tsuang, B. Dubois, D. Craig, C.-K. Wu, H. Soininen, D. Avramidou, R. L. Albin, L. Fratiglioni, A. Germanou, L. Keller, M. Koutroumani, S. E. Arnold, F. Panza, O. Gkatzima, S. Asthana, D. Hannequin, P. Whitehead, C. S. Atwood, P. Caffarra, H. Hampel, I. Quintela, Á. Carracedo, L. Lannfelt, D. C. Rubinsztein, L. L. Barnes, F. Pasquier, L. Frölich, S. Barral, B. McGuinness, T. G. Beach, J. A. Johnston, J. T. Becker, P. Passmore, E. H. Bigio, J. M. Schott, T. D. Bird, J. D. Warren, B. F. Boeve, M. K. Lupton, J. D. Bowen, P. Proitsi, A. Boxer, J. F. Powell, J. R. Burke, J. S. K. Kauwe, J. M. Burns, M. Mancuso, J. Buxbaum, U. Bonuccelli, N. J. Cairns, A. McQuillin, C. Cao, G. Livingston, C. S. Carlson, N. J. Bass, C. M. Carlsson, J. Hardy, R. M. Carney, J. Bras, M. M. Carrasquillo, R. Guerreiro, M. Allen, H. C. Chui, E. Fisher, C. Masullo, E. A. Crocco, C. DeCarli, G. Bisceglio, M. Dick, L. Ma, R. Duara, N. R. Graff-Radford, D. A. Evans, A. Hodges, K. M. Faber, M. Scherer, K. B. Fallon, M. Riemenschneider, D. W. Fardo, R. Heun, M. R. Farlow, H. Kölsch, S. Ferris, M. Leber, T. M. Foroud, I. Heuser, D. R. Galasko, I. Giegling, M. Gearing, M. Hüll, D. H. Geschwind, J. R. Gilbert, J. Morris, R. C. Green, K. Mayo, J. H. Growdon, T. Feulner, R. L. Hamilton, L. E. Harrell, D. Drichel, L. S. Honig, T. D. Cushion, M. J. Huentelman, P. Hollingworth, C. M. Hulette, B. T. Hyman, R. Marshall, G. P. Jarvik, A. Meggy, E. Abner, G. E. Menzies, L.-W. Jin, G. Leonenko, L. M. Real, G. R. Jun, C. T. Baldwin, D. Grozeva, A. Karydas, G. Russo, J. A. Kaye, R. Kim, F. Jessen, N. W. Kowall, B. Vellas, J. H. Kramer, E. Vardy, F. M. LaFerla, K.-H. Jöckel, J. J. Lah, M. Dichgans, J. B. Leverenz, D. Mann, A. I. Levey, S. Pickering-Brown, A. P. Lieberman, N. Klopp, K. L. Lunetta, H.-E. Wichmann, C. G. Lyketsos, K. Morgan, D. C. Marson, K. Brown, F. Martiniuk, C. Medway, D. C. Mash, M. M. Nöthen, E. Masliah, N. M. Hooper, W. C. McCormick, A. Daniele, S. M. McCurry, A. Bayer, A. N. McDavid, J. Gallacher, A. C. McKee, H. van den Bussche, M. Mesulam, C. Brayne, B. L. Miller, S. Riedel-Heller, C. A. Miller, J. W. Miller, A. Al-Chalabi, J. C. Morris, C. E. Shaw, A. J. Myers, J. Wiltfang, S. O’Bryant, J. M. Olichney, V. Alvarez, J. E. Parisi, A. B. Singleton, H. L. Paulson, J. Collinge, W. R. Perry, S. Mead, E. Peskind, D. H. Cribbs, M. Rossor, A. Pierce, N. S. Ryan, W. W. Poon, B. Nacmias, H. Potter, S. Sorbi, J. F. Quinn, E. Sacchinelli, A. Raj, G. Spalletta, M. Raskind, C. Caltagirone, P. Bossù, M. D. Orfei, B. Reisberg, R. Clarke, C. Reitz, A. D. Smith, J. M. Ringman, D. Warden, E. D. Roberson, G. Wilcock, E. Rogaeva, A. C. Bruni, H. J. Rosen, M. Gallo, R. N. Rosenberg, Y. Ben-Shlomo, M. A. Sager, P. Mecocci, A. J. Saykin, P. Pastor, M. L. Cuccaro, J. M. Vance, J. A. Schneider, L. S. Schneider, S. Slifer, W. W. Seeley, A. G. Smith, J. A. Sonnen, S. Spina, R. A. Stern, R. H. Swerdlow, M. Tang, R. E. Tanzi, J. Q. Trojanowski, J. C. Troncoso, V. M. Van Deerlin, L. J. Van Eldik, H. V. Vinters, J. P. Vonsattel, S. Weintraub, K. A. Welsh-Bohmer, K. C. Wilhelmsen, J. Williamson, T. S. Wingo, R. L. Woltjer, C. B. Wright, C.-E. Yu, L. Yu, Y. Saba, A. Pilotto, M. J. Bullido, O. Peters, P. K. Crane, D. Bennett, P. Bosco, E. Coto, V. Boccardi, P. L. De Jager, A. Lleo, N. Warner, O. L. Lopez, M. Ingelsson, P. Deloukas, C. Cruchaga, C. Graff, R. Gwilliam, M. Fornage, A. M. Goate, P. Sanchez-Juan, P. G. Kehoe, N. Amin, N. Ertekin-Taner, C. Berr, S. Debette, S. Love, L. J. Launer, S. G. Younkin, J.-F. Dartigues, C. Corcoran, M. A. Ikram, D. W. Dickson, G. Nicolas, D. Campion, J. Tschanz, H. Schmidt, H. Hakonarson, J. Clarimon, R. Munger, R. Schmidt, L. A. Farrer, C. Van Broeckhoven, M. C. O’Donovan, A. L. DeStefano, L. Jones, J. L. Haines, J.-F. Deleuze, M. J. Owen, V. Gudnason, R. Mayeux, V. Escott-Price, B. M. Psaty, A. Ramirez, L.-S. Wang, A. Ruiz, C. M. van Duijn, P. A. Holmans, S. Seshadri, J. Williams, P. Amouyel, G. D. Schellenberg, J.-C. Lambert, M. A. Pericak-Vance, Genetic meta-analysis of diagnosed Alzheimer’s disease identifies new risk loci and implicates Aβ, tau, immunity and lipid processing. Nature Genetics 51, 414–430 (2019).

85. M. Ximerakis, S. L. Lipnick, B. T. Innes, S. K. Simmons, X. Adiconis, D. Dionne, B. A. Mayweather, L. Nguyen, Z. Niziolek, C. Ozek, V. L. Butty, R. Isserlin, S. M. Buchanan, S. S. Levine, A. Regev, G. D. Bader, J. Z. Levin, L. L. Rubin, Single-cell transcriptomic profiling of the aging mouse brain. Nature neuroscience 22 (2019).

86. L. C. Walker, M. Jucker, The Exceptional Vulnerability of Humans to Alzheimer’s Disease. Trends Mol Med 23, 534–545 (2017).

87. L. Morelli, L. Wei, A. Amorim, J. McDermid, C. R. Abee, B. Frangione, L. C. Walker, E. Levy, Cerebrovascular amyloidosis in squirrel monkeys and rhesus monkeys: apolipoprotein E genotype. FEBS Lett 379, 132–134 (1996).

88. D. Datta, Interrogating the Etiology of Sporadic Alzheimer’s Disease Using Aging Rhesus Macaques: Cellular, Molecular, and Cortical Circuitry Perspectives. J Gerontol A Biol Sci Med Sci 78, 1523–1534 (2023).

89. N. Oikawa, N. Kimura, K. Yanagisawa, Alzheimer-type tau pathology in advanced aged nonhuman primate brains harboring substantial amyloid deposition. Brain Res 1315, 137–149 (2010).

90. C. D. Paspalas, B. C. Carlyle, S. Leslie, T. M. Preuss, J. L. Crimins, A. J. Huttner, C. H. van Dyck, D. L. Rosene, A. C. Nairn, A. F. T. Arnsten, The aged rhesus macaque manifests Braak stage III/IV Alzheimer’s-like pathology. Alzheimers Dement 14, 680–691 (2018).

91. A. F. T. Arnsten, D. Datta, S. Leslie, S.-T. Yang, M. Wang, A. C. Nairn, Alzheimer’s-like pathology in aging rhesus macaques: Unique opportunity to study the etiology and treatment of Alzheimer’s disease. Proceedings of the National Academy of Sciences 116, 26230–26238 (2019).

92. R. García-Pérez, J. M. Ramirez, A. Ripoll-Cladellas, R. Chazarra-Gil, W. Oliveros, O. Soldatkina, M. Bosio, P. J. Rognon, S. Capella-Gutierrez, M. Calvo, F. Reverter, R. Guigó, F. Aguet, P. G. Ferreira, K. G. Ardlie, M. Melé, The landscape of expression and alternative splicing variation across human traits. Cell Genom 3, 100244 (2023).

93. Y. Hou, X. Dan, M. Babbar, Y. Wei, S. G. Hasselbalch, D. L. Croteau, V. A. Bohr, Ageing as a risk factor for neurodegenerative disease. Nat. Rev. Neurol. 15, 565–581 (2019).

94. L. Hu, Y.-J. Huang, Y.-D. Wei, T. Li, W. Ke, G.-H. Chen, M.-X. Dong, Plasma metabolomics profiles indicate sex differences of lipid metabolism in patients with Parkinson’s disease. Sci Rep 14, 31262 (2024).

95. L. M. Collins, A. Toulouse, T. J. Connor, Y. M. Nolan, Contributions of central and systemic inflammation to the pathophysiology of Parkinson’s disease. Neuropharmacology 62, 2154–2168 (2012).

96. P. García-Sanz, J. M. F. Aerts, R. Moratalla, The Role of Cholesterol in α-Synuclein and Lewy Body Pathology in GBA1 Parkinson’s Disease. Movement Disorders 36, 1070–1085 (2021).

97. R. Musanti, E. Parati, E. Lamperti, G. Ghiselli, Decreased cholesterol biosynthesis in fibroblasts from patients with Parkinson disease. Biochem Med Metab Biol 49, 133–142 (1993).

98. L. Dai, J. Wang, L. Meng, X. Zhang, T. Xiao, M. Deng, G. Chen, J. Xiong, W. Ke, Z. Hong, L. Bu, Z. Zhang, The cholesterol 24-hydroxylase CYP46A1 promotes α-synuclein pathology in Parkinson’s disease. PLOS Biology 23, e3002974 (2025).

99. H. Xicoy, B. Wieringa, G. J. M. Martens, The Role of Lipids in Parkinson’s Disease. Cells 8 (2019).

100. L. Lim, V. Jackson-Lewis, L. C. Wong, G. H. Shui, A. X. H. Goh, S. Kesavapany, A. M. Jenner, M. Fivaz, S. Przedborski, M. R. Wenk, Lanosterol induces mitochondrial uncoupling and protects dopaminergic neurons from cell death in a model for Parkinson’s disease. Cell Death Differ 19, 416–427 (2012).

101. A. Widdig, L. Muniz, M. Minkner, Y. Barth, S. Bley, A. Ruiz-Lambides, O. Junge, R. Mundry, L. Kulik, Low incidence of inbreeding in a long-lived primate population isolated for 75 years. Behav. Ecol. Sociobiol. 71, 18 (2017).

102. C. Testard, L. J. N. Brent, J. Andersson, K. L. Chiou, J. E. Negron-Del Valle, A. R. DeCasien, A. Acevedo-Ithier, M. K. Stock, S. C. Antón, O. Gonzalez, C. S. Walker, S. Foxley, N. R. Compo, S. Bauman, A. V. Ruiz-Lambides, M. I. Martinez, J. H. P. Skene, J. E. Horvath, C. B. R. Unit, J. P. Higham, K. L. Miller, N. Snyder-Mackler, M. J. Montague, M. L. Platt, J. Sallet, Social connections predict brain structure in a multidimensional free-ranging primate society. Sci. Adv. 8, eabl5794 (2022).

103. A. R. DeCasien, K. L. Chiou, C. Testard, A. Mercer, J. E. Negrón-Del Valle, S. E. Bauman Surratt, O. González, M. K. Stock, A. V. Ruiz-Lambides, M. I. Martínez, S. C. Antón, C. S. Walker, J. Sallet, M. A. Wilson, L. J. N. Brent, M. J. Montague, C. C. Sherwood, M. L. Platt, J. P. Higham, N. Snyder-Mackler, Evolutionary and biomedical implications of sex differences in the primate brain transcriptome. Cell Genom. 4, 100589 (2024).

104. J. Cao, M. Spielmann, X. Qiu, X. Huang, D. M. Ibrahim, A. J. Hill, F. Zhang, S. Mundlos, L. Christiansen, F. J. Steemers, C. Trapnell, J. Shendure, The single-cell transcriptional landscape of mammalian organogenesis. Nature 566, 496–502 (2019).

105. A. Dobin, C. A. Davis, F. Schlesinger, J. Drenkow, C. Zaleski, S. Jha, P. Batut, M. Chaisson, T. R. Gingeras, STAR: ultrafast universal RNA-seq aligner. Bioinformatics 29, 15–21 (2013).

106. W. C. Warren, R. A. Harris, M. Haukness, I. T. Fiddes, S. C. Murali, J. Fernandes, P. C. Dishuck, J. M. Storer, M. Raveendran, L. W. Hillier, D. Porubsky, Y. Mao, D. Gordon, M. R. Vollger, A. P. Lewis, K. M. Munson, E. DeVogelaere, J. Armstrong, M. Diekhans, J. A. Walker, C. Tomlinson, T. A. Graves-Lindsay, M. Kremitzki, S. R. Salama, P. A. Audano, M. Escalona, N. W. Maurer, F. Antonacci, L. Mercuri, F. A. M. Maggiolini, C. R. Catacchio, J. G. Underwood, D. H. O’Connor, A. D. Sanders, J. O. Korbel, B. Ferguson, H. M. Kubisch, L. Picker, N. H. Kalin, D. Rosene, J. Levine, D. H. Abbott, S. B. Gray, M. M. Sanchez, Z. A. Kovacs-Balint, J. W. Kemnitz, S. M. Thomasy, J. A. Roberts, E. L. Kinnally, J. P. Capitanio, J. H. P. Skene, M. Platt, S. A. Cole, R. E. Green, M. Ventura, R. W. Wiseman, B. Paten, M. A. Batzer, J. Rogers, E. E. Eichler, Sequence diversity analyses of an improved rhesus macaque genome enhance its biomedical utility. Science 370 (2020).

107. I. Virshup, S. Rybakov, F. J. Theis, P. Angerer, F. Alexander Wolf, anndata: Annotated data, bioRxiv (2021)p. 2021.12.16.473007.

108. S. L. Wolock, R. Lopez, A. M. Klein, Scrublet: Computational Identification of Cell Doublets in Single-Cell Transcriptomic Data. Cell Syst 8, 281–291.e9 (2019).

109. F. A. Wolf, P. Angerer, F. J. Theis, SCANPY: large-scale single-cell gene expression data analysis. Genome Biol 19, 15 (2018).

110. H. Vuong, T. Luu, N. Nguyen, N. Tra, H. Nguyen, H. Nguyen, T. Truong, S. Pham, Benchmarking AlphaSC: A Leap in Single-Cell Data Processing, bioRxiv (2023)p. 2023.11.28.569108.

111. D. M. Jennewein, J. Lee, C. Kurtz, W. Dizon, I. Shaeffer, A. Chapman, A. Chiquete, J. Burks, A. Carlson, N. Mason, A. Kobawala, T. Jagadeesan, P. B. Basani, T. Battelle, R. Belshe, D. McCaffrey, M. Brazil, C. Inumella, K. Kuznia, J. Buzinski, D. D. Shah, S. M. Dudley, G. Speyer, J. Yalim, “The sol supercomputer at Arizona state university” in Practice and Experience in Advanced Research Computing (ACM, New York, NY, USA, 2023; 10.1145/3569951.3597573).

112. S. Mages, N. Moriel, I. Avraham-Davidi, E. Murray, J. Watter, F. Chen, O. Rozenblatt-Rosen, J. Klughammer, A. Regev, M. Nitzan, TACCO unifies annotation transfer and decomposition of cell identities for single-cell and spatial omics. Nature Biotechnology 41, 1465–1473 (2023).

113. Website. MapMyCells. https://portal.brain-map.org/atlases-and-data/bkp/mapmycells.

114. A. Subramanian, P. Tamayo, V. K. Mootha, S. Mukherjee, B. L. Ebert, M. A. Gillette, A. Paulovich, S. L. Pomeroy, T. R. Golub, E. S. Lander, J. P. Mesirov, Gene set enrichment analysis: A knowledge-based approach for interpreting genome-wide expression profiles. Proceedings of the National Academy of Sciences 102, 15545–15550 (2005).

115. A. Liberzon, C. Birger, H. Thorvaldsdóttir, M. Ghandi, J. P. Mesirov, P. Tamayo, The Molecular Signatures Database (MSigDB) hallmark gene set collection. Cell systems 1, 417 (2015).

116. B. Van de Sande, C. Flerin, K. Davie, M. De Waegeneer, G. Hulselmans, S. Aibar, R. Seurinck, W. Saelens, R. Cannoodt, Q. Rouchon, T. Verbeiren, D. De Maeyer, J. Reumers, Y. Saeys, S. Aerts, A scalable SCENIC workflow for single-cell gene regulatory network analysis. Nature Protocols 15, 2247–2276 (2020).

117. F. A. Wolf, F. K. Hamey, M. Plass, J. Solana, J. S. Dahlin, B. Göttgens, N. Rajewsky, L. Simon, F. J. Theis, PAGA: graph abstraction reconciles clustering with trajectory inference through a topology preserving map of single cells. Genome Biology 20, 1–9 (2019).

118. G. Yu, L.-G. Wang, Y. Han, Q.-Y. He, clusterProfiler: an R Package for Comparing Biological Themes Among Gene Clusters. doi: 10.1089/omi.2011.0118 (2012).

119. SynGO: An Evidence-Based, Expert-Curated Knowledge Base for the Synapse. Neuron 103, 217–234.e4 (2019).

120. M. Kanehisa, M. Furumichi, Y. Sato, Y. Matsuura, M. Ishiguro-Watanabe, KEGG: biological systems database as a model of the real world. Nucleic Acids Res 53, D672–D677 (2024).

121. Introducing Claude 4. https://www.anthropic.com/news/claude-4.

122. S. Alldritt, J. S. B. Ramirez, R. V. de Wael, R. Bethlehem, J. Seidlitz, Z. Wang, K. Nenning, N. B. Esper, J. Smallwood, A. R. Franco, K. Byeon, A. Alexander-Bloch, D. G. Amaral, C. Amiez, F. Balezeau, M. G. Baxter, G. Becker, J. Bennett, O. Berkner, E. L. A. Blezer, A. M. Brambrink, T. Brochier, B. Butler, L. J. Campos, E. Canet-Soulas, L. Chalet, A. Chen, J. Cléry, C. Constantinidis, D. J. Cook, S. Dehaene, L. Dorfschmidt, C. M. Drzewiecki, J. W. Erdman, S. Everling, A. Falchier, L. Fleysher, A. Fox, W. Freiwald, M. Froesel, S. Froudist-Walsh, J. Fudge, T. Funck, M. Gacoin, D. J. Gale, J. Gallivan, C. M. Garin, T. D. Griffiths, C. Guedj, F. Hadj-Bouziane, S. B. Hamed, N. Harel, R. Hartig, B. Hiba, B. R. Howell, B. Jarraya, B. Jung, N. Kalin, J. Karpf, S. Kastner, C. Klink, Z. A. Kovacs-Balint, C. Kroenke, M. J. Kuchan, S. C. Kwok, K. N. Lala, D. A. Leopold, G. Li, P. Lindenfors, G. Linn, R. B. Mars, K. Masiello, R. S. Menon, A. Messinger, M. Meunier, K. Mok, J. H. Morrison, J. Nacef, J. Nagy, V. Neudecker, M. Neuringer, M. P. Noonan, M. Ortiz-Rios, J. F. Perez-Zoghbi, C. I. Petkov, M. Pinsk, C. Poirier, E. Procyk, R. Rajimehr, S. M. Reader, D. A. Rudko, M. F. S. Rushworth, B. E. Russ, J. Sallet, M. M. Sanchez, M. C. Schmid, C. M. Schwiedrzik, J. A. Scott, J. Sein, K. K. Sharma, A. Shmuel, M. Styner, E. L. Sullivan, A. Thiele, O. S. Todorov, D. Tsao, A. Tusche, R. Vlasova, Z. Wang, L. Wang, J. Wang, A. R. Weiss, C. R. E. Wilson, E. Yacoub, W. Zarco, Y. Zhou, J. Zhu, D. Margulies, D. Fair, C. Schroeder, M. Milham, T. Xu, Brain Charts for the Rhesus Macaque Lifespan, bioRxiv (2024). 10.1101/2024.08.28.610193.

123. S. Dash, B. Park, C. D. Kroenke, W. D. Rooney, H. F. Urbanski, S. G. Kohama, Brain volumetrics across the lifespan of the rhesus macaque. Neurobiol Aging 126, 34–43 (2023).

124. F. M. Krienen, M. Goldman, Q. Zhang, R. C H Del Rosario, M. Florio, R. Machold, A. Saunders, K. Levandowski, H. Zaniewski, B. Schuman, C. Wu, A. Lutservitz, C. D. Mullally, N. Reed, E. Bien, L. Bortolin, M. Fernandez-Otero, J. D. Lin, A. Wysoker, J. Nemesh, D. Kulp, M. Burns, V. Tkachev, R. Smith, C. A. Walsh, J. Dimidschstein, B. Rudy, L. S Kean, S. Berretta, G. Fishell, G. Feng, S. A. McCarroll, Innovations present in the primate interneuron repertoire. Nature 586, 262–269 (2020).

125. B. Emery, Q. Richard Lu, Transcriptional and Epigenetic Regulation of Oligodendrocyte Development and Myelination in the Central Nervous System. Cold Spring Harbor Perspectives in Biology 7, a020461 (2015).

126. L. E. Dewald, J. P. Rodriguez, J. M. Levine, The RE1 binding protein REST regulates oligodendrocyte differentiation. J Neurosci 31, 3470–3483 (2011).

